# Common schizophrenia heritability concentrates in an evolutionarily young, brain-regulatory subset of fine-mapped credible-set variants

**DOI:** 10.64898/2026.07.30.741688

**Authors:** Yusuf Çiçek, Ali Tarık Altunç, Halil Aziz Velioğlu, Ömer Faruk Demirel

**Affiliations:** Department of Psychiatry, Cerrahpaşa Faculty of Medicine, Istanbul University-Cerrahpaşa, Istanbul, Türkiye; Department of Psychiatry, The Council of Forensic Medicine, Istanbul, Türkiye; Department of Physiology, Brain and Cognition Research Center, BEYKOG, Istanbul Medipol University, Istanbul, Türkiye

## Abstract

Common variation explains a substantial fraction of schizophrenia heritability, yet why these risk alleles persist remains unresolved. Of 20,766 fine-mapped PGC3 schizophrenia credible-set variants, the 4,918 with an allele-age estimate, brain-versus-blood regulatory specificity and a haplotype-based selection signal were clustered into three age-ordered subsets (Young, Mid and Old), all predating the out-of-Africa dispersal (cluster medians ≈113–508 kyr). The Young, brain-regulatory subset concentrated schizophrenia common-variant heritability after accounting for genome-wide allele age, frequency, linkage-disequilibrium and selection architecture (conditional coefficient Z = +3.05). The concentration generalized to East Asian schizophrenia (Z = +2.83), was balanced across sexes, and tracked genetic correlation across psychiatric disorders, including bipolar disorder, but not height or body-mass index. Their persistence at common frequency is most consistent with purifying selection and mutation-selection balance on these ancient, predominantly non-coding, brain-regulatory variants rather than recent adaptation.

## Introduction

Schizophrenia is a heritable psychiatric disorder with onset in late adolescence or early adulthood. Its lifetime prevalence is approximately 0.5–1%^1^, and twin studies estimate the heritability at 73–90%^2^. Common genetic variation accounts for a substantial fraction of this heritability^4,5^. The third Psychiatric Genomics Consortium (PGC) schizophreniagenome-wide association study (GWAS) identified 287 distinct genomic loci and provided fine-mapped credible-set posterior inclusion probabilities for 20,766 candidate variants^6^, building on a series of progressively larger consortium analyses^3,7^.

Why these common risk alleles persist despite the fecundity disadvantage carried by patients has been termed the evolutionary paradox of schizophrenia. Candidate mechanisms include heterozygote advantage^8^, antagonistic pleiotropy^9^ with cognitive or creative traits^10,11^, hitchhiking, mutation-selection balance^8^, and non-antagonistic pleiotropy^12^; these are not mutually exclusive, and the relative contribution of each remains an empirical question. Prior work has shown that common schizophrenia alleles concentrate in mutation-intolerant genes and in regions under strong background selection^13^, are enriched in genomic regions carrying signatures of recent positive selection along the human lineage^14^, include individual loci with directly demonstrated post-glacial selective sweeps^15^, and that derived alleles arisen during human evolution may confer protection rather than risk^12^. Brain transcriptomic profiling further indicates that psychiatric risk loci converge on shared neurodevelopmental gene-expression programs across schizophrenia, bipolar disorder and autism^16,17^, with partial overlap and dissociable subphenotypes between schizophrenia and bipolar disorder^18^. These results have operated at the level of genes or regions, leaving variant-resolution evolutionary inference unaddressed.

Two technical advances now enable a complementary variant-resolution analysis. The Atlas of Variant Age^19^ provides coalescent-based allele-age estimates from population-scale sequencing, supporting per-variant temporal inference; complementary genealogy-based frameworks^20,21^, including a unified tree sequence that incorporates ancient genomes, provide an independent cross-check at greater time depth. Ancient-DNA-derived selection coefficients^22^, building on earlier aDNA scans^23^, place recent directional selection on a directly observed ≈10,000-year West Eurasian time series, complementing modern haplotype-based scans^24^ and singleton-density signals^25^; these methods detect recent selection signatures independently of an allele’s coalescent age.

These signals can be combined with a third axis: the brain-versus-blood specificity of each variant’s regulatory effect. Prior between-population cis-eQTL adaptation scans^26^ have operated at genome-wide resolution and have not tested variant-level age effects within disease-specific credible sets. We therefore asked whether PGC3 schizophrenia credible-set variants carry evolutionary signatures of allele age and selection that vary with their tissue and cell-type of regulatory effect. We identify three coalescent-age subsets, all of which predate the out-of-Africa dispersal, and show that an evolutionarily young, brain-regulatory subset concentrates schizophrenia common-variant heritability (conditional coefficient Z = +3.05)^27^, with replication across European and East Asian populations and both sex strata, broad regulatory engagement across eight major brain cell types, and a cross-disorder signature that tracks schizophrenia genetic correlation while remaining absent in well-powered non-brain controls. Because these variants are ancient and predominantly non-coding, their persistence at common frequency is most consistent with purifying selection and mutation-selection balance on conserved brain-regulatory variation rather than recent or adaptive selection.

## Results

### Joint mixture-model clustering resolves three evolutionary variant subsets across deep, preout-of-Africa coalescent time

Three evolutionary variant subsets emerged from joint mixture-model clustering of PGC3 schizophrenia credible-set variants on log allele age × brain–blood regulatory specificity × |iHS| (integrated haplotype score) (Methods; Table 1, Fig. 1): cluster C0 (Young; *n* = 1,745; median ≈113 kyr; low–moderate |iHS|), cluster C1 (Mid; *n* = 317; median ≈358 kyr; elevated |iHS|), and cluster C2 (Old; *n* = 2,856; median ≈508 kyr; low |iHS|). These three subsets comprise 4,918 variants in total. Three clustering-specific metrics (Integrated Completed Likelihood, silhouette, Davies–Bouldin) independently selected *k* = 3, with the principal Young-cluster signal robust across *k* ∈ {2, 3, 4} (Supplementary Tables 5f, 16); the fit with 17q21.31 included produced an even more strongly preferred three-component solution (ΔBIC_k=3 vs. k=1_ < −15,000), indicating that 17q21.31 contributes mass to but does not exclusively account for the multimodality.

**Fig. 1.**
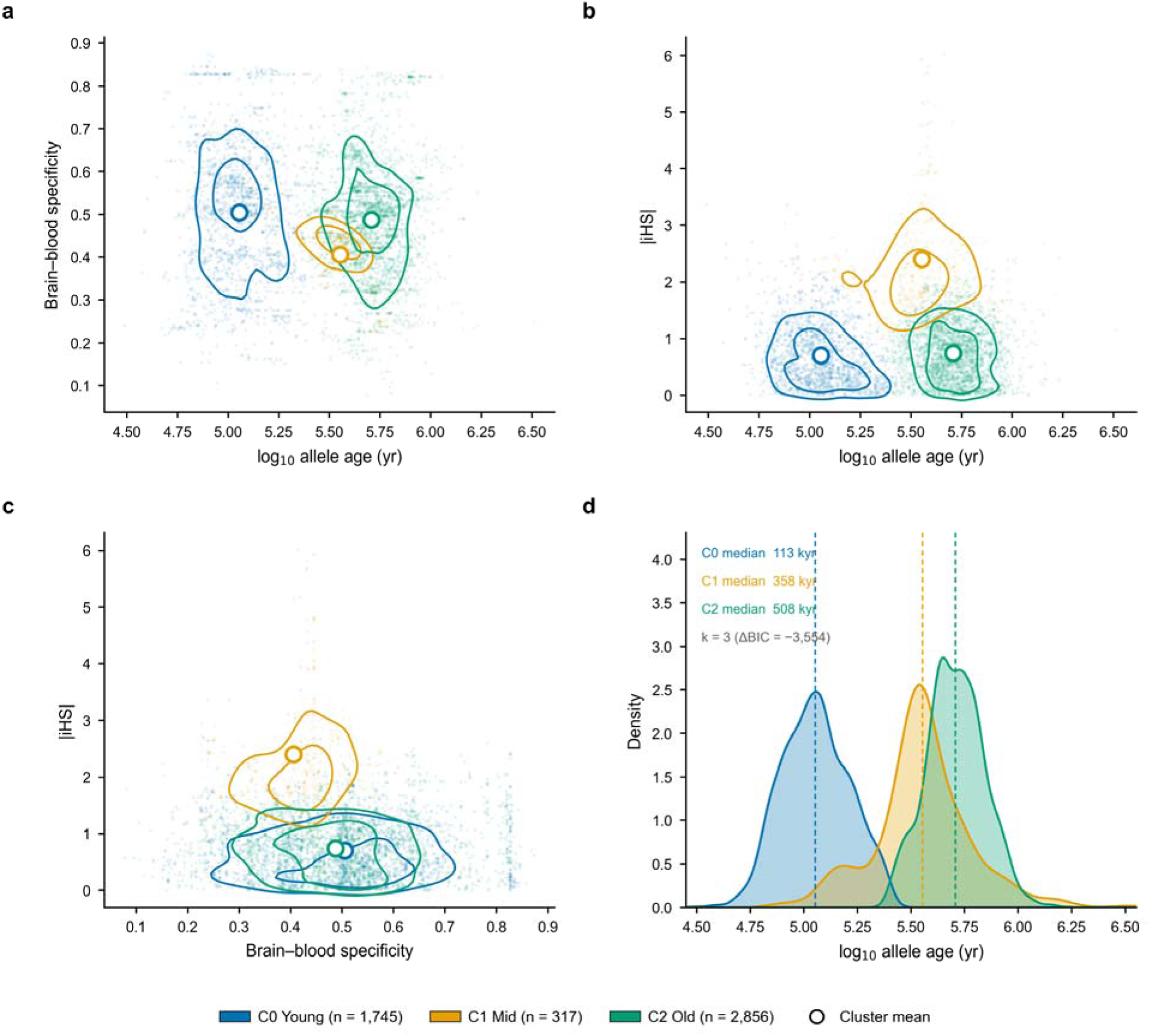
Three evolutionary variant subsets of fine-mapped PGC3 schizophrenia credible-set variants. Pairwise scatter projections of the joint feature space of log allele age, brain–blood regulatory specificity and Voight-2006-correct |iHS|, with PGC3 variants outside the 17q21.31 *MAPT* inversion locus coloured by GMM cluster assignment (C0 = Young, *n* = 1,745; C1 = Mid, *n* = 317; C2 = Old, *n* = 2,856). ΔBIC (*k* = 3 vs. *k* = 1) = −3,554 supports the multimodal structure (full *k*-selection rationale in Methods; Supplementary Table 5). (a) Brain–blood specificity versus log□□ allele age, (b) |iHS| versus log allele age and (c) |iHS| versus brain–blood specificity, each showing per-cluster two-dimensional kernel-density contours over the variant points with empirical cluster means overlaid (white circles); (d) per-cluster marginal kernel-density distributions of log allele age, with cluster medians marked (dashed lines; C0 ≈ 113 kyr, C1 ≈ 358 kyr, C2 ≈ 508 kyr). Allele ages are Atlas-of-Variant-Age coalescent estimates, reported in years after conversion from generations at 28.1 yr generation□¹ (Methods).

**Table 1.**
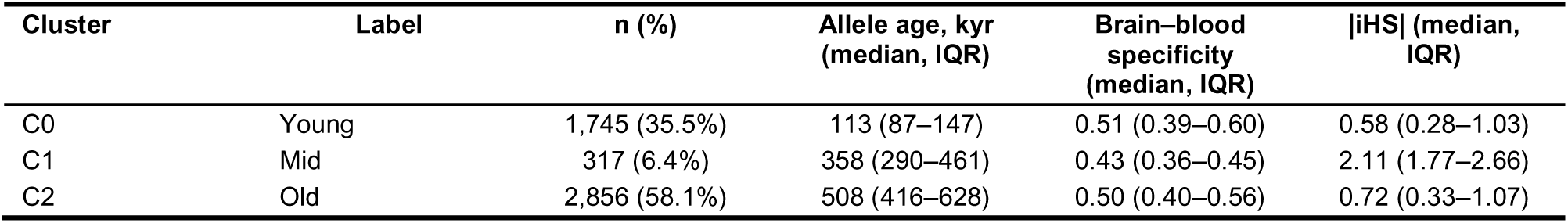
Characteristics of the three variant clusters. Three-component Gaussian mixture model on log₁₀ allele age × brain–blood regulatory specificity × Voight-corrected |iHS| for the 4,918 non-MAPT fine-mapped PGC3 schizophrenia credible-set variants. Allele ages are Atlas-of-Variant-Age coalescent estimates converted from generations to years at 28.1 yr generation⁻¹. IQR, interquartile range.

| Cluster | Label | n (%) | Allele age, kyr<br>(median, IQR) | Brain–blood<br>specificity<br>(median, IQR) | \|iHS\| (median,<br>IQR) |
| --- | --- | --- | --- | --- | --- |
| C0 | Young | 1,745 (35.5%) | 113 (87–147) | 0.51 (0.39–0.60) | 0.58 (0.28–1.03) |
| C1 | Mid | 317 (6.4%) | 358 (290–461) | 0.43 (0.36–0.45) | 2.11 (1.77–2.66) |
| C2 | Old | 2,856 (58.1%) | 508 (416–628) | 0.50 (0.40–0.56) | 0.72 (0.33–1.07) |

All three subsets are evolutionarily ancient. Their median ages are ≈113 kyr (C0), ≈358 kyr (C1) and ≈508 kyr (C2), all predating the out-of-Africa dispersal (≈65 kyr), with the oldest cluster reaching ≈1.4 Myr (Methods, Supplementary Table 21). Young, Mid and Old therefore label the youngest, intermediate and oldest strata of a single, deeply ancestral variant set. The same depth appears in an independent genealogy: in the Wohns et al.^21^ unified tree sequence, 96.9% of the credible-set variants are present in African lineages, and the three subsets retain their C0 < C1 < C2 age ordering (Fig. 5). These subsets form the substrate for the heritability, cross-ancestry, sex-stratified, cross-disorder and cell-type analyses that follow.

### The Young cluster concentrates schizophrenia common-variant heritability

The Young cluster (C0) concentrated PGC3 EUR schizophrenia common-variant heritability conditional on the 97-annotation baseline-LD v2.2 model^27^ (conditional coefficient Z = +3.05; 47.4-fold, s.e. = 14.9, P = 2.0 × 10 ³; Methods). The Mid cluster (C1) was also significantly enriched (Z = +3.32; 13.0-fold; P = 2.1 × 10□³), while the Old cluster (C2) was not (15.6-fold; P = 0.26; Table 2, Fig. 2a). Both C0 and C1 exceeded the focal-annotation significance threshold (Z = 2.81; Methods). Masking the 24 Price 2008 long-range LD regions^28^ shifted enrichment estimates by less than two-fold absolute for all comparisons (C0 47.4 → 47.9-fold, Z = +3.01; Table 2, Fig. 2c), and including the 17q21.31 *MAPT* inversion in the cluster annotation gave a near-identical principal C0 enrichment (47.8-fold, P = 1.6 × 10 ³, Z = +3.12; Supplementary Table 18), confirming the *MAPT*-exclusion choice does not drive the signal. Because the baseline-LD v2.2 model itself includes a MAF-adjusted predicted-allele-age annotation and ancient-sequence-age annotations for human promoters and enhancers (Methods), the C0 coefficient was estimated conditional on the genome-wide relationship between allele age and heritability, and therefore does not restate it.

**Fig. 2.**
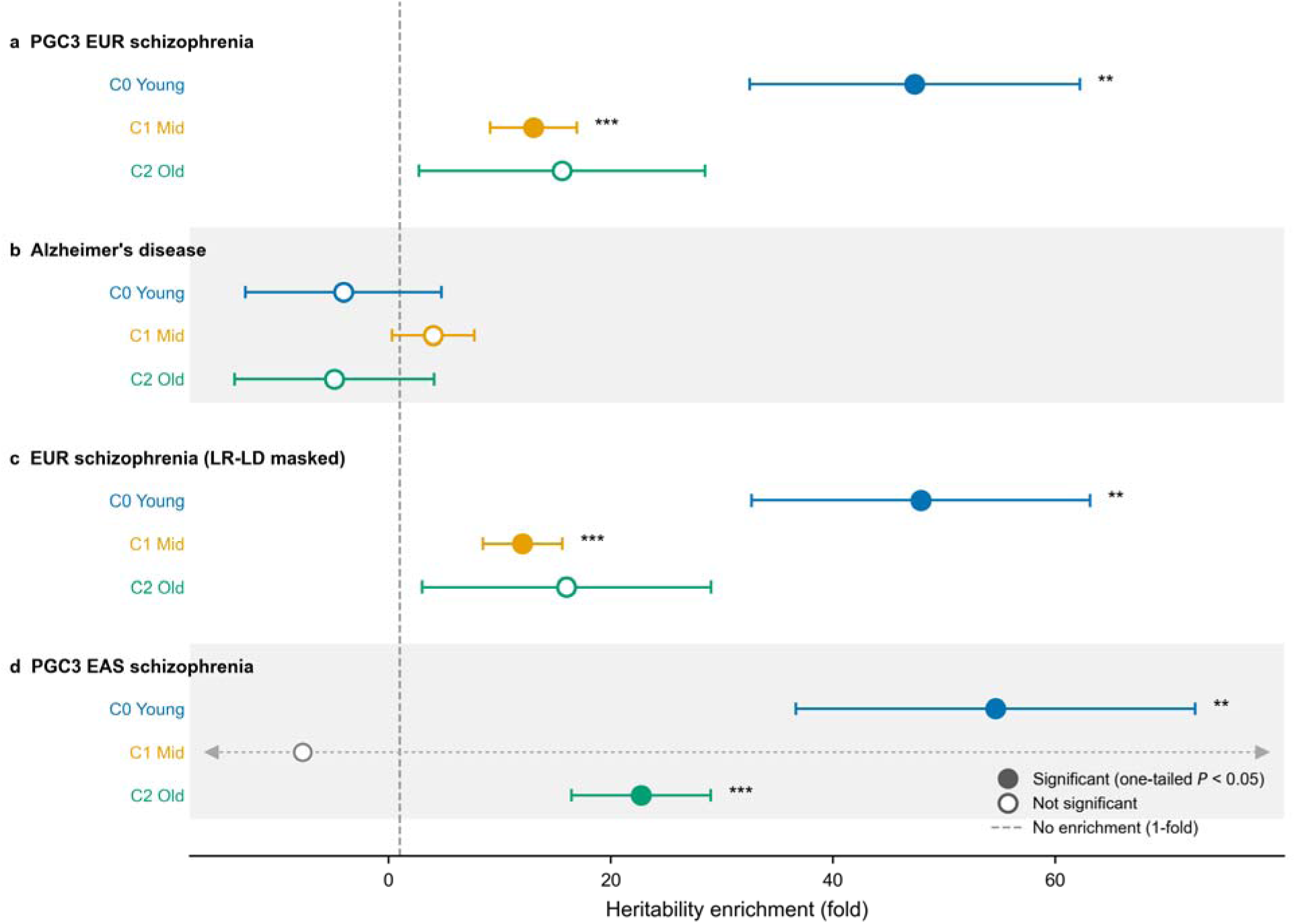
Cluster-level partitioned heritability across SCZ EUR, Alzheimer’s disease, Price 2008 LR-LD-masked SCZ EUR and SCZ EAS, conditional on baseline-LD v2.2^27^. Forest plot of cluster heritability enrichment (point = fold enrichment; whiskers = ±1 standard error) for the three variant clusters (C0 Young, C1 Mid, C2 Old) within each GWAS sub-group; the dashed line at 1-fold indicates no enrichment. Filled markers denote a significant conditional coefficient (one-tailed *P* < 0.05); open markers, non-significant. Asterisks indicate the conditional-coefficient *Z*-score significance (* *P* ≤ 0.05; ** *P* ≤ 0.01; *** *P* ≤ 0.001). (a) PGC3 EUR SCZ^6^: C0 47.4-fold, C1 13.0-fold, C2 15.6-fold; both C0 and C1 conditional coefficient *Z* > 2.81 (baseline-LD v2.2 single-test Bonferroni). (b) Wightman 2021 AD^29^: all clusters non-significant; the C0 estimate (−4.1-fold, s.e. = 8.8) has a standard error exceeding the point estimate in absolute magnitude and is therefore underpowered to reject shared SCZ–AD architecture rather than evidence for SCZ specificity. (c) Price 2008 LR-LD-masked^28^ SCZ EUR: enrichments shifted by less than 2-fold absolute relative to (a) (C0 47.4 → 47.9-fold). (d) PGC3 EAS SCZ^6^: C0 54.6-fold point-estimate replicates EUR (within ≈0.5 s.e., *Z* = +2.83); C2 22.7-fold emerges with Bonferroni-significant support (*Z* = +3.59); the C1 estimate is uninformative after propagation to the East Asian reference (s.e. ≈ 96, small mapped-SNP count) and is drawn as an open grey point with its standard error extending beyond the axis.

**Table 2.** Cluster-level partitioned heritability (stratified LD-score regression, baseline-LD v2.2). Enrichment (fold) and conditional coefficient z for each cluster within each GWAS, conditional on the baseline-LD v2.2 model. P is the LDSC block-jackknife enrichment test (H₀: enrichment = 1); the conditional coefficient z (focal-annotation significance threshold 2.81) is the primary significance criterion. Negative fold values are unconstrained S-LDSC ratio estimates with large standard errors (non-significant); the East Asian C1 estimate is uninformative (s.e. ≈ 96, small mapped-SNP count). Prop. SNPs, proportion of SNPs in the annotation. Cross-disorder rows show the Young cluster (C0) only; full C0/C1/C2 results for all disorders are in Supplementary Table 11.

| GWAS | Cluster | Prop. SNPs | Enrichment<br>(fold) | Enrichment<br>s.e. | Coefficient $z$ | P (enrichment) |
| --- | --- | --- | --- | --- | --- | --- |
| PGC3 EUR SCZ | C0 Young | 0.0003 | 47.4 | 14.9 | +3.05 | 2.0e-03 |
| PGC3 EUR SCZ | C1 Mid | 0.0003 | 13.0 | 3.9 | +3.32 | 2.1e-03 |
| PGC3 EUR SCZ | C2 Old | 0.0004 | 15.6 | 12.9 | +1.04 | 2.6e-01 |
| Alzheimer's disease | C0 Young | 0.0003 | -4.1 | 8.8 | -0.70 | 5.6e-01 |
| Alzheimer's disease | C1 Mid | 0.0003 | 4.0 | 3.7 | +1.13 | 3.8e-01 |
| Alzheimer's disease | C2 Old | 0.0004 | -4.9 | 9.0 | -0.83 | 4.8e-01 |
| EUR SCZ (LR-LD masked) | C0 Young | 0.0003 | 47.9 | 15.2 | +3.01 | 2.3e-03 |
| EUR SCZ (LR-LD masked) | C1 Mid | 0.0003 | 12.1 | 3.6 | +3.32 | 2.2e-03 |
| EUR SCZ (LR-LD masked) | C2 Old | 0.0003 | 16.0 | 13.0 | +1.06 | 2.5e-01 |
| PGC3 EAS SCZ | C0 Young | 0.0003 | 54.6 | 18.0 | +2.83 | 4.4e-03 |
| PGC3 EAS SCZ | C1 Mid | 0.0001 | -7.7 | 95.7 | -0.09 | 9.3e-01 |
| PGC3 EAS SCZ | C2 Old | 0.0004 | 22.7 | 6.3 | +3.59 | 3.2e-04 |
| PGC bipolar disorder | C0 Young | 0.0003 | 23.9 | 10.1 | +2.14 | 2.4e-02 |
| PGC major depression | C0 Young | 0.0003 | 12.1 | 6.5 | +1.55 | 8.9e-02 |
| PGC ADHD | C0 Young | 0.0003 | 12.8 | 17.1 | +0.62 | 4.9e-01 |
| iPSYCH-PGC autism | C0 Young | 0.0003 | 5.5 | 12.7 | +0.27 | 7.2e-01 |
| CDG3 F3 (neurodevelopmental) | C0 Young | 0.0003 | 5.2 | 8.8 | +0.39 | 6.4e-01 |
| CDG3 F4 (internalising) | C0 Young | 0.0003 | 14.2 | 7.7 | +1.58 | 8.5e-02 |

### Variant-level evolutionary architecture distinguishes schizophrenia from Alzheimer’s disease

In an Alzheimer’s disease GWAS^29^, the Young cluster (C0) carried a non-significant negative point estimate (−4.1-fold, s.e. = 8.8, *P* = 0.56), opposite in direction to the SCZ enrichment (47.4-fold, s.e. = 14.9); the Mid and Old clusters were similarly non-significant (C1: +4.0-fold, *P* = 0.38; C2: −4.9-fold, *P* = 0.48; Table 2, Fig. 2b; Methods). Although the AD GWAS *h*² (0.016) is approximately 23-fold smaller than the schizophrenia estimate and the AD-alone test is therefore underpowered to detect a cluster-level enrichment, the observed AD point estimates are quantitatively incompatible with a shared architecture as strong as that of schizophrenia: an AD C0 enrichment matching the SCZ value (47.4-fold) would lie more than five standard errors above the observed point estimate.

### Cross-disorder enrichment tracks schizophrenia genetic correlation (rg) in proportion to power

Across six psychiatric phenotypes, C0 enrichment scaled with genetic correlation to schizophrenia (Table 2, Fig. 2; Supplementary Table 11). The strongest signal came from PGC bipolar disorder^30^, where C0 was nominally enriched (Z = +2.14; 23.9-fold; P = 0.024) but did not survive correction for the six disorders tested (Bonferroni α = 8.3 × 10 ³). Major depression and the CDG3 internalising factor F4 were not significant (12.1-fold, P = 0.089; 14.2-fold, P = 0.085). We found no significant enrichment in ADHD, autism or the CDG3 neurodevelopmental factor F3 (all P > 0.4).

This gradient is consistent with a shared-architecture model. Because C0 is defined on PGC3 SCZ, any cross-disorder enrichment should scale with its genetic correlation to schizophrenia (Methods). Predicted enrichments were ≈31-fold for BD, ≈16-fold for MDD, ≈9-fold for ADHD and ≈10-fold for autism. The BD and MDD point estimates fell within approximately one standard error of these predictions (Supplementary Table 11); the ADHD, autism and F3 estimates were too imprecise to test individually (all P > 0.4) but none exceeded the genetic-correlation expectation. C0 therefore reflects psychiatric architecture in proportion to genetic correlation with schizophrenia.

To test whether the signal is specific to psychiatric phenotypes, we applied the same S-LDSC procedure to two non-brain traits, GIANT adult height and body-mass index^31^. Neither was enriched for C0 (height: 7.5-fold, *Z* = +0.62, *P* = 0.37; BMI: −3.7-fold, *Z* = −0.99, *P* = 0.44; Supplementary Table 13). The BMI estimate fell marginally below zero, which in S-LDSC indicates a negligible heritability contribution, not depletion (Methods §Partitioned LD score regression). C0 therefore lies within psychiatric, schizophrenia-correlated architecture, and outside both generic ancient-allele signal and non-brain polygenicity.

### The Young-cluster signal is portable across ancestries and corroborated in deep-time genealogies

The Young cluster (C0) enrichment replicated in PGC3 East Asian schizophrenia (conditional coefficient Z = +2.83; 54.6-fold, s.e. = 18.0, P = 4.4 × 10 ³; Methods, Table 2, Fig. 2d), consistent with the European estimate (Z = +3.05; 47.4-fold), so the principal C0 signal is portable across the two Eurasian populations. Because the clusters are defined on European allele ages, European |iHS| and largely European cis-eQTL annotations and propagated to the East Asian reference, this tests portability of the European-defined subsets, not independent ancestry-specific replication. The Mid cluster (C1) retained only ≈56 of its 317 European SNPs after propagation, leaving its East Asian estimate uninformative (Z = −0.09, P = 0.93), paralleling its lowest brain cell-type cis-eQTL coverage (Fig. 3, Supplementary Table 7). The Old cluster (C2) was significant in East Asian schizophrenia (Z = +3.59, P = 3.2 × 10□□; 22.7-fold) but not in European (Z = +1.04); with the present data this lineage difference is not separable from East Asian LD, allele frequency and power. The portability of the C0 signal is the central cross-ancestry finding.

**Fig. 3.**
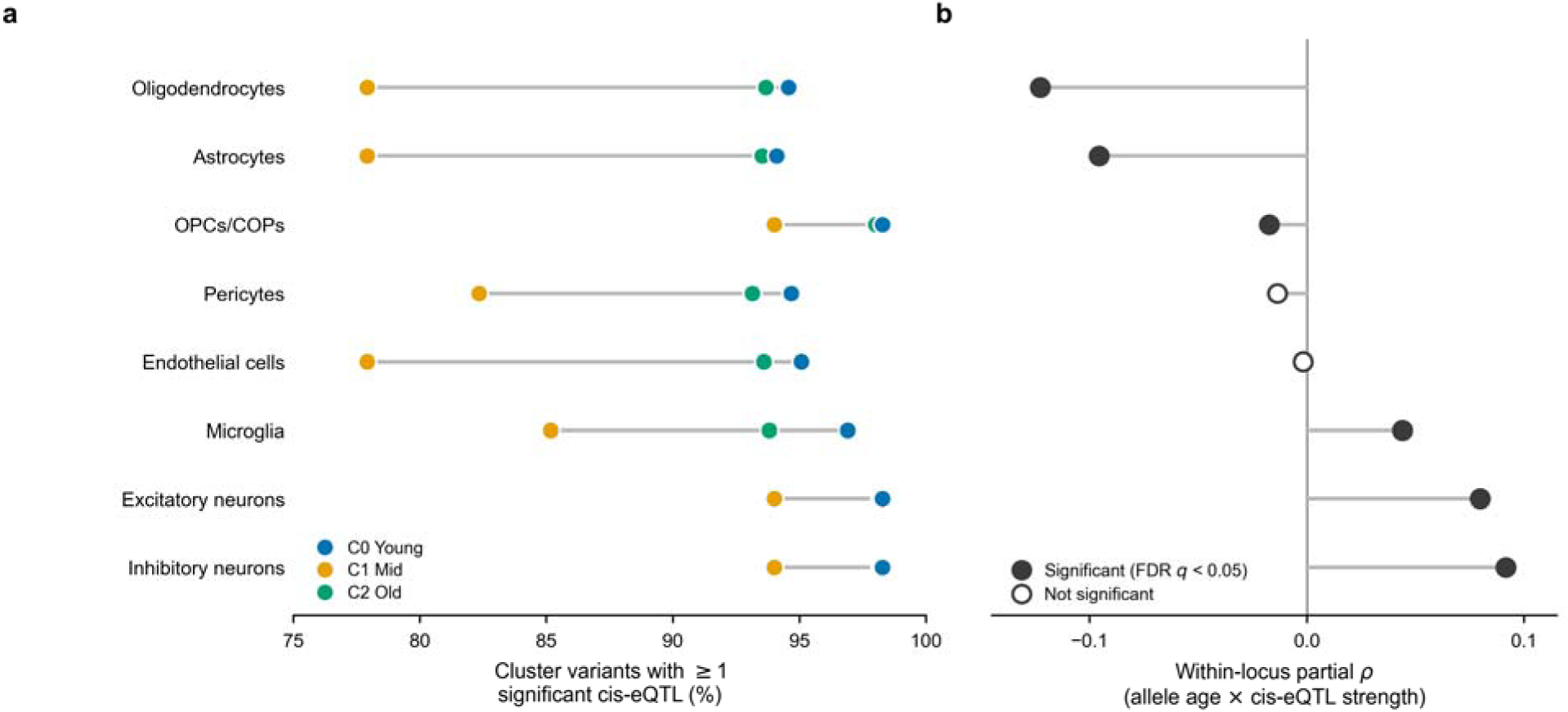
Cluster cell-type *cis*-eQTL coverage and within-locus age–effect dynamics across the eight Bryois 2022^34^ brain cell types. (a) Proportion of each cluster’s variants with at least one significant single-cell *cis*-eQTL in each cell type, as a dumbbell plot (C0 / C1 / C2 points joined by a grey connector), cell types ordered by the panel-b correlation. The Mid cluster (C1, *n* = 317) has the lowest coverage across all eight cell types, 4–17 percentage points below both C0 and C2; the Young cluster (C0) has the highest coverage in every cell type, with C0–C2 difference uniformly small (0.1–3.1 percentage points). Three-way cluster χ² *P*-values are uniformly highly significant (range 6 × 10 to 6 × 10□²□); microglia shows the largest Young–Old (C0–C2) coverage difference (3.1 percentage points; three-way χ² *P* = 1.4 × 10□¹□), a contrast dominated by depressed Mid-cluster (C1, 85.2%) engagement rather than by elevated Young-cluster microglia-specificity. (b) Within-locus allele-age × per-cell-type *cis*-eQTL-strength partial rank correlation (MAPT-excluded, MAF-rank residualised), as a diverging lollipop about ρ = 0; filled markers denote BH-FDR *q* < 0.05. The sign reverses across cell types: oligodendrocytes ρ = −0.123 and astrocytes ρ = −0.096 (older variants carry weaker eQTLs) versus microglia ρ = +0.044, excitatory neurons ρ = +0.080 and inhibitory neurons ρ = +0.092 (full per-cell-type ρ, *n*, *P* in Supplementary Table 7).

The credible-set variants are ancestrally shared as well as old. In the Wohns et al.^21^ unified tree sequence, which incorporates ancient and present-day genomes, 96.9% of the variants are present in African lineages, and 76% have African-lineage coalescent ages older than 200 kyr (Fig. 5a; Methods). The C0, C1 and C2 subsets keep their age ordering in this African-context dating, C0 youngest and C2 oldest, while all three remain predominantly older than 200 kyr (Fig. 5a), so the relative ordering holds on a second, deeper genealogy. In an independent African-ancestry schizophrenia GWAS (PGC3 African-American), per-variant association strength (mean χ²) is carried by the deeply shared variants and falls to 0.66, against a 1.23 baseline, for the small set with recent African-lineage ages (Fig. 5b, Supplementary Table 22), placing the association in the ancient, ancestrally shared component. These analyses date and weight the European-ascertained variants in African samples and do not constitute independent African fine-mapping.

### The Young-cluster enrichment is balanced across sexes

C0 enrichment was robust to sex stratification in both European and East Asian samples. Every stratum was significantly enriched (39.6–54.4-fold; P range 3.9 × 10□³ to 0.029; Methods, Supplementary Table 12). All four point estimates fell within approximately one standard error of the corresponding full-sample estimate.

We assessed sex-specificity with a formal heterogeneity statistic (Z_diff_; Methods). The test did not reject sex-symmetry in either ancestry (Z_diff_ = −0.64, P = 0.53 for EUR; Z_diff_ = +0.03, P = 0.98 for EAS).

The EUR female cohort has a 30% smaller effective sample size than the EUR male cohort, so this analysis is not equally powered across strata. We therefore frame the result as an absence of detectable sex-specific evolutionary architecture, not as positive evidence of strict sex-symmetry.

### Within-locus correlations reproduce the cluster-level pattern

We next asked whether the cluster-level pattern was reproduced at the within-locus level, using partial rank correlations among individual credible-set variants (Methods §Within-locus partial rank correlation).

Brain–blood regulatory specificity correlated negatively with log allele age (ρ = −0.093, *n* = 5,724 variants from the 84 credible sets with sufficient dual-compartment (brain and blood) cis-eQTL coverage, asymptotic *P* = 1.8 × 10 ¹²; Fig. 4a,d). The signal was sensitive to between-locus heterogeneity, however: a 1,000-iteration per-locus block-bootstrap returned a 95% CI of [−0.242, +0.021], which crossed zero (Supplementary Tables 2, 3). This correlation is therefore directionally supportive of the cluster-level finding rather than an independent statistical anchor.

**Fig. 4.**
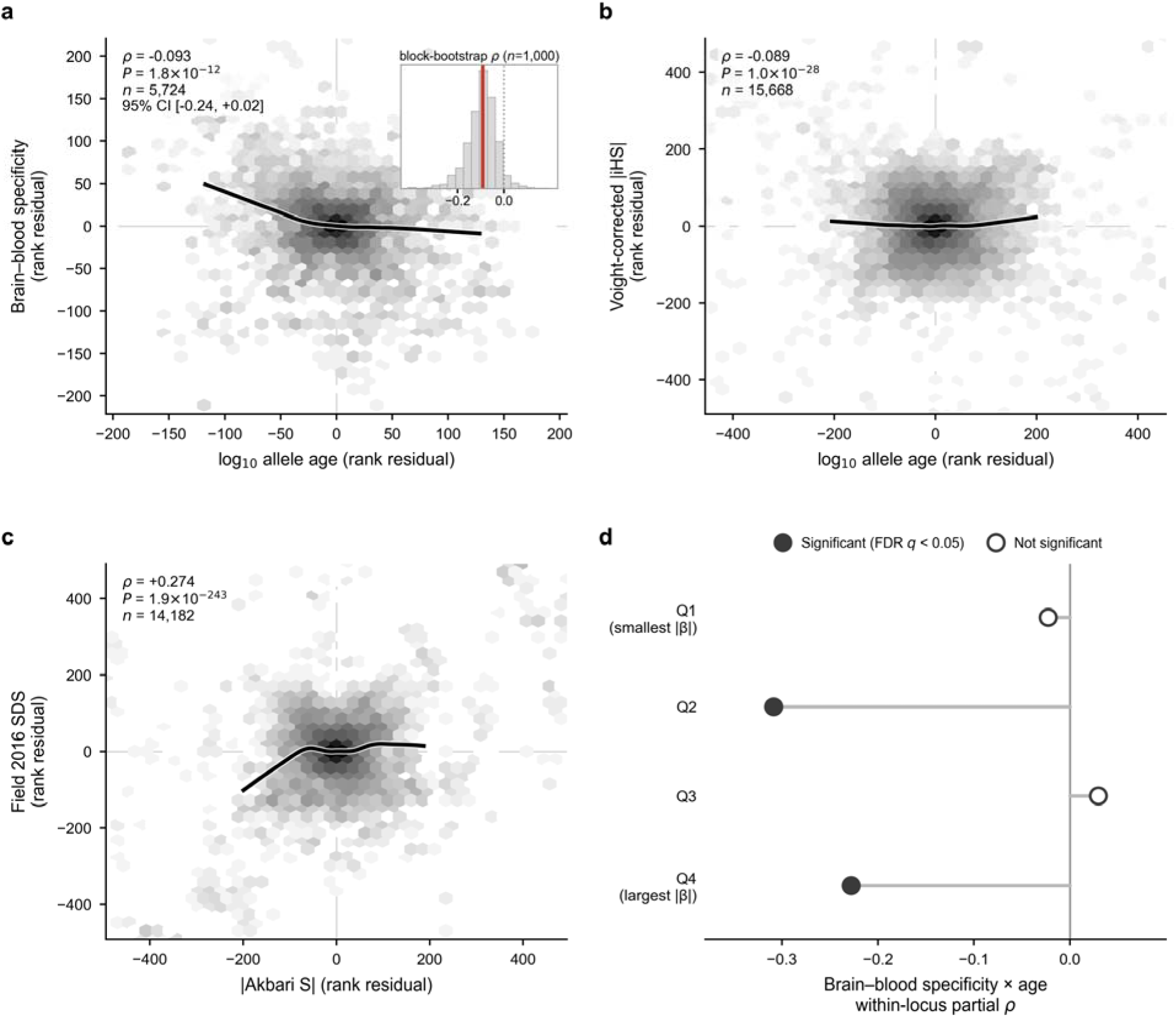
Within-locus partial rank correlations between regulatory specificity, selection metrics and allele age, pooled and stratified. (a) Brain–blood specificity × log□□ allele age, pooled within-locus rank residuals (no *MAPT*, MAF-rank residualised; ρ = −0.093, *P* = 1.8 × 10□¹², *n* = 5,724), shown as a two-dimensional density (hexagonal bins) with a LOWESS trend; the inset is the 1,000-iteration per-locus block-bootstrap distribution of ρ (observed value in red; 95% CI [−0.242, +0.021], which crosses zero). (b) Voight-correct |iHS| × log allele age (no *MAPT*, MAF-rank residualised; ρ = −0.089, *P* = 1.0 × 10□²□, *n* = 15,668). (c) Cross-method triangulation: |Akbari S| × Field 2016 SDS within-locus partial rank ρ = +0.274 (*P* = 1.9 × 10□²□³, *n* = 14,182). (d) Brain–blood specificity × allele-age within-locus partial ρ stratified by GWAS |β| quartile (Q1 smallest to Q4 largest); filled markers denote BH-FDR *q* < 0.05. The specificity–age signal concentrates in Q2 (ρ = −0.31) and Q4 (ρ = −0.23); Q1 and Q3 are non-significant.

**Fig. 5.**
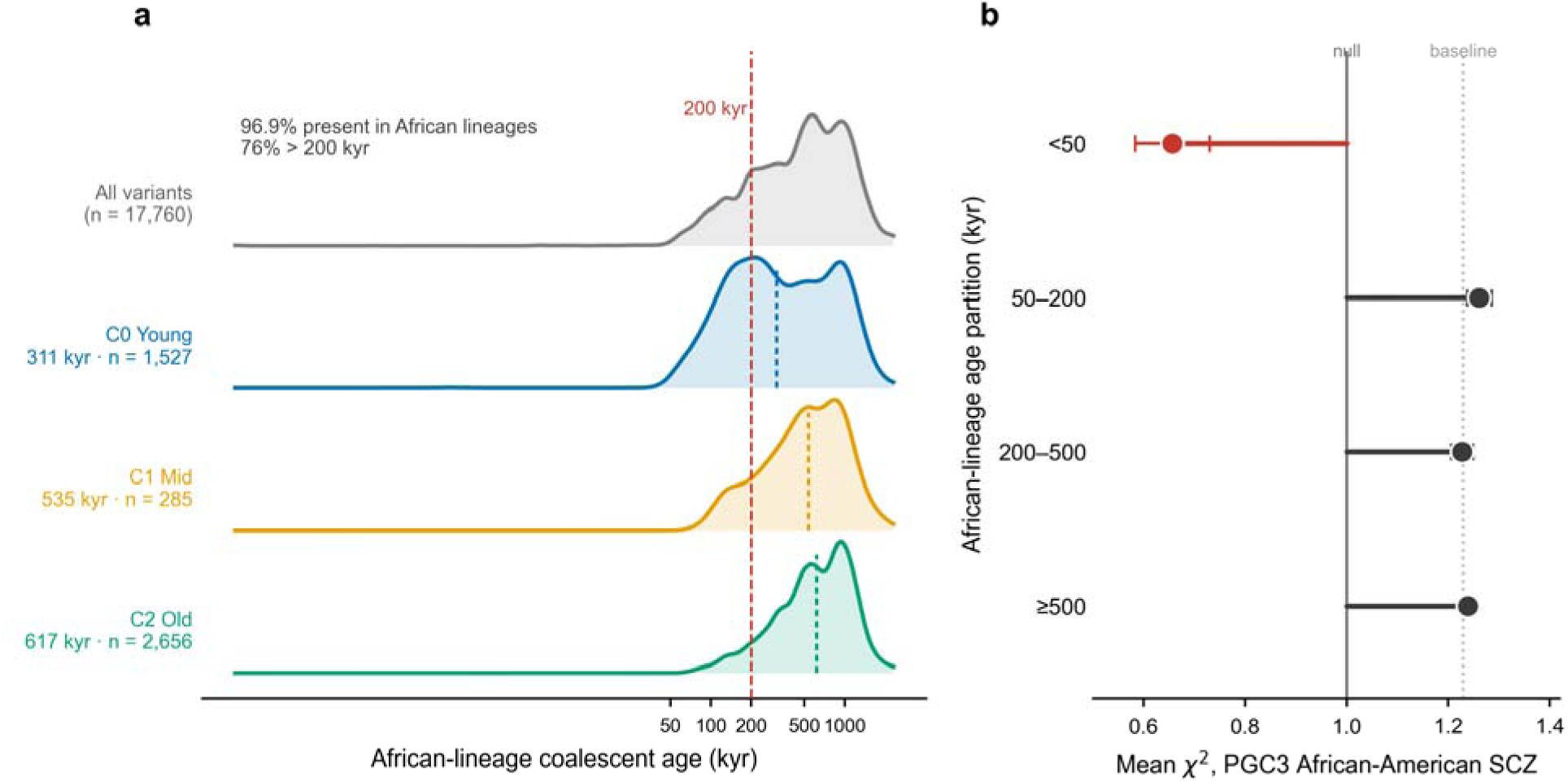
Deep-time and cross-ancestry corroboration of the variant subsets in African genealogies. (a) Distribution of African-lineage coalescent ages of the credible-set variants in the Wohns et al.^21^ unified tree sequence, shown as a ridgeline for all variants and by cluster (C0 Young, C1 Mid, C2 Old): 96.9% of variants are present in African lineages and 76% have ages older than 200 kyr (dashed line), and the three clusters retain their C0 < C1 < C2 ordering (African-lineage medians 311 / 535 / 617 kyr), all predominantly older than 200 kyr. (b) Per-variant association strength (mean χ² ± s.e.m.) in PGC3 African-American schizophrenia across African-lineage age partitions, anchored at the null (χ² = 1) with the credible-set baseline (χ² ≈ 1.23) marked; mean χ² is lowest (0.66) for the small set with recent (<50 kyr) African-lineage ages and sits at the baseline for the older, deeply shared partitions. Methods; Supplementary Table 22.

The |iHS| × log□□ allele age correlation was negative when 17q21.31 was excluded (ρ = −0.089, *n* = 15,668, *P* = 1.0 × 10□²□; Fig. 4b). This direction is what standard population-genetic theory predicts: ancient sweeps have decayed below the |iHS| detection threshold, leaving recent partial sweeps to dominate the genome-wide signal^32,33^. The sign reversed when 17q21.31 was included (ρ = +0.097, *P* = 1.0 × 10 ³). This reversal reflects the unusual structural-variant architecture of the *MAPT* inversion, which suppresses recombination and inflates the |iHS| signal independently of selection. We retain the 17q21.31-excluded analysis as the primary reading.

The |Akbari S| and Field 2016 SDS metrics correlated strongly (ρ = +0.274, *n* = 14,182, *P* = 1.9 × 10□²□³; Table 3 and Supplementary Table 4; Fig. 4c), indicating a coherent directional-selection signal from both ancient-DNA and modern-frequency frameworks. These metrics detect recent frequency change on standing variation of any coalescent age, indexing selection acting on ancient variants rather than a recent origin. Brain-specific *cis*-eQTLs in turn concentrated among variants with elevated |Akbari S| (ρ = +0.226, *P* = 3.6 × 10□□¹; Table 3), so brain regulatory specificity tracks the selection signal at the variant level.

**Table 3.**
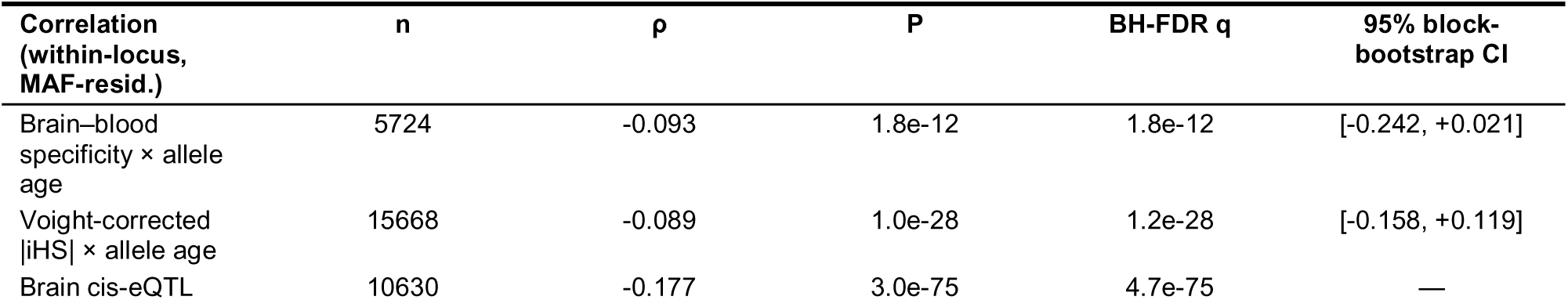
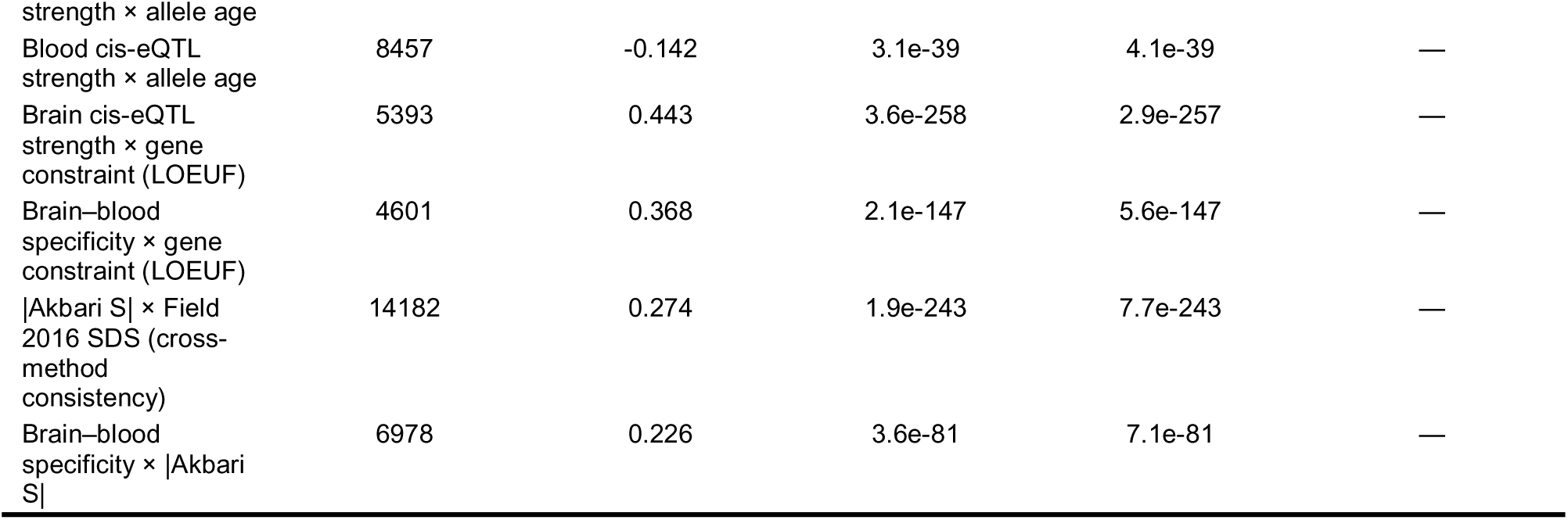
Primary within-locus partial-rank correlations. Within-credible-set rank residuals (credible sets with ≥5 variants), MAF-rank residualised, with the 17q21.31 MAPT inversion excluded. ρ, Spearman partial rank correlation; P, asymptotic; BH-FDR q across the eight correlations. The 95% confidence interval is from a 1,000-iteration per-locus block bootstrap (computed for the two headline allele-age correlations). LOEUF, gnomAD loss-of-function observed/expected upper-bound fraction; SDS, Field 2016 singleton density score.

### The Young cluster engages all eight brain cell types

C0 carried the highest *cis*-eQTL coverage in all eight brain cell types^34^ (Methods; Fig. 3, Supplementary Table 7). The C0–C2 differences were uniformly small (0.1–3.1 percentage points), indicating that the Young-cluster signal reflects broad regulatory engagement across cell types rather than concentration in any single one. C1 was consistently the low cluster, sitting 4– 17 percentage points below both C0 and C2.

The three-way cluster contrast was statistically significant across all eight cell types (χ² P range 6 × 10□□ to 6 × 10□²□); microglia showed the largest Young–Old (C0–C2) coverage difference (3.1 percentage points; χ² P = 1.4 × 10□¹□), reflecting reduced Mid-cluster (C1, 85.2%) engagement rather than elevated Young-cluster microglia-specificity.

Within-locus age × per-cell-type *cis*-eQTL strength correlations revealed a cell-type-specific temporal pattern. Oligodendrocytes and astrocytes showed the strongest negative correlations (ρ = −0.123 and ρ = −0.096, respectively), so older variants carry weaker eQTLs in these glial subtypes. Microglia, inhibitory neurons and excitatory neurons showed positive correlations in the opposite direction (ρ = +0.044 to +0.092; full per-cell-type ρ, *n*, *P* in Supplementary Table 7). This sign reversal points to cell-type-specific temporal dynamics in the regulatory architecture of these variants.

Pathway-level enrichment of cluster-specific gene lists showed little differentiation (Supplementary Table 8). Cluster differentiation was confined to brain region of strongest *cis*-eQTL,broad cell-type usage and gene-level constraint (Supplementary Table 10), rather than to distinct biological pathways.

## Discussion

Schizophrenia common-variant heritability concentrates in an evolutionarily young, brain-regulatory subset of fine-mapped PGC3 credible-set variants^6^. Joint clustering on allele age, regulatory specificity and |iHS| resolves three evolutionary subsets spanning a deep, pre-out-of-Africa coalescent-age range; the Young cluster (C0) carries this heritability concentration conditional on the full baseline-LD model^27^. The signal replicated in PGC3 East Asian schizophrenia^6^, across all four sex × ancestry strata, and across a seven-axis robustness battery designed for the small-annotation regime^35^, including long-range LD region exclusion^28^. In an Alzheimer’s disease GWAS^29^, the C0 estimate had the opposite sign to the SCZ signal and is inconsistent with a shared architecture as strong as that of schizophrenia; the substantially smaller AD heritability nevertheless leaves the analysis underpowered to rule out subtler shared components. The C0 annotation extended to bipolar disorder, major depression and an internalising factor in proportion to schizophrenia genetic correlation, but not to non-psychiatric anthropometric traits.

The cross-ancestry behaviour of the three clusters constrains their evolutionary interpretation. C0 retains its enrichment magnitude across both Eurasian populations, whereas C1 is enriched only in European schizophrenia and C2 only in East Asian schizophrenia. Because the partition is defined on European allele ages, |iHS| and *cis*-eQTL annotations and then propagated to East Asia, this pattern indicates cross-ancestry portability of the principal C0 signal together with population differences in the detectability of the smaller subsets; it does not require three distinct evolutionary regimes. All three subsets predate the out-of-Africa dispersal (cluster medians ≈113–508 kyr; concordant with an independent genealogy^21^ at ≈311–617 kyr, where the variants are predominantly present in African lineages), so they are ancient and ancestrally shared rather than recently arisen. Their persistence at common frequency is therefore better explained by purifying selection and mutation-selection balance^36^ acting on weakly deleterious, deeply conserved brain-regulatory variation than by recent positive or soft-sweep adaptation. Consistent with this, derived alleles at selected schizophrenia loci preferentially confer protection rather than risk^12^, with the implicated genes mapping to non-brain tissues, supporting persistence as a by-product of selection on conserved regulatory function rather than a cognition-versus-disease trade-off.

Beyond the evolutionary-regime question, cross-disorder testing places C0 within psychiatric-spectrum genetic architecture, scaled by genetic correlation with schizophrenia. Bipolar disorder was nominally enriched but did not survive six-test Bonferroni correction, major depression and the internalising factor F4 were not significant, and ADHD, autism and the neurodevelopmental factor F3 did not exceed the genetic-correlation expectation. This pattern rejects strict schizophrenia-specificity and universal neurodevelopmental extension; smaller signals in ADHD or autism cannot, however, be ruled out given the limited power in those datasets. Neither non-psychiatric anthropometric trait was enriched, confirming that the signal does not extend beyond psychiatric architecture. Sex-stratified replication detected significant C0 enrichment in all four sex × ancestry strata, and formal heterogeneity tests did not reject sex-symmetry. The EUR female cohort is, however, 30% smaller than the EUR male cohort, limiting power to detect modest sex-specific effects. Subject to that caveat, the genetic architecture appears sex-shared even where the downstream clinical phenotype, including the fertility deficit, is sex-biased^39,40^.

At the cell-type level, C0 showed elevated *cis*-eQTL coverage across all eight brain cell types^34^, indicating broad regulatory engagement rather than concentration in any single cell type. The cell-type contrast was driven by reduced Mid-cluster engagement rather than by Young-cluster microglia-specificity. This pattern distinguishes the variant-level evolutionary architecture of C0 from the microglia-centred gene-level signature of late-onset neurodegeneration^41^ and from the synaptic-pruning literature centred on the *C4* locus^42,43,44,45,46^. Within-locus age × eQTL strength correlations showed a cell-type-specific sign reversal, with negative correlations in oligodendrocytes and astrocytes and positive correlations in microglia and neurons. This pattern is compatible with the hypothesis that glial regulatory effects of schizophrenia credible-set variants map to the relatively younger end of the uniformly pre-out-of-Africa age range relative to those operating through microglia and neurons, in line with evidence that human-lineage cortical evolution has elaborated glial as well as neuronal regulatory layers^47,48,49^. This interpretation requires testing with cell-type-resolved comparative-genomic data.

Several limitations qualify these results. First, the three-dimensional Gaussian mixture is fit on the subset of non-*MAPT* credible-set variants with all three features available (4,918 of 20,766), biasing the fitted set toward variants with dual-compartment regulatory annotation; the two-dimensional feature-space sensitivity recovers broader coverage but dissolves the heritability concentration. Second, the partitioned-heritability analysis was not extended to admixed PGC3 cohorts, because standard LDSC underestimates single-nucleotide polymorphism (SNP) heritability in admixed populations with out-of-sample reference panels^50^; replication in additional non-European ancestries awaits future PGC4 releases. Third, the cross-ancestry analysis propagates European-defined cluster labels to the East Asian reference, with no East Asian fine-mapping, allele-age or selection data, so it tests portability of the European-defined subsets rather than independent replication; likewise, the African-genealogy analysis dates and weights the European-ascertained variants in African samples and is not independent African fine-mapping. Fourth, the cross-disorder panel omits the PGC cross-disorder meta-analysis^51^ because of sample overlap and instead uses CDG3 Genomic-SEM factors^52^. Fifth, high-PIP causal candidates account for only a small fraction of credible-set variants^53,54,55^, so findings are credible-set-level signatures, not per-causal-variant claims. Sixth, cluster ages are European-coalescent Atlas estimates (reported in generations, converted at 28.1 yr generation ¹; ref. 21) cross-checked against an independent genealogy^21^, and are deep, pre-out-of-Africa order-of-magnitude estimates. Seventh, we did not perform a neutral-demographic simulation, and a formal simulation-based null is left to future work. Notwithstanding these caveats, the results identify an evolutionarily young, brain-regulatory subset that concentrates schizophrenia common-variant heritability, generalizes to East Asian schizophrenia and across both sex strata, and captures shared psychiatric architecture in proportion to schizophrenia genetic correlation.

## Methods

### Variant set and ascertainment

We obtained 20,766 fine-mapped credible-set variants from the PGC3 schizophrenia GWAS (their Supplementary Table S11d). Trubetskoy and colleagues generated these credible sets using FINEMAP^53^ at each genome-wide significant locus, retaining variants whose cumulative posterior inclusion probability reached 95% within the locus. Of the 287 distinct loci, five chromosome X loci were excluded in line with standard GWAS practice^84^ (see Supplementary Methods §chrX exclusion), leaving 20,637 autosomal credible-set variants distributed across 282 autosomal loci (annotation coverage for all variants is summarised in Supplementary Table 1). Of these, 250 loci met the minimum within-locus variant count required for the partial rank correlation framework (≥ 5 variants). The 17q21.31 *MAPT* inversion locus^56^, represented by a single credible set (CS_224) of 1,742 variants, was excluded from all primary analyses (rationale and held-out reintegration sensitivity in Supplementary Methods §*MAPT*-excluded primary substrate), leaving 249 non-*MAPT* credible sets for within-locus analyses.

### Allele age coverage and EUR-coalescent reference frame

Atlas of Variant Age estimates were available for 20,565 of the 20,637 autosomal credible-set variants (99.7%). The Atlas reports coalescent ages in generations; converted at 28.1 yr generation ¹, the distribution spans a broad, deep range (minimum ≈7.6 kyr, maximum ≈3.25 Myr, median ≈382 kyr; 99.0% younger than ≈954 kyr; 99.9% younger than ≈1.38 Myr). The Atlas GEVA (Genealogical Estimation of Variant Age) estimator^19^ infers the time to the most recent common ancestor (TMRCA) of present-day European carriers; the median coalescent age of ≈382 kyr substantially predates the out-of-Africa dispersal (≈65 kyr), placing the bulk of the credible-set substrate in the ancestral, pre-out-of-Africa human lineage. The Wohns et al.^21^ unified tree-sequence framework provides an independent cross-check at comparable depth (African-lineage cluster medians ≈311–617 kyr). Distinguishing pre-out-of-Africa-shared from European-private lineages at single-variant resolution requires polarity-aware cross-population analysis and is deferred to future work.

To test whether the cluster-level age stratification within the EUR-coalescent frame is a signal-driven feature of the credible-set substrate rather than an artefact of estimator behaviour at the cluster boundaries, we compared PGC3 credible-set variants with minor-allele-frequency (MAF)- and linkage-disequilibrium (LD)-matched HapMap3 controls (Supplementary Methods §MAF-and-LD-matched control baseline; Supplementary Table 17). PGC3 variants are modestly younger than matched controls within this deep, predominantly pre-out-of-Africa window (median ≈377 kyr vs ≈472 kyr; 13,399 vs 16,809 generations × 28.1 yr generation□¹; Mann– Whitney U *P* < 10□³□□), and each of the three GMM-derived clusters (C0, C1, C2) shows distinct stratification against MAF-matched controls.

### Allele age and selection metrics

Per-variant median coalescent allele ages were obtained from the Atlas of Variant Age^19^ (https://human.genome.dating); for variants with multiple Atlas estimates, the highest-quality “Combined” estimate was preferred. The integrated haplotype score (|iHS|)^24^ was computed per variant with scikit-allel v1.3.13 on European-subset 1000 Genomes Phase 3 phased haplotypes^57^, with ancestral-allele polarisation against the Ensembl GRCh37 high-confidence ancestral FASTA; raw scores were standardised genome-wide in 50 derived-allele-count bins. Singleton Density Scores^25^ were obtained from the original UK10K-derived release and matched on rsID and GRCh37 position. We use SDS as a per-variant frequency-derivative signal and not as a polygenic-adaptation test, given the documented inflation of polygenic-adaptation inference by uncorrected population stratification^37,38^. Ancient-DNA-derived selection statistics for 9,739,624 quality-controlled variants^22^ were downloaded from the Reich Lab Harvard Dataverse deposit; allele orientation was reconciled by forward and reverse-complement match for non-palindromic strand-flipped variants, with palindromic A/T and C/G variants dropped from sign-sensitive Akbari analyses, yielding 19,564 of 20,637 PGC3 variants (94.8%) matched. Regional background-selection scores^58,59^ were used as a contextual annotation.

#### cis-eQTL data

Three *cis*-eQTL data sources were integrated. The primary tissue-level layer was GTEx v10 fine-mapped per-tissue significant variant–gene pairs^60^, processed by taking the minimum nominal *P*- value per PGC3 variant across all tested geneswithin each of 13 brain tissues and three blood/immune tissues (with per-tissue allele-age correlations reported in Supplementary Table 9). Variants were lifted from GRCh37 to GRCh38, with allele orientation reconciled across forward, flip, forward-rev-comp and flip-rev-comp branches; palindromic A/T and C/G hits were retained but flagged. The single-cell brain layer used Bryois et al.^34^ summary statistics across eight cell types, yielding per-cell-type per-rsID minimum *P*-valuesat 89.7% coverage. The PsychENCODE 2 atlas^61^ was used at the bulk level as a cross-atlas consistency check. Full data provenance, brain–blood specificity derivation and tissue-power matching are detailed in Supplementary Methods §eQTL data and §Tissue-power rescaling. We did not apply colocalisation^62^ or SMR/HEIDI^63^, as the primary inference concerns heritability concentration in cluster annotations rather than per-locus causal-gene assignment.

Brain–blood regulatory specificity, the second clustering axis, was defined per variant as brain_spec = b / (b + l), where b = −log (P_brain) and l = −log (P_blood) and P_brain (P_blood) is the minimum nominal cis-eQTL P across all GTEx v10 genome-wide-significant variant–gene pairs in the brain (blood/immune) tissue group, each floored at 1□×□10□³□□. Scores lie in (0, 1); values > 0.5 indicate a brain-biased regulatory profile. Significance was inherited from the GTEx significant-pair calls, with no additional threshold applied.

### Within-locus partial rank correlation

Variant-pooled and locus-aggregate correlations can disagree because individual loci tend to concentrate variants of similar age and similar regulatory effect through joint demographic and ascertainment processes, producing a Simpson’s-paradox structure. We therefore used within-locus residualisation as the primary analytical framework. For each variant in each credible set with ≥ 5 variants, the two focal features *x* and *y* were converted to within-credible-set ranks; when MAF residualisation was requested, the within-credible-set MAF rank was computed in parallel and the within-locus ranks of *x* and *y* were residualised against the within-locus MAF rank by ordinary-least-squares regression. Residualised ranks were pooled across the 249 non- *MAPT* credible sets, and the within-locus partial Spearman ρ was reported as the Pearson correlation on the pooled within-locus rank residuals, with the asymptotic Pearson *P*-value reported alongside.Confidence intervals were obtained from per-locus block bootstrap (below). Full procedural detail, including inverse-variance pooling under the small-sample Spearman variance approximation, is given in Supplementary Methods §Within-locus partial-rank residualisation. Multiple testing across the eight primary correlations reported in Table 3 was controlled with the Benjamini–Hochberg false-discovery rate across the eight primary correlations, in line with practical guidance^64^.

### Bootstrap confidence intervals

95% percentile confidence intervals were computed from 1,000 iterations of a per-locus block bootstrap in which credible sets were resampled with replacement and the full within-locus and MAF-rank residualisation was recomputed inside each iteration. Bootstrap design and the rationale for inside-iteration recomputation are detailed in Supplementary Methods §Block bootstrap confidence intervals.

### Gaussian mixture models

Gaussian mixture models were fit on the standardised joint distribution of log□□ allele age, brain–blood regulatory specificity and |iHS| using scikit-learn with full covariance, ten random initialisations per *k* (random_state = 42). BIC and AIC both supported multimodality over a single-component fit (ΔBIC_k=3 vs. k=1_ = −3,554 with 17q21.31 excluded; ΔBIC_k=3 vs. k=1_ < −15,000 with 17q21.31 included) but continued to decrease monotonically as *k* increased, a known over- fitting tendency of likelihood-based criteria in low-sample Gaussian mixture estimation^65^. We therefore selected *k* on the three clustering-specific metrics that are less susceptible to this monotonic drift (Integrated Completed Likelihood, silhouette and Davies–Bouldin), all three of which independently converged on *k* = 3, and we adopted *k* = 3 for parsimony and interpretability. The full 6-metric battery is reported in Supplementary Table 5f and the selection rationale in Supplementary Methods §GMM component selection. Mixture components were ordered by unscaled mean log allele age. Cluster-membership stability at *k* = 3 was assessed by 1,000 bootstrap re-fits (bootstrap mean adjusted Rand index = 0.967 ± 0.019; 95% percentile CI [0.919, 0.990]).

### Prior three-regime specification

Three variant subsets were specified before clustering, defined by joint position in (coalescent allele age × |iHS| × brain–blood regulatory specificity) space rather than by absolute age: (i) a relatively young subset with the youngest coalescent ages and low-to-moderate |iHS|; (ii) an intermediate subset with elevated |iHS|, capturing recent haplotype-extending (partial-sweep) signatures, which can act on standing variation of any age; and (iii) a relatively old subset with the longest coalescent ages and low |iHS|, consistent with background-selection-tolerant maintenance at common allele frequency^13^. Because Atlas of Variant Age estimates are reported in generations (converted at 28.1 yr generation ¹), all three subsets fall before the out-of-Africa dispersal (cluster medians ≈113–508 kyr); the subset labels therefore denote relative coalescent- age strata within a pre-out-of-Africa range, not absolute recent or post-out-of-Africa origin.

### Negative controls

Two complementary negative-control sets were used. The first comprised MAF- and LD-score-matched non-PGC3 HapMap3 variants (per-SNP matching tolerance of one MAF decile and one LD-score decile; *n* = 20,637 PGC3 variants matched against ≈2.05 M controls; Wilcoxon paired tests). The second comprised within-window neighbour single-nucleotide variants drawn from 1000 Genomes European phased haplotypes^57^ within ± 50 kb of each PGC3 variant. Both designs are detailed in Supplementary Methods §MAF-and-LD-matched control baseline and §Neighbour-control comparison.

### Partitioned LD score regression

Partitioned LDSC was applied per the LDSC v1.0.1 framework^66,67^ with seven minor Python-3 compatibility patches (Supplementary Methods §LDSC Python-3 patches). These patches are strictly syntactic and do not alter any statistical procedure or numerical input. They do not include the Tashman et al.^35^ small-annotation type-1-error mitigation, which we addressed separately via the --n-blocks 1000 sensitivity (Supplementary Table 15) and the empirical matched-LD-MAF permutation null (Supplementary Table 20). The conditional coefficient *Z*- score is the standardised regression coefficient for the focal annotation in a model that simultaneously includes the 97-annotation baseline-LD v2.2 model^27^, comprising the 53 functional annotations of Finucane et al.^67^ extended with 10 MAF bins, 6 LD-related annotations (including a MAF-adjusted predicted-allele-age annotation), McVicker background-selection coefficient, recombination rate, CpG content, replication timing, conserved-primate sequence annotations, and 11 additional annotations from Hujoel et al.^68^, Villar et al.^69^ and Marnetto et al.^70^, including ancient-sequence-age annotations for human promoters and enhancers (precise list at the Steven Gazal Zenodo deposit https://zenodo.org/records/10515792 in readme_baseline_versions.txt). The S-LDSC fold-enrichment statistic is the ratio of the partition heritability estimate (prop_h²) to the partition SNP fraction (prop_SNPs); because prop_h² is a regression estimate rather than a constrained proportion, it can take values slightly below zero for annotations whose true contribution is negligible, yielding negative fold values that are statistically indistinguishable from zero given the associated standard error. We retain such negative point estimates in the supplementary tables for transparency and interpret them as no enrichment rather than as depletion. PGC3 EUR SCZ^6^ was munged to LDSC format with HapMap3 restriction (1,171,345 final SNPs); Wightman et al.^29^ Alzheimer’s disease and PGC3 EAS schizophrenia^6^ sumstats were munged with palindromic pre-filter and reverse-complement allele matching (1,190,301 and 1,073,245 final SNPs, respectively). For EAS, *N*_eff_ = 4 · *N*_cas_ · *N*_con_ / (*N*_cas_ + *N*_con_) was computed per row (mean *N*_eff_ = 30,231). Cluster annotation files were built per chromosome and matched to either the EUR baseline-LD v2.2 SNP set or the EAS baseline-LD v2.2 SNP set (also from the Zenodo 10515792 deposit); LD scores were computed via ldsc.py --l2 --ld-wind-cm 1 per chromosome. Regression was run with --overlap-annot --frqfile-chr --print-coefficients. For EAS, cluster .l2.ldscore.gz files were post-filtered to the EAS baseline regression-SNP set. PolyFun^71^, MAGMA^72^, sc-linker^73^ and SMR/HEIDI^63^ were considered as alternative frameworks but not applied here (Supplementary Methods §LDSC alternative frameworks).

### Long-range LD region exclusion

The 24 long-range LD regions catalogued in GRCh37 coordinates by Price et al.^28^ were masked from the cluster annotation files; LD scores were recomputed from the masked annotations and partitioned heritability was re-run on PGC3 EUR schizophrenia and on Wightman et al.^29^ Alzheimer’s disease (Supplementary Table 6).

### Robustness battery

The principal partitioned heritability finding on PGC3 EUR schizophrenia was tested along a seven-axis robustness battery comprising block-jackknife (Tashman) sensitivity, feature-space sensitivity (2D age + |iHS|), *MAPT*-included sensitivity, brain-specificity 4-test adjudication, tissue-power-matched brain-specificity sensitivity, empirical matched-LD-MAF permutation null, and component-number (*k*) sensitivity. The principal Young-cluster heritability concentration is preserved across all seven axes under the baseline-LD v2.2 model (Supplementary Tables 14, 15, 16, 18,19, 19A, 20). Full procedural detail and per-axis numerical results are reported in Supplementary Methods Part G (§Robustness and sensitivity battery, §G1 to §G7), with each axis cross-referenced to its mechanistic description in Parts B, D and E.

### Sex-stratified replication

Sex-stratified PGC3 wave 3 daner-format summary statistics were munged to LDSC format using a daner-aware adapter (Supplementary Methods §Daner-aware munge). Effective sample sizes were 68,003 (EUR male), 47,652 (EUR female), 13,017 (EAS male) and 13,163 (EAS female). Sex-specificity was assessed by a heterogeneity test, *Z*_diff_ = (enrichment_male_ − enrichment_female_) / √(s.e.²_male_ + s.e.²_female_), under the independent-stratum assumption.

### Cross-disorder analyses

Partitioned LDSC (cluster annotations × baseline-LD v2.2 × EUR LDSC reference) was applied to six additional disorder summary-statistic substrates: PGC bipolar disorder^30^, PGC major depressive disorder^74^, PGC ADHD^75^, iPSYCH-PGC autism^76^, and the CDG3 Genomic-SEM-derived F3 and F4 factors^52^. The CDG3 p-factor and F2 factor were excluded because both contain direct schizophrenia contribution at the underlying multivariable LDSC stage, which would induce sample-overlap bias when intersected with our PGC3 SCZ-derived cluster annotations. The three munge implementations required (daner-aware, minimal-daner and Genomic-SEM-tsv) and per-trait implied *N* are documented in Supplementary Methods §Cross-disorder munge implementations. Disorder-level multiple testing used six-test Bonferroni correction (α = 0.05/6 = 0.0083); within each disorder the conditional coefficient *Z*-score was evaluated against a focal-annotation Bonferroni threshold of approximately Z = 2.81 (one-tailed α = 2.4 × 10□³). This threshold derives from 21 focal-annotation hypothesis tests (three clusters × seven phenotypes) under a baseline-LD v2.2 reference, following the convention that the 97 baseline-LD annotations are regression-level control covariates rather than independent hypothesis tests^67,68^. Admixed AFR and Latino summary statistics were not used, as cov-LDSC^50^ is required for admixed populations and requires in-sample LD reference data that are not available without raw-genotype access. The r_g-prediction model used to benchmark cross-disorder enrichments against schizophrenia genetic correlation is detailed in Supplementary Methods §r_g-prediction.

### Negative-control external traits

Identical partitioned LDSC was applied to two well-powered non-psychiatric, non-brain comparators: the GIANT 2018 height and body-mass index meta-analyses^31^. Summary statistics were obtained from the GIANT consortium portal and munged using a GIANT-aware adapter (Supplementary Methods §GIANT-aware munge); 1,010,434 (height) and 1,011,649 (BMI) HapMap3 SNPs were retained. Partitioned LDSC was run with the same EUR cluster annotations and baseline-LD v2.2 model used for discovery (Supplementary Table 13).

### Computational environment

Variant-level genotype manipulation used PLINK 2^77^, BCFtools/SAMtools^78^ and VCFtools^79^; numerical and scientific computing used NumPy^80^ and SciPy^81^. Analysis-specific Python libraries (scikit-allel v1.3.13 for |iHS|, scikit-learn for Gaussian mixture modelling, ldsc.py for partitioned heritability) are noted alongside the analyses they supported.

### Statistics and reproducibility

Gaussian mixture models were fitted with scikit-learn GaussianMixture (n_components = 3, covariance_type = "full", n_init = 10, random_state = 42) on z-score-standardised log allele age, brain–blood specificity and |iHS| for the 4,918 non-MAPT variants, with components ordered by mean log age to label Young, Mid and Old. Within-locus correlations are Spearman partial rank correlations after double residualisation (within-credible-set mean-centring then MAF-rank residualisation); confidence intervals are percentile intervals from a 1,000-iteration per-locus block bootstrap (resampling credible sets; seed 42). Multiple testing across the eight primary within-locus correlations (Table 3) used the Benjamini–Hochberg false-discovery rate. Partitioned-heritability significance is the conditional coefficient Z against the focal-annotation threshold (Z ≈ 2.81); the reported P is the LDSC block-jackknife enrichment test. Cell-type coverage contrasts are three-way χ² tests; matched-control and paired age comparisons are two-sided Mann–Whitney U and Wilcoxon signed-rank tests. Where a test statistic underflows the floating-point floor, P is reported as the software limit (P < 10 ³). Sample size n is the number of variants unless loci or LD-reference SNPs are specified. No randomisation or blinding applied, and no statistical method predetermined sample size.

## Data availability and code availability

PGC3 schizophrenia GWAS summary statistics and fine-mapped credible sets^6^ are publicly available from the Psychiatric Genomics Consortium (https://www.med.unc.edu/pgc/). Atlas of Variant Age^19^ coalescent allele-age estimates are available at https://human.genome.dating. GTEx v10 cis-eQTL summary statistics^60^ are available from the GTEx Portal (https://gtexportal.org). Bryois 2022 single-cell brain cis-eQTL summary statistics^34^ are available from the corresponding Zenodo deposit. Akbari 2026 ancient-DNA-derived selection coefficients^22^ are available from the Reich Lab Harvard Dataverse deposit. Field 2016 Singleton Density Scores^25^ are available from the original UK10K release. 1000 Genomes Phase 3 phased haplotypes^57^ are available from the 1000 Genomes Project portal. gnomAD v4 gene-level constraint^82,83^ is available from https://gnomad.broadinstitute.org. Baseline-LD v2.2^27^ EUR and EAS LD-score reference files are publicly available from the Steven Gazal Zenodo deposit https://zenodo.org/records/10515792 (DOI 10.5281/zenodo.10515792). PGC bipolar disorder^30^, PGC major depression^74^, PGC ADHD^75^, iPSYCH-PGC autism^76^ and CDG3 F3/F4 Genomic- SEM factors^52^ are available from the corresponding PGC and Cross-Disorder Group portals. Yengo 2018 GIANT height and BMI summary statistics^31^ are available from the GIANT consortium. The full analysis pipeline (Python 3.11) is publicly available at https://github.com/dryusufcicek/EVOSCZ (release v1.0) and archived at Zenodo (DOI 10.5281/zenodo.21020749); the per-variant master table, cluster assignments and partitioned-heritability result files are regenerated by this code from the public source data listed above.

## Web Resources

- Psychiatric Genomics Consortium portal: https://www.med.unc.edu/pgc/
- Atlas of Variant Age: https://human.genome.dating
- GTEx Portal: https://gtexportal.org
- gnomAD v4 browser: https://gnomad.broadinstitute.org
- Reich Lab Harvard Dataverse: https://dataverse.harvard.edu/dataverse/reichlab
- GIANT consortium: https://portals.broadinstitute.org/collaboration/giant/
- LDSC v1.0.1 (Bulik-Sullivan, Finucane): https://github.com/bulik/ldsc
- Steven Gazal S-LDSC reference files (baseline + baseline-LD v1.2 / v2.2 / v2.3 for EUR and EAS): https://zenodo.org/records/10515792
- EVOSCZ analysis pipeline (this study): https://github.com/dryusufcicek/EVOSCZ

## Author contributions (CRediT)

Y.Ç.: Conceptualization, Data Curation, Formal Analysis, Investigation, Methodology, Software, Visualization, Writing – Original Draft. A.T.A.: Conceptualization, Methodology, Supervision, Writing – Review & Editing. Ö.F.D.: Supervision, Resources, Writing – Review & Editing. H.A.V.: Supervision, Writing – Review & Editing.

## Inclusion and ethics

This study used publicly available, de-identified GWAS summary statistics and reference resources (PGC3 schizophrenia, PGC bipolar disorder, PGC major depression, PGC ADHD, iPSYCH–PGC autism, CDG3 Genomic-SEM factors, GIANT 2018, Wightman 2021 Alzheimer’s disease, GTEx v10, Bryois 2022 single-cell brain *cis*-eQTL, Akbari 2026 ancient-DNA selection coefficients, Field 2016 Singleton Density Scores, Atlas of Variant Age, 1000 Genomes Phase 3, gnomAD v4, baseline-LD v2.2). No new human-subjects research was conducted and no individual-level genotype or phenotype data were accessed. The authors are based at institutions in Türkiye and acknowledge contributions from the international consortia listed above.

## Declaration of generative AI

In accordance with standard editorial policy on the use of generative AI in scientific publishing, the authors disclose that generative AI tools (Anthropic Claude, model versions Opus 4.6, 4.7 and 4.8) were used during manuscript preparation to draft prose from author-provided outlines and analytical outputs, to edit language for clarity and concision, to discuss analytical and interpretive options, and to review and debug analysis scripts. All statistical analyses were run by the authors using established open-source software (LDSC v1.0.1, scikit-learn, SciPy, scikit-allel, PLINK 2, bedtools, pysam, BCFtools; full versions in the code-availability environment file), and no data, results or images were generated by generative AI. The authors retain full responsibility for the integrity, accuracy, originality, and scientific content of the manuscript.

## Acknowledgments

We thank the Psychiatric Genomics Consortium and all contributing investigators, study participants and consortium analysts for the generation and release of PGC3 schizophrenia, bipolar, major depression and ADHD summary statistics, including the sex-stratified PGC3 wave 3 daner-format files that enabled the sex-stratified replication. We thank the GTEx Consortium, the Bryois single-cell brain *cis*-eQTL contributors, the iPSYCH-PGC autism collaboration, the Cross-Disorder Group of the PGC, the Akbari ancient-DNA cohort, the 1000 Genomes Project, the Atlas of Variant Age developers, the GIANT consortium and Steven Gazal for public data release. We thank the Cerrahpaşa Faculty of Medicine and The Scientific and Technological Research Council of Türkiye (TÜBİTAK) for computational resources. The numerical calculations reported in this paper were partially performed at TÜBİTAK ULAKBİM, High Performance and Grid Computing Center (TRUBA resources).

