## Supplementary Tables for "Common schizophrenia heritability concentrates in an evolutionarily young, brain-regulatory subset of fine-mapped credible-set variants"

All P-values are reported to one significant figure in the mantissa (e.g. 1.4 × 10⁻¹⁶); correlations are reported to three decimal places. Negative numbers use the Unicode minus sign (−). Bonferroni P-values are capped at 1.000. Sample sizes are written with thousand separators. Abbreviations are defined in the glossary below and at first use.

**Abbreviations:** GEVA, Genealogical Estimation of Variant Age; TGP, 1000 Genomes Project; SGDP, Simons Genome Diversity Project; TMRCA, time to most recent common ancestor; MAF, minor allele frequency; LD, linkage disequilibrium; OOA, out-of-Africa; S-LDSC, stratified LD-score regression; GMM, Gaussian mixture model; |iHS|, absolute integrated haplotype score; SDS, singleton density score; LOEUF, loss-of-function observed/expected upper-bound fraction; eQTL, expression quantitative trait locus; pp, percentage points; daner, PGC summary-statistics format.

**Supplementary Table 1 | Variant master annotation coverage across all 20,637 autosomal PGC3 fine-mapped credible-set variants.**

Coverage of each annotation in the variant master file underlying the analyses in Trubetskoy et al. 2022¹ PGC3 schizophrenia credible sets, ordered by data source.

| **Annotation** | **Source** | **n variants annotated** | **Coverage (%)** | **Notes** |  |  |
| --- | --- | --- | --- | --- | --- | --- |
| Coalescent allele age (median, years) | Atlas of Variant Age (Albers and McVean 2020²) | 20,565 | 99.7 | Combined preferred over TGP/SGDP-only |  |  |
| Singleton Density Score | Field 2016 (UK10K⁵) | 14,843 | 71.9 | rsID + GRCh37 position match |  |  |
| GTEx v10 brain min cis-eQTL P | GTEx Consortium 2020⁴ | 10,727 | 52.0 | Min nominal P across 13 brain tissues; lifted hg19 to hg38 |  |  |
| GTEx v10 blood/immune min cis-eQTL P | GTEx Consortium 2020⁴ | 8,527 | 41.3 | Min nominal P across whole blood and spleen |  |  |
| Voight-correct ancestral-polarised genome-wide-DAF-binned standardised | iHS |  | Voight 2006³; 1000G Phase 3 EUR¹⁷ | 17,352 | 84.1 | scikit-allel v1.3.13 with Ensembl GRCh37 ancestral FASTA; min_ehh = 0.05; min_maf = 0.05 |
| Akbari 2026 ancient-DNA selection coefficient | S |  | Akbari et al. 2026⁷ | 19,564 | 94.8 | Reverse-complement allele matching; palindromic A/T,C/G dropped from sign-sensitive analyses |
| gnomAD v4 LOEUF (gene-level) | Karczewski 2020; gnomAD v4¹³,³⁴ | 9,464 | 45.9 | Constraint mapped via nearest gene |  |  |
| gnomAD v4 pLI (gene-level) | Karczewski 2020; gnomAD v4¹³,³⁴ | 9,464 | 45.9 | Same gene mapping as LOEUF |  |  |
| gnomAD v4 missense Z (gene-level) | Karczewski 2020; gnomAD v4¹³,³⁴ | 9,505 | 46.1 | Same gene mapping |  |  |
| CADD PHRED | Functional annotation⁴⁹ | 20,637 | 100.0 | Variant-level deleteriousness |  |  |
| Posterior inclusion probability (FINEMAP) | Trubetskoy 2022¹; Benner 2016¹⁶ | 20,637 | 100.0 | PGC3 fine-mapped credible-set; complete by ascertainment |  |  |
| GWAS effect size β | Trubetskoy 2022¹ | 20,637 | 100.0 | PGC3 EUR primary |  |  |
| Minor allele frequency | Trubetskoy 2022¹ | 20,637 | 100.0 | PGC3 EUR primary |  |  |
| Ensembl ancestral allele (GRCh37) | Ensembl GRCh37 | 20,565 | 99.7 | Uppercase high-confidence; low-confidence and unknown excluded |  |  |
| HAR overlap (Cui 2025 v3) | Cui et al. 2025¹² | 20,637 | 100.0 | Variant-level overlap; 0/1 indicator |  |  |
| Bryois 2022 Astrocytes min cis-eQTL P | Bryois et al. 2022⁹ | 16,911 | 81.9 | Min P across all tested genes per rsID per cell type |  |  |
| Bryois 2022 Endothelial cells | Bryois et al. 2022⁹ | 15,187 | 73.6 | Same |  |  |
| Bryois 2022 Excitatory neurons | Bryois et al. 2022⁹ | 18,385 | 89.1 | Same |  |  |
| Bryois 2022 Inhibitory neurons | Bryois et al. 2022⁹ | 18,350 | 88.9 | Same |  |  |
| Bryois 2022 Microglia | Bryois et al. 2022⁹ | 14,700 | 71.2 | Same |  |  |
| Bryois 2022 OPCs and COPs | Bryois et al. 2022⁹ | 17,942 | 86.9 | Same |  |  |
| Bryois 2022 Oligodendrocytes | Bryois et al. 2022⁹ | 16,347 | 79.2 | Same |  |  |
| Bryois 2022 Pericytes | Bryois et al. 2022⁹ | 14,070 | 68.2 | Same |  |  |
| Bryois 2022 any of 8 cell types | Bryois et al. 2022⁹ | 18,519 | 89.7 | Aggregate single-cell brain cis-eQTL coverage |  |  |

*Coverage is the number of variants with a non-missing value for each annotation, divided by the total of 20,637 autosomal credible-set variants. The total of 20,637 is the autosomal subset of the variant master file; the corresponding count in Trubetskoy et al. 2022 was 20,766, from which 129 chromosome-X variants were excluded under the autosomal restriction (20,766 − 129 = 20,637). See Methods for the per-source ascertainment protocol. Bryois 2022 cell-type coverage is per cell type; the 'any of 8 cell types' row is the union. SDS, Singleton Density Score; eQTL, expression-quantitative-trait locus; LOEUF, loss-of-function observed/expected upper bound fraction; pLI, probability of loss-of-function intolerance; CADD, Combined Annotation-Dependent Depletion; PIP, posterior inclusion probability; HAR, human accelerated region; OPC, oligodendrocyte precursor cell; COP, committed oligodendrocyte precursor. This annotation master underlies the variant substrate used in all main-text analyses (Figs 1–5).*

**Supplementary Table 2 | Within-locus partial rank correlation between brain–blood regulatory specificity and log₁₀ allele age, with bootstrap and matched-control sensitivity tests.**

Primary specification and sensitivity analyses for the brain–blood specificity × log₁₀ allele age within-credible-set correlation reported in the main text.

| **Test** | **Specification** | **n variants (loci)** | **Spearman ρ** | **Asymptotic P** | **Bootstrap 95% CI (1,000-iteration per-locus block, residualise inside loop)** |
| --- | --- | --- | --- | --- | --- |
| Brain–blood specificity × log₁₀ allele age | within-locus rank, no MAPT | 5,724 (84) | −0.055 | 3.6 × 10⁻⁵ | n/a |
| Brain–blood specificity × log₁₀ allele age | within-locus rank, MAF-rank residualised, no MAPT | 5,724 (84) | −0.093 | 1.8 × 10⁻¹² | [−0.242, +0.021] |
| Brain–blood specificity × log₁₀ allele age | within-locus rank + MAF, all loci including 17q21.31 (MAPT) | 7,398 (85) | −0.021 | 0.072 | [−0.196, −0.016] |
| Brain–blood specificity × log₁₀ allele age (paired matched-control) | MAF and LD-score-decile-matched HapMap controls; per-locus paired Wilcoxon | 85 loci paired | PGC3 median −0.043; control median +0.012 | 0.330 | n/a (paired Wilcoxon test) |
| Brain–blood specificity × log₁₀ allele age (within-window neighbour control) | ±50 kb 1000G EUR neighbour SNVs; PGC3 excluded by chr/pos match | 7,173 (PGC3); 1,733 (neighbours) | PGC3 −0.021; neighbour −0.014 | PGC3 0.077; neighbour 0.560 | n/a (paired by window) |

*Brain–blood regulatory specificity is defined per variant as b_spec = −log₁₀ P_brain / [−log₁₀ P_brain + −log₁₀ P_blood] (Methods). The asymptotic P-value uses the standard Pearson formula on pooled within-locus rank residuals; the per-locus block-bootstrap CI places credible sets as the resampling unit and recomputes residualisation inside each iteration. Both controls compare schizophrenia credible-set variants to MAF and LD-score-decile-matched HapMap variants and to within-window neighbour SNVs; see Methods.*

**Supplementary Table 3 | Five alternative formulations of brain–blood regulatory specificity tested for sign concordance.**

Within-locus partial rank correlation of each specificity formulation against log₁₀ allele age, in the full set of credible-set variants and after excluding 17q21.31 (MAPT inversion).

| **Formulation** | **Description** | **Definition** | **Subset** | **n variants** | **Spearman ρ** | **Direction** | **P** |  |  |  |  |
| --- | --- | --- | --- | --- | --- | --- | --- | --- | --- | --- | --- |
| M1 | Primary ratio specification | −log₁₀(P_brain) / [−log₁₀(P_brain) + −log₁₀(P_blood)] | All credible sets | 7,398 | −0.021 | negative | 0.072 |  |  |  |  |
| M1 | Primary ratio specification | −log₁₀(P_brain) / [−log₁₀(P_brain) + −log₁₀(P_blood)] | Excluding 17q21.31 MAPT inversion | 5,724 | −0.093 | negative | 1.8 × 10⁻¹² |  |  |  |  |
| M2 | Log-ratio of −log₁₀ P signals | log₁₀[(−log₁₀ P_brain + 1) / (−log₁₀ P_blood + 1)] | All credible sets | 7,398 | −0.140 | negative | 1.3 × 10⁻³³ |  |  |  |  |
| M2 | Log-ratio of −log₁₀ P signals | log₁₀[(−log₁₀ P_brain + 1) / (−log₁₀ P_blood + 1)] | Excluding 17q21.31 MAPT inversion | 5,724 | +0.093 | positive | 1.6 × 10⁻¹² |  |  |  |  |
| M3 | Standardised Z-score difference | Z(−log₁₀ P_brain) − Z(−log₁₀ P_blood) | All credible sets | 7,398 | −0.039 | negative | 9.2 × 10⁻⁴ |  |  |  |  |
| M3 | Standardised Z-score difference | Z(−log₁₀ P_brain) − Z(−log₁₀ P_blood) | Excluding 17q21.31 MAPT inversion | 5,724 | −0.081 | negative | 7.3 × 10⁻¹⁰ |  |  |  |  |
| M4 | Dominance categorical encoding | Categorical: brain-only / blood-only / both / neither | All credible sets | 3,843 | −0.205 | negative | 1.4 × 10⁻³⁷ |  |  |  |  |
| M4 | Dominance categorical encoding | Categorical: brain-only / blood-only / both / neither | Excluding 17q21.31 MAPT inversion | 3,843 | −0.205 | negative | 1.4 × 10⁻³⁷ |  |  |  |  |
| M5 | Effect-size-based comparison |  | β_brain | − | β_blood | (effect-size slope difference) | All credible sets | 7,398 | +0.180 | positive | 1.2 × 10⁻⁵⁴ |
| M5 | Effect-size-based comparison |  | β_brain | − | β_blood | (effect-size slope difference) | Excluding 17q21.31 MAPT inversion | 5,724 | −0.142 | negative | 4.1 × 10⁻²⁷ |

*Four of the five formulations (M1, M3–M5) are negative for the within-locus age × specificity correlation when the 17q21.31 MAPT inversion is excluded; M2 (the log-ratio formulation) is weakly positive (ρ = +0.093). M1 is the primary specification used throughout the main text; M2–M5 are sensitivity formulations. Spearman ρ is the within-locus rank residual correlation; P is the asymptotic Pearson P on pooled rank residuals (Methods).*

**Supplementary Table 4 | Cross-method consistency of selection metrics, allele age and gene constraint at PGC3 credible-set variants.**

Within-locus partial rank correlations for 22 primary cross-method tests, with Benjamini-Hochberg FDR and per-test Bonferroni multiplicity adjustment.

| **Test** | **n variants** | **Spearman ρ** | **Asymptotic P** | **BH-FDR q** | **Bonferroni P (n_tests = 22)** | **Pass BH-FDR 5%** | **Pass Bonferroni 5%** |  |  |  |  |
| --- | --- | --- | --- | --- | --- | --- | --- | --- | --- | --- | --- |
| Brain eQTL × LOEUF | 5,393 | +0.443 | 3.6 × 10⁻²⁵⁸ | 7.9 × 10⁻²⁵⁷ | 7.9 × 10⁻²⁵⁷ | yes | yes |  |  |  |  |
|  | Akbari S | × SDS | 14,182 | +0.274 | 1.9 × 10⁻²⁴³ | 2.1 × 10⁻²⁴² | 4.2 × 10⁻²⁴² | yes | yes |  |  |
| Brain spec × LOEUF | 4,601 | +0.368 | 2.1 × 10⁻¹⁴⁷ | 1.5 × 10⁻¹⁴⁶ | 4.6 × 10⁻¹⁴⁶ | yes | yes |  |  |  |  |
| Brain spec × | Akbari S |  | 6,978 | +0.226 | 3.6 × 10⁻⁸¹ | 2.0 × 10⁻⁸⁰ | 7.8 × 10⁻⁸⁰ | yes | yes |  |  |
| Brain eQTL × age | 10,630 | −0.177 | 3.0 × 10⁻⁷⁵ | 1.3 × 10⁻⁷⁴ | 6.5 × 10⁻⁷⁴ | yes | yes |  |  |  |  |
| Brain spec × Akbari S signed | 6,978 | +0.180 | 1.2 × 10⁻⁵¹ | 4.2 × 10⁻⁵¹ | 2.5 × 10⁻⁵⁰ | yes | yes |  |  |  |  |
| Blood eQTL × age | 8,457 | −0.142 | 3.1 × 10⁻³⁹ | 9.8 × 10⁻³⁹ | 6.8 × 10⁻³⁸ | yes | yes |  |  |  |  |
|  | iHS | × age (within, MAF) | 17,280 | +0.097 | 1.5 × 10⁻³⁷ | 4.0 × 10⁻³⁷ | 3.2 × 10⁻³⁶ | yes | yes |  |  |
|  | iHS | × age, no 17q21.31 | 15,668 | −0.089 | 1.0 × 10⁻²⁸ | 2.5 × 10⁻²⁸ | 2.3 × 10⁻²⁷ | yes | yes |  |  |
|  | Akbari S | × age | 19,428 | −0.076 | 2.3 × 10⁻²⁶ | 5.0 × 10⁻²⁶ | 5.0 × 10⁻²⁵ | yes | yes |  |  |
| Brain eQTL × age, no 17q21.31 | 8,956 | −0.086 | 3.4 × 10⁻¹⁶ | 6.7 × 10⁻¹⁶ | 7.4 × 10⁻¹⁵ | yes | yes |  |  |  |  |
| Brain spec × age, no 17q21.31 | 5,724 | −0.093 | 1.8 × 10⁻¹² | 3.4 × 10⁻¹² | 4.1 × 10⁻¹¹ | yes | yes |  |  |  |  |
| Blood eQTL × age, no 17q21.31 | 6,783 | +0.069 | 1.2 × 10⁻⁸ | 2.0 × 10⁻⁸ | 2.7 × 10⁻⁷ | yes | yes |  |  |  |  |
|  | iHS | × SDS | 14,239 | −0.044 | 2.0 × 10⁻⁷ | 3.1 × 10⁻⁷ | 4.3 × 10⁻⁶ | yes | yes |  |  |
|  | Akbari X | × age | 19,428 | −0.036 | 6.5 × 10⁻⁷ | 9.6 × 10⁻⁷ | 1.4 × 10⁻⁵ | yes | yes |  |  |
|  | iHS | × | Akbari X |  | 16,572 | −0.024 | 0.002 | 0.003 | 0.050 | yes | no |
|  | Akbari S | × age, no 17q21.31 | 17,715 | −0.021 | 0.005 | 0.007 | 0.113 | yes | no |  |  |
|  | iHS | × | Akbari S |  | 16,572 | −0.020 | 0.010 | 0.012 | 0.212 | yes | no |
| Brain spec × age (within, MAF) | 7,398 | −0.021 | 0.072 | 0.084 | 1.000 | no | no |  |  |  |  |
| SDS × age | 14,772 | −0.009 | 0.265 | 0.292 | 1.000 | no | no |  |  |  |  |
| SDS × age, no 17q21.31 | 14,581 | −0.009 | 0.291 | 0.305 | 1.000 | no | no |  |  |  |  |
| Akbari π × age | 18,848 | −0.002 | 0.820 | 0.820 | 1.000 | no | no |  |  |  |  |

*Tests are sorted by ascending P. The strongest cross-method triangulation is between |Akbari S| (Akbari et al. 2026 ancient-DNA-derived selection coefficient⁷) and Field 2016 SDS⁵: within-locus partial rank ρ = +0.274, P = 1.9 × 10⁻²⁴³, indicating the variant-level signal is robust across the ancient-DNA and modern-frequency-derivative frameworks. Bonferroni P uses n_tests = 22 and is capped at 1.000. Pass FDR 5%: BH-FDR q < 0.05. Pass Bonferroni 5%: per-test P < 0.05/22 = 2.27 × 10⁻³. eQTL, expression-quantitative-trait locus; SDS, Singleton Density Score; LOEUF, loss-of-function observed/expected upper bound fraction. These cross-method selection-metric checks support the within-locus analyses in Fig. 4.*

**Supplementary Table 5 | Gaussian mixture model selection, cluster centroids, top genes per cluster, and within-cluster correlations (17q21.31 MAPT-excluded).**

Five sub-tables describing the GMM model selection and cluster characterisation underlying the C0/C1/C2 partition.

**Section A. Bayesian information criterion across k = 1 to 6 (no MAPT)**

| **k components** | **BIC** | **AIC** | **log-likelihood** | **ΔBIC vs k = 1** |
| --- | --- | --- | --- | --- |
| 1 | 41875.4 | 41816.9 | −20899.5 | 0.0 |
| 2 | 39466.1 | 39342.6 | −19652.3 | −2409.4 |
| 3 | 38320.5 | 38132.0 | −19037.0 | −3554.9 |
| 4 | 38151.5 | 37898.0 | −18910.0 | −3723.9 |
| 5 | 38074.4 | 37755.8 | −18828.9 | −3801.1 |
| 6 | 37773.0 | 37389.5 | −18635.7 | −4102.4 |

**Section B. Per-cluster feature medians (17q21.31 MAPT-excluded)**

| **Feature** | **C0 (Young) n** | **C0 median** | **C1 (Mid) n** | **C1 median** | **C2 (Old) n** | **C2 median** |  |  |
| --- | --- | --- | --- | --- | --- | --- | --- | --- |
| Age (yr) | 1,745 | 112,682 | 317 | 357,570 | 2,856 | 508,337 |  |  |
| MAF | 1,745 | 0.3190 | 317 | 0.4100 | 2,856 | 0.3155 |  |  |
| GWAS β | 1,745 | 0.0350 | 317 | −0.0403 | 2,856 | 0.0149 |  |  |
| PIP | 1,745 | 0.0058 | 317 | 0.0038 | 2,856 | 0.0046 |  |  |
| CADD | 1,745 | 2.88 | 317 | 2.61 | 2,856 | 2.56 |  |  |
| Brain spec | 1,745 | 0.5109 | 317 | 0.4294 | 2,856 | 0.4963 |  |  |
|  | iHS |  | 1,745 | 0.5820 | 317 | 2.11 | 2,856 | 0.7180 |
| SDS | 1,426 | 0.2168 | 273 | −0.6753 | 2,353 | −0.0559 |  |  |
| Akbari S | 1,628 | 0.0005 | 310 | −0.0019 | 2,659 | 0.0002 |  |  |
| Akbari π | 1,628 | 0.0306 | 310 | 0.1067 | 2,659 | 0.0347 |  |  |
| LOEUF | 1,347 | 0.6036 | 243 | 0.8311 | 2,012 | 0.6402 |  |  |
| pLI | 1,347 | 0.0654 | 243 | 0.0000 | 2,012 | 0.0002 |  |  |
| −log10 brain p | 1,745 | 11.68 | 317 | 11.97 | 2,856 | 12.58 |  |  |
| −log10 blood p | 1,745 | 10.99 | 317 | 15.78 | 2,856 | 14.11 |  |  |

*Age (yr) is the per-cluster median GEVA/Atlas-of-Variant-Age allele age, reported in years after converting the source generation estimates at 28.1 years per generation (Wohns et al. 2022; main-text Methods and Limitations). The corresponding generation medians are C0 4,010, C1 12,725 and C2 18,090 generations. These cluster medians (≈113 / 358 / 508 thousand years) all predate the out-of-Africa expansion (~65 kyr) and reproduce main-text Table 1; the C0/C1/C2 ("Young"/"Mid"/"Old") labels denote the youngest, intermediate and oldest strata of a single deeply ancestral, pre-out-of-Africa allele set rather than recently arisen variation. All remaining feature rows are unchanged from the primary analysis.*

**Section C. Cluster centroids (unscaled feature space)**

| **Cluster** | **n GMM-assigned (no MAPT)** | **n LDSC EUR baseline annotation** | **Unscaled centroid log₁₀ age** | **Unscaled centroid age (years)** | **Unscaled centroid brain–blood specificity** | **Unscaled centroid** | **iHS** | **(Voight-correct)** |
| --- | --- | --- | --- | --- | --- | --- | --- | --- |
| C0 (Young) | 1,745 | 1,744 | 5.054 | 113,215 | 0.506 | 0.701 |  |  |
| C1 (Mid) | 317 | 317 | 5.557 | 360,102 | 0.404 | 1.892 |  |  |
| C2 (Old) | 2,856 | 2,855 | 5.709 | 511,083 | 0.491 | 0.740 |  |  |

Allele-age centroids are reported in years, obtained from the Atlas-of-Variant-Age² estimates — which are reported in generations — converted at 28.1 yr generation⁻¹ (Wohns 2022); the unscaled centroid log₁₀ age is on this year scale (log₁₀ age in generations + log₁₀ 28.1 = + 1.449). The three centroids correspond to ~113 kyr (C0, Young), ~360 kyr (C1, Mid) and ~511 kyr (C2, Old), consistent with main-text Table 1; all three lie deep in pre-Out-of-Africa time (OOA ~65 kyr), so 'Young/Mid/Old' denote the youngest, intermediate and oldest strata of one deeply ancestral, ancestrally shared variant set rather than recent post-OOA alleles. Brain–blood specificity and |iHS| centroids are unit-free and unchanged.

**Section D. Top 15 genes per cluster by variant count**

| **Cluster** | **Gene symbol** | **n variants** |
| --- | --- | --- |
| C0 (Young) | no gene assignment | 225 |
| C0 (Young) | ST3GAL3 | 84 |
| C0 (Young) | MARK3 | 74 |
| C0 (Young) | MAN2A1 | 48 |
| C0 (Young) | SPPL3 | 40 |
| C0 (Young) | GATAD2B | 40 |
| C0 (Young) | VPS45 | 36 |
| C0 (Young) | GIGYF2 | 35 |
| C0 (Young) | AC079807.4 | 32 |
| C0 (Young) | MLXIP | 30 |
| C0 (Young) | SPG7 | 27 |
| C0 (Young) | STAG1 | 27 |
| C0 (Young) | CYP7B1 | 25 |
| C0 (Young) | RERE | 25 |
| C0 (Young) | SZT2 | 25 |
| C1 (Mid) | SLC9B1 | 93 |
| C1 (Mid) | RP11-586K2.1 | 28 |
| C1 (Mid) | UBE2D3 | 25 |
| C1 (Mid) | no gene assignment | 23 |
| C1 (Mid) | WHSC1L1 | 13 |
| C1 (Mid) | MARK3 | 12 |
| C1 (Mid) | PPP1R13B | 12 |
| C1 (Mid) | RBM26 | 9 |
| C1 (Mid) | SZT2 | 8 |
| C1 (Mid) | EMB | 7 |
| C1 (Mid) | GIGYF2 | 7 |
| C1 (Mid) | GATAD2A | 7 |
| C1 (Mid) | ZNF664 | 6 |
| C1 (Mid) | OLA1 | 6 |
| C1 (Mid) | POC1B | 5 |
| C2 (Old) | no gene assignment | 324 |
| C2 (Old) | AC079807.4 | 144 |
| C2 (Old) | ALMS1 | 124 |
| C2 (Old) | CNNM2 | 117 |
| C2 (Old) | RP11-586K2.1 | 106 |
| C2 (Old) | NT5C2 | 77 |
| C2 (Old) | FANCA | 76 |
| C2 (Old) | ELAC2 | 69 |
| C2 (Old) | SMG6 | 63 |
| C2 (Old) | PCDHAC1 | 61 |
| C2 (Old) | MLXIP | 60 |
| C2 (Old) | PPP1R13B | 60 |
| C2 (Old) | PTK2B | 53 |
| C2 (Old) | WHSC1L1 | 49 |
| C2 (Old) | RERE | 48 |

**Section E. Within-cluster age × selection feature partial rank correlations**

| **Cluster** | **Test** | **n variants** | **Spearman ρ** | **P** |  |  |
| --- | --- | --- | --- | --- | --- | --- |
| C0 (Young) | brain_spec × age | 1,712 | −0.081 | 7.6 × 10⁻⁴ |  |  |
| C0 (Young) |  | iHS | × age | 1,712 | −0.146 | 1.1 × 10⁻⁹ |
| C0 (Young) | Akbari π × age | 1,487 | +0.226 | 9.6 × 10⁻¹⁹ |  |  |
| C1 (Mid) | brain_spec × age | 288 | −0.124 | 0.036 |  |  |
| C1 (Mid) |  | iHS | × age | 293 | +0.384 | 1.0 × 10⁻¹¹ |
| C1 (Mid) | Akbari π × age | 164 | +0.041 | 0.604 |  |  |
| C2 (Old) | brain_spec × age | 2,811 | +0.043 | 0.024 |  |  |
| C2 (Old) |  | iHS | × age | 2,811 | −0.020 | 0.292 |
| C2 (Old) | Akbari π × age | 2,592 | +0.049 | 0.013 |  |  |

*Section A: BIC, AIC and log-likelihood across k = 1 to 6, fitted on the standardised joint distribution of log₁₀ allele age, brain–blood specificity and Voight-correct |iHS| with full covariance and 10 random initialisations per k. Section B: feature medians per cluster across all 14 annotated features. Section C: cluster centroids in unscaled feature space and the corresponding LDSC EUR baseline annotation counts. Section D: top 15 genes per cluster ordered by variant count; 'no gene assignment' indicates intergenic credible-set variants. Section E: within-cluster age × selection feature partial rank correlations (no MAF residualisation; descriptive).*

**Supplementary Table 5f | Comprehensive GMM model-selection battery and stability analysis.**

To rigorously justify the *k* = 3 component selection, we computed six complementary model-selection metrics over candidate *k* = 1–8 in the primary 3D feature space (log₁₀ allele age × brain–blood specificity × |iHS|, no MAPT, *n* = 4,918) and the 2D sensitivity feature space (log₁₀ allele age × |iHS|, no MAPT, *n* = 15,739): classical likelihood-fit criteria (BIC, AIC), the clustering-specific Integrated Completed Likelihood (ICL = BIC + 2 × entropy of posterior cluster probabilities, which penalises overlapping components), and three classical cluster-validity indices (silhouette score, Calinski–Harabasz index, Davies–Bouldin index).

**Section A. Full battery — 3D feature space (log_age × brain_spec × |iHS|, n = 4,918)**

| ***k*** | **BIC ↓** | **AIC ↓** | **ICL ↓** | **Silhouette ↑** | **Calinski–Harabasz ↑** | **Davies–Bouldin ↓** |
| --- | --- | --- | --- | --- | --- | --- |
| 1 | 41,875 | 41,817 | 41,875 | — | — | — |
| 2 | 39,466 | 39,343 | 39,879 | 0.319 | 1,899 | 1.353 |
| **3** | **38,321** | **38,132** | **39,870** ★ | **0.321** ★ | 1,642 | **1.164** ★ |
| 4 | 38,152 | 37,898 | 40,643 | 0.316 | 1,261 | 1.466 |
| 5 | 38,074 | 37,756 | 42,547 | 0.203 | 1,244 | 1.285 |
| 6 | 37,773 | 37,389 | 43,870 | 0.185 | 1,049 | 1.931 |
| 7 | 37,565 | 37,117 | 43,226 | 0.185 | 1,185 | 1.347 |
| 8 | 37,496 | 36,982 | 42,613 | 0.214 | 1,330 | 1.187 |

★ = optimal *k* by metric. ICL, silhouette and Davies–Bouldin all identify *k* = 3 as optimal. Calinski–Harabasz favours *k* = 2 by a small margin (1,899 vs 1,642 at *k* = 3). BIC and AIC continue to decrease monotonically with increasing *k*, a known over-fitting tendency of likelihood-fit criteria in low-sample Gaussian mixture estimation.

**Section B. Full battery — 2D sensitivity feature space (log_age × |iHS|, n = 15,739)**

| ***k*** | **BIC ↓** | **AIC ↓** | **ICL ↓** | **Silhouette ↑** | **Calinski–Harabasz ↑** | **Davies–Bouldin ↓** |
| --- | --- | --- | --- | --- | --- | --- |
| 1 | 89,347 | 89,308 | 89,347 | — | — | — |
| **2** | 82,599 | 82,515 | **84,470** ★ | 0.435 | 10,982 | 0.973 |
| 3 | 80,001 | 79,871 | 88,957 | **0.457** ★ | 13,092 | 0.816 |
| 4 | 79,038 | 78,862 | 91,586 | 0.437 | 13,259 | **0.808** ★ |
| 5 | 78,336 | 78,113 | 93,220 | 0.404 | **13,795** ★ | 0.820 |
| 6 | 77,621 | 77,353 | 96,037 | 0.339 | 11,310 | 0.941 |
| 7 | **77,271** | 76,957 | 96,395 | 0.318 | 11,456 | 0.970 |
| 8 | 77,283 | **76,923** | 95,080 | 0.312 | 11,006 | 0.999 |

In the 2D feature space, the metrics disagree on optimal *k* (ICL: 2; silhouette: 3; CH: 5; DB: 4; BIC: 7; AIC: 8), reflecting weaker discrete cluster structure in the 2D feature space than in 3D. Note that in absolute terms the 2D feature space yields better silhouette (0.457 at *k* = 3 vs 0.321 at 3D *k* = 3) and better Davies–Bouldin (0.816 vs 1.164) — i.e., the 2D clusters are more cleanly separated as pure pattern-recognition clusters. However, these clusters do not concentrate SCZ heritability (Supplementary Table 14): cluster *quality* and cluster *biological informativeness* are distinct properties, and the 3D feature space — although yielding mathematically less crisp clusters — produces clusters that map onto SCZ heritability concentration.

**Section C. Cluster-membership stability under bootstrap resampling (k = 3)**

For each feature space we generated 1,000 bootstrap re-samples of the GMM input data (sample with replacement to original *n*), refit the GMM at *k* = 3 with `n_init = 5` and a different `random_state` per bootstrap, predicted cluster labels for the full original sample using the bootstrap-fitted model, and computed the Adjusted Rand Index (ARI) of bootstrap cluster assignments against the reference (random_state = 42) cluster assignments.

| **Feature space** | **Mean ARI** | **SD** | **Min** | **Max** | **95% percentile CI** | **% bootstraps ARI > 0.8** | **% bootstraps ARI > 0.9** |
| --- | --- | --- | --- | --- | --- | --- | --- |
| **3D primary** | **0.967** | 0.019 | 0.899 | 0.994 | [0.919, 0.990] | **1,000 / 1,000** | 999 / 1,000 |
| 2D sensitivity | 0.978 | 0.010 | 0.936 | 0.996 | [0.956, 0.993] | 1,000 / 1,000 | 1,000 / 1,000 |

Both feature spaces yield extremely stable cluster assignments at *k* = 3. In the 3D primary feature space, 1,000 of 1,000 bootstrap samples yielded ARI > 0.8 and 999 of 1,000 yielded ARI > 0.9; in the 2D sensitivity feature space, 1,000 of 1,000 yielded ARI > 0.9. The cluster identification is therefore robust to sample-level perturbation and not an artefact of the single random initialisation used for the primary fit.

*Model-selection criteria (BIC, AIC, ICL, silhouette score, Calinski–Harabasz index, Davies–Bouldin index) for Gaussian mixture models with k = 1–8 in the three-dimensional (n = 4,918 variants) and two-dimensional (n = 15,739) feature spaces (17q21.31 MAPT excluded), and cluster-membership stability quantified as the adjusted Rand index (ARI) across 1,000 bootstrap resamples at k = 3. BIC, Bayesian information criterion; AIC, Akaike information criterion; ICL, integrated completed likelihood; ARI, adjusted Rand index.*

**Supplementary Table 6 | Sensitivity to long-range linkage disequilibrium region exclusion: Price et al. 2008 region list and partitioned heritability comparison.**

Variants masked from each cluster annotation under the 24 Price 2008 long-range LD regions (canonical-corrected to GRCh37), and the resulting shift in partitioned heritability enrichment.

**Section A. Price et al. 2008 long-range LD region list (GRCh37, 24 regions; canonical-corrected)**

| **Chromosome** | **Start (bp, GRCh37)** | **End (bp, GRCh37)** | **Cytoband / region label** | **Length (Mb)** |
| --- | --- | --- | --- | --- |
| 1 | 48,000,000 | 52,000,000 | 1p13.3 | 4.0 |
| 2 | 86,000,000 | 100,500,000 | 2p13 | 14.5 |
| 2 | 134,500,000 | 138,000,000 | 2q13 | 3.5 |
| 2 | 183,000,000 | 190,000,000 | 2q21 | 7.0 |
| 3 | 47,500,000 | 50,000,000 | 3p21.31 | 2.5 |
| 3 | 83,500,000 | 87,000,000 | 3p11 | 3.5 |
| 3 | 89,000,000 | 97,500,000 | 3p11/q12 | 8.5 |
| 5 | 45,500,000 | 50,500,000 | 5q11 (canonical-corrected from 44.5–50.5 Mb) | 5.0 |
| 5 | 98,000,000 | 100,500,000 | 5q14 | 2.5 |
| 5 | 129,000,000 | 132,000,000 | 5q23 | 3.0 |
| 5 | 135,500,000 | 138,500,000 | 5q31 | 3.0 |
| 6 | 25,500,000 | 33,500,000 | 6p21 HLA (canonical-corrected from 25–35 Mb) | 8.0 |
| 6 | 57,000,000 | 64,000,000 | 6p11 | 7.0 |
| 6 | 140,000,000 | 142,500,000 | 6q23 | 2.5 |
| 7 | 55,000,000 | 66,000,000 | 7p11 | 11.0 |
| 8 | 8,000,000 | 12,000,000 | 8p23 (canonical-corrected from 7–13 Mb) | 4.0 |
| 8 | 43,000,000 | 50,000,000 | 8p11 | 7.0 |
| 8 | 112,000,000 | 115,000,000 | 8q23 | 3.0 |
| 10 | 37,000,000 | 43,000,000 | 10p11 | 6.0 |
| 11 | 46,000,000 | 57,000,000 | 11p11 | 11.0 |
| 11 | 87,500,000 | 90,500,000 | 11q14 | 3.0 |
| 12 | 33,000,000 | 40,000,000 | 12p11 | 7.0 |
| 12 | 109,500,000 | 112,000,000 | 12q21 | 2.5 |
| 20 | 32,000,000 | 34,500,000 | 20p12 | 2.5 |

**Section B. Variants masked per cluster**

| **Cluster** | **n in EUR baseline annotation (pre-mask)** | **n masked by Price 2008 LR-LD regions** | **% masked** | **n in annotation (post-mask)** |
| --- | --- | --- | --- | --- |
| C0 (Young) | 1,744 | 66 | 3.8 | 1,678 |
| C1 (Mid) | 317 | 25 | 7.9 | 292 |
| C2 (Old) | 2,855 | 75 | 2.6 | 2,780 |

**Section C. SCZ EUR partitioned heritability: primary vs Price 2008 LR-LD-masked**

| **Cluster** | **Primary enrichment** | **Primary P** | **Primary coef. Z** | **LR-LD-masked enrichment** | **LR-LD-masked P** | **LR-LD-masked coef. Z** | **Δ enrichment** |
| --- | --- | --- | --- | --- | --- | --- | --- |
| C0 (Young) | 47.35× ± 14.86 | 2.0 × 10⁻³ | +3.05 | 47.91× ± 15.22 | 2.3 × 10⁻³ | +3.01 | +0.56× |
| C1 (Mid) | 13.04× ± 3.89 | 2.1 × 10⁻³ | +3.32 | 12.06× ± 3.59 | 2.2 × 10⁻³ | +3.32 | −0.98× |
| C2 (Old) | 15.61× ± 12.89 | 0.259 | +1.04 | 16.02× ± 13.01 | 0.250 | +1.06 | +0.41× |

**Section D. Wightman 2021 AD partitioned heritability: primary vs Price 2008 LR-LD-masked**

| **Cluster** | **Primary enrichment** | **Primary P** | **LR-LD-masked enrichment** | **LR-LD-masked P** | **Δ enrichment** |
| --- | --- | --- | --- | --- | --- |
| C0 (Young) | −4.07× ± 8.81 | 0.564 | −4.67× ± 9.27 | 0.541 | −0.60× |
| C1 (Mid) | 4.01× ± 3.71 | 0.380 | 3.77× ± 3.47 | 0.390 | −0.24× |
| C2 (Old) | −4.88× ± 8.98 | 0.485 | −5.13× ± 9.00 | 0.468 | −0.25× |

*Section A: 24 long-range LD regions from Price et al. 2008²⁵ (their Table 1) in GRCh37 coordinates with three canonical-corrected boundaries documented in Methods. Section B: variants masked per cluster. Section C: SCZ EUR partitioned heritability comparison (PGC3 EUR¹). Section D: Wightman 2021 AD partitioned heritability comparison¹⁹. All regressions condition on the 97-annotation baseline-LD v2.2 model²⁴. Δ enrichment is the LR-LD-masked minus primary value. The LR-LD-masked results recapitulate the primary findings within Δ < 2× absolute for all six SCZ + AD comparisons.*

**Supplementary Table 7 | Single-cell brain cell-type cis-eQTL coverage by cluster and within-locus age × cell-type cis-eQTL strength correlations.**

Section A: proportion of cluster variants with at least one significant cis-eQTL in each Bryois 2022 cell type, with three-way χ² test. Section B: within-locus partial rank correlation between log₁₀ allele age and per-cell-type minimum cis-eQTL −log₁₀ P.

**Section A. Cluster × cell-type cis-eQTL coverage and three-way χ² P**

| **Cell type (Bryois 2022⁹)** | **C0 (Young) coverage (%)** | **C1 (Mid) coverage (%)** | **C2 (Old) coverage (%)** | **C0–C2 difference (pp)** | **C0–C1 difference (pp)** | **Three-way χ² statistic** | **P (χ², 2 d.f.)** | **BH-FDR q** |
| --- | --- | --- | --- | --- | --- | --- | --- | --- |
| Astrocytes | 94.10 | 77.92 | 93.52 | +0.58 | +16.18 | 110.53 | 1.0 × 10⁻²⁴ | 2.7 × 10⁻²⁴ |
| Endothelial cells | 95.07 | 77.92 | 93.59 | +1.48 | +17.15 | 125.50 | 5.6 × 10⁻²⁸ | 4.5 × 10⁻²⁷ |
| Excitatory neurons | 98.28 | 94.01 | 98.18 | +0.10 | +4.27 | 26.21 | 2.0 × 10⁻⁶ | 2.3 × 10⁻⁶ |
| Inhibitory neurons | 98.28 | 94.01 | 98.18 | +0.10 | +4.27 | 26.21 | 2.0 × 10⁻⁶ | 2.3 × 10⁻⁶ |
| Microglia | 96.91 | 85.17 | 93.80 | +3.10 | +11.73 | 73.02 | 1.4 × 10⁻¹⁶ | 2.8 × 10⁻¹⁶ |
| OPCs and COPs | 98.28 | 94.01 | 98.00 | +0.28 | +4.27 | 24.06 | 5.9 × 10⁻⁶ | 5.9 × 10⁻⁶ |
| Oligodendrocytes | 94.56 | 77.92 | 93.66 | +0.89 | +16.64 | 118.61 | 1.8 × 10⁻²⁶ | 7.0 × 10⁻²⁶ |
| Pericytes | 94.67 | 82.33 | 93.14 | +1.53 | +12.34 | 62.83 | 2.3 × 10⁻¹⁴ | 3.6 × 10⁻¹⁴ |

**Section B. Within-locus partial rank correlations: log10 allele age × cell-type cis-eQTL strength**

| **Cell type (Bryois 2022⁹)** | **n full (variant-pooled, all loci)** | **ρ full** | **P full** | **n no MAPT (within-locus, MAF-residualised)** | **ρ no MAPT** | **P no MAPT** | **BH-FDR q (no MAPT)** |
| --- | --- | --- | --- | --- | --- | --- | --- |
| Astrocytes | 16,801 | +0.063 | 3.8 × 10⁻¹⁶ | 15,260 | −0.096 | 2.4 × 10⁻³² | 6.4 × 10⁻³² |
| Endothelial cells | 15,078 | +0.166 | 1.4 × 10⁻⁹³ | 13,537 | −0.002 | 0.833 | 0.833 |
| Excitatory neurons | 18,276 | +0.146 | 3.2 × 10⁻⁸⁷ | 16,735 | +0.080 | 5.0 × 10⁻²⁵ | 1.0 × 10⁻²⁴ |
| Inhibitory neurons | 18,241 | +0.186 | 3.1 × 10⁻¹⁴¹ | 16,700 | +0.092 | 2.2 × 10⁻³² | 6.4 × 10⁻³² |
| Microglia | 14,595 | +0.187 | 2.0 × 10⁻¹¹⁴ | 13,054 | +0.044 | 4.9 × 10⁻⁷ | 7.8 × 10⁻⁷ |
| OPCs and COPs | 17,834 | +0.100 | 2.9 × 10⁻⁴¹ | 16,293 | −0.017 | 0.027 | 0.036 |
| Oligodendrocytes | 16,239 | +0.097 | 2.4 × 10⁻³⁵ | 14,698 | −0.123 | 2.0 × 10⁻⁵⁰ | 1.6 × 10⁻⁴⁹ |
| Pericytes | 13,958 | +0.209 | 1.2 × 10⁻¹³⁷ | 12,417 | −0.014 | 0.127 | 0.145 |

*Bryois et al. 2022 cell types⁹: astrocytes, endothelial cells, excitatory neurons, inhibitory neurons, microglia, OPCs and COPs, oligodendrocytes, pericytes. Section A χ² is a three-way Pearson χ² with 2 degrees of freedom. Section B is computed on within-locus rank residuals after MAF-rank residualisation (no MAPT). The full specification (variant-pooled) is provided for sensitivity. Three-way χ² P-values are dominated by depressed C1 (Mid) cluster engagement rather than by elevated C0 (Young) cluster specificity (manuscript Results, Fig. 3).*

**Supplementary Table 8 | Cluster-specific pathway enrichment and brain region distribution.**

Section A: top 5 pathway enrichments per Enrichr library (GO Biological Process 2023, KEGG 2021 Human, Reactome 2022, MSigDB Hallmark 2020) per cluster. Section B: cluster variant counts at the GTEx v10 brain tissue with strongest cis-eQTL.

**Section A. Cluster-specific pathway enrichment (top 5 per library × cluster; Enrichr 2026)**

| **Cluster** | **Library** | **Term** | **Overlap (genes/total)** | **Genes** | **Enrichr P** | **BH-adjusted P** |
| --- | --- | --- | --- | --- | --- | --- |
| C0 (Young) | GO_Biological_Process_2023 | Dendrite Arborization (GO:0140059) | 2/6 | TAOK2;ZNF365 | 0.001 | 0.447 |
| C0 (Young) | GO_Biological_Process_2023 | Positive Regulation Of Protein Dephosphorylation (GO:0035307) | 3/33 | PPP1R16B;AMBRA1;SPPL3 | 0.004 | 0.447 |
| C0 (Young) | GO_Biological_Process_2023 | Positive Regulation Of Mitotic Cell Cycle Spindle Assembly Checkpoint (GO:0090267) | 2/11 | XRCC3;MAD1L1 | 0.005 | 0.447 |
| C0 (Young) | GO_Biological_Process_2023 | Positive Regulation Of Spindle Checkpoint (GO:0090232) | 2/11 | XRCC3;MAD1L1 | 0.005 | 0.447 |
| C0 (Young) | GO_Biological_Process_2023 | Interstrand Cross-Link Repair (GO:0036297) | 3/36 | FANCL;XRCC3;FANCA | 0.006 | 0.447 |
| C0 (Young) | KEGG_2021_Human | Parathyroid hormone synthesis, secretion and action | 4/106 | MMP16;CREB3L4;FGFR1;GNAI2 | 0.022 | 0.807 |
| C0 (Young) | KEGG_2021_Human | Insulin resistance | 4/108 | MLXIP;CRTC2;CREB3L4;PTPRF | 0.023 | 0.807 |
| C0 (Young) | KEGG_2021_Human | Cholinergic synapse | 4/113 | CHRNA3;CREB3L4;CACNA1D;GNAI2 | 0.027 | 0.807 |
| C0 (Young) | KEGG_2021_Human | Adherens junction | 3/71 | CTNND1;PTPRF;FGFR1 | 0.034 | 0.807 |
| C0 (Young) | KEGG_2021_Human | Glyoxylate and dicarboxylate metabolism | 2/30 | PCCB;HYI | 0.036 | 0.807 |
| C0 (Young) | Reactome_2022 | Highly Calcium Permeable Nicotinic Acetylcholine Receptors R-HSA-629597 | 3/9 | CHRNA3;CHRNA2;CHRNA5 | 7.9 × 10⁻⁵ | 0.030 |
| C0 (Young) | Reactome_2022 | Highly Calcium Permeable Postsynaptic Nicotinic Acetylcholine Receptors R-HSA-629594 | 3/11 | CHRNA3;CHRNA2;CHRNA5 | 1.5 × 10⁻⁴ | 0.030 |
| C0 (Young) | Reactome_2022 | Presynaptic Nicotinic Acetylcholine Receptors R-HSA-622323 | 3/12 | CHRNA3;CHRNA2;CHRNA5 | 2.0 × 10⁻⁴ | 0.030 |
| C0 (Young) | Reactome_2022 | Acetylcholine Binding And Downstream Events R-HSA-181431 | 3/14 | CHRNA3;CHRNA2;CHRNA5 | 3.3 × 10⁻⁴ | 0.037 |
| C0 (Young) | Reactome_2022 | TP53 Regulates Transcription Of Death Receptors And Ligands R-HSA-6803211 | 2/12 | PPP1R13B;TMEM219 | 0.006 | 0.545 |
| C0 (Young) | MSigDB_Hallmark_2020 | Apical Junction | 5/200 | LIMA1;TAOK2;CTNND1;EXOC4;GNAI2 | 0.051 | 0.867 |
| C0 (Young) | MSigDB_Hallmark_2020 | Heme Metabolism | 4/200 | BNIP3L;SLC6A9;CLCN3;MARK3 | 0.141 | 0.867 |
| C0 (Young) | MSigDB_Hallmark_2020 | Apoptosis | 3/161 | BNIP3L;ETF1;CLU | 0.218 | 0.867 |
| C0 (Young) | MSigDB_Hallmark_2020 | Protein Secretion | 2/96 | VPS45;CLCN3 | 0.249 | 0.867 |
| C0 (Young) | MSigDB_Hallmark_2020 | Angiogenesis | 1/36 | FGFR1 | 0.304 | 0.867 |
| C1 (Mid) | GO_Biological_Process_2023 | Regulation Of Cell Fate Commitment (GO:0010453) | 2/13 | GATAD2B;GATAD2A | 3.2 × 10⁻⁴ | 0.052 |
| C1 (Mid) | GO_Biological_Process_2023 | Regulation Of Cell Fate Specification (GO:0042659) | 2/14 | GATAD2B;GATAD2A | 3.7 × 10⁻⁴ | 0.052 |
| C1 (Mid) | GO_Biological_Process_2023 | Positive Regulation Of Protein Targeting To Mitochondrion (GO:1903955) | 2/30 | UBE2D3;ATG13 | 0.002 | 0.116 |
| C1 (Mid) | GO_Biological_Process_2023 | Positive Regulation Of Establishment Of Protein Localization To Mitochondrion (GO:1903749) | 2/33 | UBE2D3;ATG13 | 0.002 | 0.116 |
| C1 (Mid) | GO_Biological_Process_2023 | Histone Deacetylation (GO:0016575) | 2/36 | GATAD2B;GATAD2A | 0.002 | 0.116 |
| C1 (Mid) | KEGG_2021_Human | Ubiquitin mediated proteolysis | 2/140 | CDC20;UBE2D3 | 0.033 | 0.405 |
| C1 (Mid) | KEGG_2021_Human | Primary bile acid biosynthesis | 1/17 | CYP7B1 | 0.034 | 0.405 |
| C1 (Mid) | KEGG_2021_Human | Fatty acid elongation | 1/27 | ELOVL1 | 0.054 | 0.405 |
| C1 (Mid) | KEGG_2021_Human | Biosynthesis of unsaturated fatty acids | 1/27 | ELOVL1 | 0.054 | 0.405 |
| C1 (Mid) | KEGG_2021_Human | Fanconi anemia pathway | 1/54 | FANCA | 0.105 | 0.405 |
| C1 (Mid) | Reactome_2022 | Regulation Of TP53 Activity Thru Acetylation R-HSA-6804758 | 2/29 | GATAD2B;GATAD2A | 0.002 | 0.152 |
| C1 (Mid) | Reactome_2022 | ERCC6 (CSB) And EHMT2 (G9a) Positively Regulate rRNA Expression R-HSA-427389 | 2/45 | GATAD2B;GATAD2A | 0.004 | 0.152 |
| C1 (Mid) | Reactome_2022 | RNA Polymerase I Transcription Initiation R-HSA-73762 | 2/46 | GATAD2B;GATAD2A | 0.004 | 0.152 |
| C1 (Mid) | Reactome_2022 | Regulation Of TP53 Activity R-HSA-5633007 | 3/157 | PPP1R13B;GATAD2B;GATAD2A | 0.004 | 0.152 |
| C1 (Mid) | Reactome_2022 | HDACs Deacetylate Histones R-HSA-3214815 | 2/60 | GATAD2B;GATAD2A | 0.007 | 0.152 |
| C1 (Mid) | MSigDB_Hallmark_2020 | Bile Acid Metabolism | 2/112 | EPHX2;CYP7B1 | 0.022 | 0.265 |
| C1 (Mid) | MSigDB_Hallmark_2020 | E2F Targets | 2/200 | CDC20;CENPM | 0.063 | 0.338 |
| C1 (Mid) | MSigDB_Hallmark_2020 | TGF-beta Signaling | 1/54 | UBE2D3 | 0.105 | 0.338 |
| C1 (Mid) | MSigDB_Hallmark_2020 | Peroxisome | 1/104 | EPHX2 | 0.193 | 0.338 |
| C1 (Mid) | MSigDB_Hallmark_2020 | PI3K/AKT/mTOR Signaling | 1/105 | UBE2D3 | 0.194 | 0.338 |
| C2 (Old) | GO_Biological_Process_2023 | Signal Peptide Processing (GO:0006465) | 3/13 | FURIN;SPPL3;SEC11A | 2.6 × 10⁻⁴ | 0.303 |
| C2 (Old) | GO_Biological_Process_2023 | Positive Regulation Of CREB Transcription Factor Activity (GO:0032793) | 3/18 | OPRD1;CRTC2;TSSK6 | 7.2 × 10⁻⁴ | 0.337 |
| C2 (Old) | GO_Biological_Process_2023 | Phosphorylation (GO:0016310) | 12/429 | SRPK2;NEK4;TAF12;TAOK2;PAK6;TSSK6;PTK2B;ATG13;DGKZ;MARK3;CAMKK2;FGFR1 | 0.001 | 0.337 |
| C2 (Old) | GO_Biological_Process_2023 | Clathrin-Coated Vesicle Cargo Loading (GO:0035652) | 2/6 | AP3D1;AP3B2 | 0.001 | 0.337 |
| C2 (Old) | GO_Biological_Process_2023 | Clathrin-Coated Vesicle Cargo Loading, AP-3-mediated (GO:0035654) | 2/6 | AP3D1;AP3B2 | 0.001 | 0.337 |
| C2 (Old) | KEGG_2021_Human | Fanconi anemia pathway | 3/54 | FANCI;FANCL;FANCA | 0.017 | 0.910 |
| C2 (Old) | KEGG_2021_Human | Various types of N-glycan biosynthesis | 2/39 | MAN2A1;ST3GAL3 | 0.058 | 0.910 |
| C2 (Old) | KEGG_2021_Human | Other types of O-glycan biosynthesis | 2/47 | GALNT10;ST3GAL3 | 0.080 | 0.910 |
| C2 (Old) | KEGG_2021_Human | Insulin resistance | 3/108 | MLXIP;CRTC2;PTPRF | 0.094 | 0.910 |
| C2 (Old) | KEGG_2021_Human | Cholinergic synapse | 3/113 | CHRNB4;CHRNA3;CACNA1D | 0.104 | 0.910 |
| C2 (Old) | Reactome_2022 | Highly Calcium Permeable Nicotinic Acetylcholine Receptors R-HSA-629597 | 3/9 | CHRNA3;CHRNB4;CHRNA2 | 7.9 × 10⁻⁵ | 0.023 |
| C2 (Old) | Reactome_2022 | Chromatin Modifying Enzymes R-HSA-3247509 | 10/238 | SMARCD1;KDM4A;PBRM1;KDM3B;TAF12;ATXN7;SETD6;DOT1L;GATAD2B;GATAD2A | 1.5 × 10⁻⁴ | 0.023 |
| C2 (Old) | Reactome_2022 | Highly Calcium Permeable Postsynaptic Nicotinic Acetylcholine Receptors R-HSA-629594 | 3/11 | CHRNA3;CHRNB4;CHRNA2 | 1.5 × 10⁻⁴ | 0.023 |
| C2 (Old) | Reactome_2022 | Presynaptic Nicotinic Acetylcholine Receptors R-HSA-622323 | 3/12 | CHRNA3;CHRNB4;CHRNA2 | 2.0 × 10⁻⁴ | 0.023 |
| C2 (Old) | Reactome_2022 | Acetylcholine Binding And Downstream Events R-HSA-181431 | 3/14 | CHRNA3;CHRNB4;CHRNA2 | 3.3 × 10⁻⁴ | 0.031 |
| C2 (Old) | MSigDB_Hallmark_2020 | TGF-beta Signaling | 2/54 | UBE2D3;FURIN | 0.102 | 0.867 |
| C2 (Old) | MSigDB_Hallmark_2020 | mTORC1 Signaling | 4/200 | TXNRD1;UBE2D3;SEC11A;HSPD1 | 0.141 | 0.867 |
| C2 (Old) | MSigDB_Hallmark_2020 | Androgen Response | 2/100 | PTK2B;CAMKK2 | 0.264 | 0.867 |
| C2 (Old) | MSigDB_Hallmark_2020 | Angiogenesis | 1/36 | FGFR1 | 0.304 | 0.867 |
| C2 (Old) | MSigDB_Hallmark_2020 | Bile Acid Metabolism | 2/112 | EPHX2;CYP7B1 | 0.309 | 0.867 |

**Section B. Brain region distribution: cluster variant counts at strongest cis-eQTL tissue (GTEx v10)**

| **GTEx brain tissue (best cis-eQTL)** | **C0 (Young)** | **C1 (Mid)** | **C2 (Old)** |
| --- | --- | --- | --- |
| Brain_Amygdala | 36 | 3 | 21 |
| Brain_Anterior_cingulate_cortex_BA24 | 22 | 0 | 11 |
| Brain_Caudate_basal_ganglia | 242 | 30 | 197 |
| Brain_Cerebellar_Hemisphere | 214 | 129 | 571 |
| Brain_Cerebellum | 351 | 39 | 851 |
| Brain_Cortex | 165 | 39 | 417 |
| Brain_Frontal_Cortex_BA9 | 155 | 12 | 265 |
| Brain_Hippocampus | 129 | 28 | 94 |
| Brain_Hypothalamus | 2 | 0 | 28 |
| Brain_Nucleus_accumbens_basal_ganglia | 215 | 8 | 91 |
| Brain_Putamen_basal_ganglia | 25 | 0 | 96 |
| Brain_Spinal_cord_cervical_c-1 | 182 | 29 | 213 |
| Brain_Substantia_nigra | 7 | 0 | 1 |

*Pathway enrichment was computed against the credible-set gene universe using Enrichr-style hypergeometric tests; the BH-adjusted P-value (Adjusted P) is the per-library Benjamini-Hochberg q-value. Genes are listed semicolon-separated per term. All three clusters enrich for shared schizophrenia pathology pathways (e.g. nicotinic acetylcholine receptor signalling), and cluster differentiation in pathway space is largely null; cluster differentiation is confined to brain region of strongest cis-eQTL (Section B), broad cell-type usage (Supplementary Table 7), and gene-level constraint (Supplementary Table 10).*

**Supplementary Table 9 | Per-tissue brain and blood cis-eQTL × log₁₀ allele age correlations across GTEx v10.**

Within-locus partial rank correlation of variant-level minimum cis-eQTL strength against log₁₀ allele age, in 13 brain tissues plus whole blood and spleen, with and without the 17q21.31 MAPT inversion.

| **GTEx v10 tissue** | **n full** | **ρ full (variant-pooled)** | **P full** | **n no-MAPT** | **ρ no-MAPT (within-locus rank, MAF-residualised)** | **P no-MAPT** | **BH-FDR q (full)** | **BH-FDR q (no-MAPT)** |
| --- | --- | --- | --- | --- | --- | --- | --- | --- |
| Brain – Hypothalamus | 5,120 | +0.190 | 7.5 × 10⁻⁴³ | 3,446 | +0.076 | 8.4 × 10⁻⁶ | 3.8 × 10⁻⁴² | 1.7 × 10⁻⁵ |
| Brain – Hippocampus | 6,011 | +0.189 | 2.4 × 10⁻⁴⁹ | 4,337 | +0.067 | 9.1 × 10⁻⁶ | 3.6 × 10⁻⁴⁸ | 1.7 × 10⁻⁵ |
| Brain – Anterior cingulate cortex (BA24) | 5,362 | +0.181 | 8.6 × 10⁻⁴¹ | 3,688 | +0.126 | 1.9 × 10⁻¹⁴ | 2.1 × 10⁻⁴⁰ | 2.8 × 10⁻¹³ |
| Whole blood | 7,794 | +0.164 | 5.5 × 10⁻⁴⁸ | 6,120 | +0.065 | 3.5 × 10⁻⁷ | 4.1 × 10⁻⁴⁷ | 1.1 × 10⁻⁶ |
| Brain – Caudate (basal ganglia) | 7,027 | +0.162 | 2.0 × 10⁻⁴² | 5,353 | +0.082 | 2.2 × 10⁻⁹ | 7.7 × 10⁻⁴² | 8.3 × 10⁻⁹ |
| Brain – Amygdala | 3,898 | +0.161 | 4.9 × 10⁻²⁴ | 2,224 | +0.082 | 1.0 × 10⁻⁴ | 6.6 × 10⁻²⁴ | 1.5 × 10⁻⁴ |
| Brain – Cerebellar hemisphere | 6,838 | +0.161 | 9.7 × 10⁻⁴¹ | 5,164 | +0.059 | 2.2 × 10⁻⁵ | 2.1 × 10⁻⁴⁰ | 3.6 × 10⁻⁵ |
| Brain – Cerebellum | 7,091 | +0.161 | 3.9 × 10⁻⁴² | 5,417 | +0.082 | 1.3 × 10⁻⁹ | 1.2 × 10⁻⁴¹ | 6.6 × 10⁻⁹ |
| Brain – Putamen (basal ganglia) | 5,841 | +0.154 | 2.4 × 10⁻³² | 4,167 | +0.001 | 0.950 | 3.6 × 10⁻³² | 0.950 |
| Brain – Cortex | 6,944 | +0.148 | 3.9 × 10⁻³⁵ | 5,270 | +0.090 | 7.5 × 10⁻¹¹ | 7.4 × 10⁻³⁵ | 5.7 × 10⁻¹⁰ |
| Brain – Frontal cortex (BA9) | 6,580 | +0.147 | 2.7 × 10⁻³³ | 4,906 | +0.030 | 0.034 | 4.6 × 10⁻³³ | 0.042 |
| Brain – Substantia nigra | 3,682 | +0.142 | 4.5 × 10⁻¹⁸ | 2,008 | +0.111 | 6.5 × 10⁻⁷ | 4.8 × 10⁻¹⁸ | 1.6 × 10⁻⁶ |
| Brain – Spinal cord (cervical c-1) | 5,131 | +0.121 | 4.2 × 10⁻¹⁸ | 3,457 | −0.003 | 0.860 | 4.8 × 10⁻¹⁸ | 0.922 |
| Spleen | 6,456 | +0.113 | 6.1 × 10⁻²⁰ | 4,782 | +0.018 | 0.215 | 7.7 × 10⁻²⁰ | 0.248 |
| Brain – Nucleus accumbens (basal ganglia) | 6,000 | +0.083 | 1.1 × 10⁻¹⁰ | 4,326 | −0.038 | 0.012 | 1.1 × 10⁻¹⁰ | 0.017 |

*ρ full is the variant-pooled rank correlation including all credible-set variants in the tissue. ρ no-MAPT is the within-locus partial rank correlation after MAF-rank residualisation, excluding the 17q21.31 MAPT inversion (CS_224). The no-MAPT specification is the primary specification used throughout the main text; the full specification is provided for sensitivity. Tissues are sorted by descending |ρ full|.*

**Supplementary Table 10 | Within-locus partial rank correlations between gene-level constraint, allele age and brain cis-eQTL strength.**

Within-credible-set partial rank correlations between gnomAD v4 gene-level constraint metrics, log₁₀ allele age, and brain cis-eQTL strength.

| **Test** | **Specification** | **n variants** | **n loci** | **Spearman ρ** | **Asymptotic P** | **BH-FDR q** | **Source / notes** |
| --- | --- | --- | --- | --- | --- | --- | --- |
| LOEUF × log₁₀ allele age | within-locus rank residualisation | 6,526 | 73 | −0.184 | 1.3 × 10⁻⁵⁰ | 3.4 × 10⁻⁵⁰ | gnomAD v4 LOEUF¹³,³⁴; higher LOEUF means lower constraint |
| Brain min cis-eQTL −log₁₀ P × LOEUF | within-locus rank residualisation | 5,393 | 64 | +0.451 | 1.2 × 10⁻²⁶⁸ | 1.2 × 10⁻²⁶⁷ | GTEx v10 brain × gnomAD v4 LOEUF; positive ρ indicates stronger eQTLs at less-constrained genes |
| Brain–blood specificity × LOEUF | within-locus rank residualisation | 4,601 | 53 | +0.342 | 4.9 × 10⁻¹²⁶ | 2.5 × 10⁻¹²⁵ | Brain-restricted cis-regulation × gene constraint |
| pLI × log₁₀ allele age | within-locus rank + MAF-rank residualisation, no MAPT | 5,637 | 72 | +0.081 | 1.3 × 10⁻⁹ | n/a (descriptive) | gnomAD v4 pLI; higher pLI = stronger LoF constraint |
| Missense Z × log₁₀ allele age | within-locus rank + MAF-rank residualisation, no MAPT | 5,664 | 72 | +0.182 | 1.3 × 10⁻⁴³ | n/a (descriptive) | gnomAD v4 missense constraint Z |
| LoF Z × log₁₀ allele age | within-locus rank + MAF-rank residualisation, no MAPT | 5,637 | 72 | +0.084 | 2.1 × 10⁻¹⁰ | n/a (descriptive) | gnomAD v4 LoF constraint Z |

*Constraint metrics are gnomAD v4 LOEUF, pLI, missense Z and LoF Z¹³,³⁴, mapped via the nearest-gene assignment of each credible-set variant. The within-locus rank specification is the primary; MAF-rank residualisation is added where the n is large enough (no MAPT). LOEUF is the loss-of-function observed/expected upper-bound fraction; lower LOEUF means stronger constraint. The negative LOEUF × age correlation (ρ = −0.184, P = 1.3 × 10⁻⁵⁰) recapitulates established gene-constraint signals¹³,³⁴ at the variant level within disease-specific credible sets. Supports the within-locus correlations in Fig. 4.*

**Supplementary Table 11 | Full cross-disorder partitioned LD score regression: C0, C1 and C2 cluster enrichments across six psychiatric comparator sumstats.**

All regressions condition on the 97-annotation baseline-LD v2.2 model²⁴ over the EUR LDSC reference. The C0/C1/C2 cluster annotations are identical to those used for the PGC3 EUR SCZ discovery analysis (Methods, manuscript main text, Table 2). The disorder-level Bonferroni threshold across six comparators is α = 0.05 / 6 = 0.0083; the focal-annotation significance threshold for the conditional coefficient *Z*-score is approximately 2.81 (Methods). Sample-overlap LDSC intercepts are reported in the last column to flag cohorts where shared PGC umbrella controls or UK Biobank ascertainment may inflate enrichment estimates.

| **Trait (sumstats)** | **Cluster** | **Prop. SNPs** | **Prop. h²** | **Prop. h² s.e.** | **Enrichment** | **s.e.** | ***P*** | **Coef. *Z*** | **LDSC intercept** |
| --- | --- | --- | --- | --- | --- | --- | --- | --- | --- |
| **PGC BD2024⁵⁹ EUR** (h² = 0.128; *n*<sub>cas</sub> = 59,287) | C0 | 0.000288 | 0.00686 | 0.0029 | 23.87× | 10.07 | 0.024 | +2.14 | 1.070† |
|  | C1 | 0.000261 | 0.000853 | 0.000653 | 3.27× | 2.51 | 0.364 | +1.19 | — |
|  | C2 | 0.000357 | 0.00346 | 0.00323 | 9.68× | 9.04 | 0.335 | +0.79 | — |
| **PGC MDD2025⁸³ EUR** (h² = 0.045; *n*<sub>cas</sub> = 412,305) | C0 | 0.000288 | 0.00349 | 0.00187 | 12.14× | 6.52 | 0.089 | +1.55 | 1.099† |
|  | C1 | 0.000261 | 0.000774 | 0.000323 | 2.97× | 1.24 | 0.114 | +2.22 | — |
|  | C2 | 0.000357 | −0.000908 | 0.00128 | −2.54× | 3.59 | 0.326 | −1.26 | — |
| **PGC ADHD2022⁵⁷** (h² = 0.170; *n*<sub>cas</sub> = 38,691) | C0 | 0.000288 | 0.00367 | 0.00492 | 12.77× | 17.10 | 0.491 | +0.62 | 1.036 |
|  | C1 | 0.000261 | −1 × 10⁻⁶ | 0.000346 | −0.01× | 1.33 | 0.451 | −0.14 | — |
|  | C2 | 0.000357 | 0.00852 | 0.00457 | 23.84× | 12.80 | 0.074 | +1.71 | — |
| **iPSYCH-PGC autism⁸¹** (h² = 0.199; *n*<sub>cas</sub> = 18,381) | C0 | 0.000288 | 0.00159 | 0.00364 | 5.54× | 12.67 | 0.719 | +0.27 | 1.009 |
|  | C1 | 0.000261 | 0.000686 | 0.000868 | 2.63× | 3.33 | 0.621 | +0.72 | — |
|  | C2 | 0.000357 | 0.0019 | 0.00544 | 5.32× | 15.24 | 0.777 | +0.21 | — |
| **CDG3 F3⁸⁴ Neurodevelopmental** (h² = 0.172; implied *N* = 84,760) | C0 | 0.000288 | 0.00148 | 0.00252 | 5.15× | 8.76 | 0.636 | +0.39 | 0.963 |
|  | C1 | 0.000261 | 0.000199 | 0.000543 | 0.77× | 2.08 | 0.911 | +0.26 | — |
|  | C2 | 0.000357 | 0.00676 | 0.00573 | 18.91× | 16.04 | 0.268 | +1.06 | — |
| **CDG3 F4⁸⁴ Internalising** (h² = 0.039; implied *N* = 1,637,337) | C0 | 0.000288 | 0.00408 | 0.0022 | 14.19× | 7.66 | 0.085 | +1.58 | 1.087† |
|  | C1 | 0.000261 | 0.000451 | 0.000207 | 1.73× | 0.80 | 0.360 | +1.87 | — |
|  | C2 | 0.000357 | −0.000246 | 0.00127 | −0.69× | 3.56 | 0.635 | −0.78 | — |

† LDSC intercept > 1.05 indicates moderate sample overlap with reference cohorts (PGC umbrella controls; UK Biobank ascertainment) and warrants cautious quantitative interpretation. The CDG3 F4 factor includes major depression as one of three constituent disorders (F4 = PTSD + MDD + anxiety), so the F4 enrichment is partly a recapitulation of the MDD enrichment rather than an independent observation.

*The bipolar disorder enrichment is the strongest cross-disorder signal and is nominally significant, but does not survive the disorder-level Bonferroni threshold (P = 0.024 > 0.0083); it is directionally consistent with the principal Young-cluster signal across the psychotic-spectrum architecture. The marginal MDD and CDG3 F4 enrichments (P = 0.089 and 0.085) also fail Bonferroni and are partly confounded by sample overlap. The ADHD null at matched cohort properties (n<sub>cas</sub> ≈ BD; intercept similar; SNP-h² similar) is the load-bearing evidence for a substantive psychotic-mood-vs-neurodevelopmental dissociation. The autism null is power-limited (mean χ² = 1.19) and is hypothesis-generating rather than confirmatory. The CDG3 F3 null partially recapitulates the ADHD null because ADHD is one of the three F3 constituent disorders.*

**Supplementary Table 12 | Full sex-stratified partitioned LD score regression: C0, C1 and C2 cluster enrichments across four PGC3 wave 3 sex × ancestry strata.**

All regressions condition on the 97-annotation baseline-LD v2.2 model per ref. 24. The cluster annotations are identical to those used for the corresponding full-sample SCZ EUR or SCZ EAS analyses (Methods, manuscript main text, Table 2). Effective sample size *N*<sub>eff</sub> = 4 · *N*<sub>cas</sub> · *N*<sub>con</sub> / (*N*<sub>cas</sub> + *N*<sub>con</sub>) was computed per row from the daner-format `Nca` and `Nco` columns. Sex-specificity of the C0 (Young) cluster enrichment was assessed by formal heterogeneity test *Z*<sub>diff</sub> = (enrichment<sub>male</sub> − enrichment<sub>female</sub>) / sqrt(s.e.²<sub>male</sub> + s.e.²<sub>female</sub>) under the assumption of independent stratum estimates.

| **Stratum** | **Cluster** | **Prop. SNPs** | **Prop. h²** | **Prop. h² s.e.** | **Enrichment** | **s.e.** | ***P*** | **Coef. *Z*** | **LDSC intercept** |
| --- | --- | --- | --- | --- | --- | --- | --- | --- | --- |
| **PGC3 EUR SCZ male¹** (*N*<sub>eff</sub> = 68,003) | C0 | 0.000288 | 0.0114 | 0.00384 | **39.62×** | 13.34 | **3.9 × 10⁻³** | **+2.84** | 1.023 |
|  | C1 | 0.000261 | 0.00362 | 0.000828 | **13.90×** | 3.18 | **7.5 × 10⁻⁵** | **+4.28** | — |
|  | C2 | 0.000357 | 0.00402 | 0.00373 | 11.26× | 10.45 | 0.326 | +0.88 | — |
| **PGC3 EUR SCZ female¹** (*N*<sub>eff</sub> = 47,652) | C0 | 0.000288 | 0.0156 | 0.00544 | **54.35×** | 18.93 | **4.9 × 10⁻³** | **+2.76** | 1.074† |
|  | C1 | 0.000261 | 0.00259 | 0.00166 | 9.92× | 6.38 | 0.160 | +1.53 | — |
|  | C2 | 0.000357 | 0.00968 | 0.00790 | 27.10× | 22.10 | 0.239 | +1.11 | — |
| **PGC3 EAS SCZ male¹** (*N*<sub>eff</sub> = 13,017) | C0 | 0.000279 | 0.013 | 0.0057 | **46.45×** | 20.41 | **0.023** | **+2.26** | 0.992 |
|  | C1 | 5.6 × 10⁻⁵ | −0.000116 | 0.00345 | −2.07× | 61.59 | 0.960 | −0.05 | — |
|  | C2 | 0.000438 | 0.00448 | 0.00372 | 10.21× | 8.49 | 0.271 | +1.06 | — |
| **PGC3 EAS SCZ female¹** (*N*<sub>eff</sub> = 13,163) | C0 | 0.000279 | 0.0128 | 0.00544 | **45.67×** | 19.50 | **0.029** | **+2.15** | 1.004 |
|  | C1 | 5.6 × 10⁻⁵ | −0.000514 | 0.00296 | −9.19× | 52.95 | 0.849 | −0.19 | — |
|  | C2 | 0.000438 | 0.0057 | 0.00384 | 13.02× | 8.76 | 0.156 | +1.36 | — |

† The EUR female stratum LDSC intercept of 1.074 indicates moderate sample overlap with reference cohorts, consistent with shared PGC umbrella controls present in the female-only daner. The other three strata have intercepts essentially at 1.0, consistent with negligible inflation.

**Formal heterogeneity tests for C0 (Young) cluster enrichment.** Under the assumption of independent stratum estimates, the EUR male-versus-female test is *Z*<sub>diff</sub> = (39.62 − 54.35) / sqrt(13.34² + 18.93²) = −0.64, *P* = 0.525 (two-sided). The EAS male-versus-female test is *Z*<sub>diff</sub> = (46.45 − 45.67) / sqrt(20.41² + 19.50²) = +0.03, *P* = 0.978 (two-sided). Both tests fail to reject sex-symmetry of the Young-cluster enrichment.

*All four strata exhibit significant C0 enrichment by single-test* P*-value, with point estimates within approximately half a standard error of the corresponding full-sample estimates (EUR 47.4×; EAS 54.6×). The EUR female cohort has 30% smaller effective sample size than the EUR male cohort (47,652 vs 68,003), corresponding to an approximately 23% larger female-stratum standard error; this asymmetry means a small sex-specific difference below the heterogeneity-test detection threshold cannot be excluded. C1 (Mid) enrichment is significant only in EUR male; this likely reflects power asymmetry given the small C1 cluster size (317 GMM-assigned variants). C2 (Old) enrichment is non-significant in all four strata.*

**Supplementary Table 13 | Negative-control external trait partitioned LD score regression: C0, C1 and C2 cluster enrichments across two well-powered non-psychiatric, non-brain GIANT 2018 sumstats.**

All regressions condition on the 97-annotation baseline-LD v2.2 model²⁴ over the EUR LDSC reference. The C0/C1/C2 cluster annotations are identical to those used for the PGC3 EUR SCZ discovery analysis (Methods, manuscript main text, Table 2). Both Yengo *et al.* 2018 GIANT sumstats (Wood + UKBiobank for height; Locke + UKBiobank for BMI) were munged to LDSC format using a GIANT-aware adapter (Methods §Negative-control external traits). The negative-control test asks whether the SCZ-derived C0 cluster annotation captures generic ancient-allele or non-brain polygenic architecture (in which case it should enrich for height and BMI in proportion to their h²) or psychiatric-architecture-specific variation (in which case it should show null enrichment for non-psychiatric anthropometric traits).

| **Trait (sumstats)** | **Cluster** | **Prop. SNPs** | **Prop. h²** | **Prop. h² s.e.** | **Enrichment** | **s.e.** | ***P*** | **Coef. *Z*** | **Trait h²** | **LDSC intercept** | **Mean χ²** |
| --- | --- | --- | --- | --- | --- | --- | --- | --- | --- | --- | --- |
| **Yengo 2018 GIANT height⁸⁹** (mean *N* = 696,975) | C0 | 0.000288 | 0.00217 | 0.00212 | 7.55× | 7.36 | 0.374 | +0.62 | 0.527 | 1.560† | 8.44 |
|  | C1 | 0.000261 | −0.000252 | 0.000265 | −0.97× | 1.02 | 0.057 | −1.21 | — | — | — |
|  | C2 | 0.000357 | 0.000124 | 0.00153 | 0.35× | 4.29 | 0.879 | −0.67 | — | — | — |
| **Yengo 2018 GIANT body-mass index⁸⁹** (mean *N* = 686,182) | C0 | 0.000288 | −0.00107 | 0.00174 | −3.73× | 6.04 | 0.435 | −0.99 | 0.217 | 1.059† | 3.94 |
|  | C1 | 0.000261 | 0.000586 | 0.000737 | 2.25× | 2.83 | 0.658 | +0.71 | — | — | — |
|  | C2 | 0.000357 | 0.000957 | 0.00235 | 2.68× | 6.57 | 0.797 | +0.03 | — | — | — |

† Yengo 2018 height has elevated LDSC intercept (1.56) and high Lambda GC (3.63) reflecting both the very large effective sample size and residual stratification not fully absorbed by the LDSC intercept; the cluster-level conclusion (null C0 enrichment) is unaffected because the residual structure is absorbed by the intercept rather than spuriously attributed to the cluster annotation. The BMI intercept (1.06) is essentially clean, providing the cleaner negative-control comparison.

*The C0 (Young) cluster shows null enrichment for both height (7.55×;* P *= 0.37; conditional* Z *= +0.62) and BMI (−3.73×;* P *= 0.44;* Z *= −0.99), despite both traits having substantially larger effective sample size and polygenic signal-to-noise (mean χ² 8.44 and 3.94 respectively) than the PGC3 EUR schizophrenia analogue (mean χ² 2.02). The C0 cluster annotation therefore does not capture generic ancient-allele or non-brain polygenic architecture, indicating that the within-psychiatric-rg-shared signal in cross-disorder analyses (Supplementary Table 11) is specific to psychiatric (and brain-relevant) genetic architecture rather than reflecting broader genome-wide ancient-allele or anthropometric architecture. The C1 height nominal* P *= 0.057 (Z = −1.21) does not survive the focal-annotation significance threshold (Z ≈ 2.81) and is in the opposite direction (negative enrichment); we interpret this as noise.*

**Supplementary Table 14 | Two-dimensional GMM sensitivity (allele age × |iHS| only) — joint-feature requirement check.**

To test whether the principal Young-cluster heritability concentration is robust to dropping the brain–blood regulatory specificity dimension from the cluster definition, we re-fit the Gaussian mixture model on only two evolutionary features — log₁₀ allele age and Voight-correct |iHS| — and ran identical partitioned LD score regression on PGC3 EUR schizophrenia. The two-dimensional GMM achieves substantially higher coverage of the credible-set variant set (15,739 of 18,895 non-MAPT variants; 83.3%) versus the three-dimensional primary (4,918 variants; 26.0%) by removing the requirement for both GTEx brain and blood eQTLs.

| **Trait** | **Cluster** | **Prop. SNPs** | **Prop. h²** | **Prop. h² s.e.** | **Enrichment** | **s.e.** | ***P*** | **Coef. *Z*** |
| --- | --- | --- | --- | --- | --- | --- | --- | --- |
| **PGC3 EUR SCZ** (3D primary, k=3, 4,918 variants) | C0 (Young) | 0.000288 | 0.0136 | 0.00427 | **47.35×** | 14.86 | **2.0 × 10⁻³** | **+3.05** |
|  | C1 (Mid) | 0.000261 | 0.0034 | 0.00101 | **13.04×** | 3.89 | **2.1 × 10⁻³** | **+3.32** |
|  | C2 (Old) | 0.000357 | 0.00558 | 0.0046 | 15.61× | 12.89 | 0.259 | +1.04 |
| **PGC3 EUR SCZ** (2D sensitivity, k=3, 15,739 variants) | C0 (Young) | 0.000976 | 0.0295 | 0.0172 | 30.26× | 17.58 | 0.101 | +1.61 |
|  | C1 (Mid) | 0.000363 | 0.00896 | 0.00791 | 24.67× | 21.77 | 0.275 | +1.10 |
|  | C2 (Old) | 0.0013 | 0.0627 | 0.0147 | **48.22×** | 11.27 | **5.6 × 10⁻⁵** | **+4.10** |

**Interpretation.** The 2D Young cluster (n=5,820 in EUR LD reference, 3.3× larger than 3D Young's 1,744 by inclusion of an additional ~4,200 credible-set variants previously uncoverable in 3D for lack of GTEx brain∩blood eQTL data) shows attenuated, non-significant heritability concentration (30.3×; *P* = 0.101). Conversely, the 2D Old cluster (n=7,753) is the dominant heritability-concentrating cluster (48.2×; *Z* = +4.10). The 3D Young cluster overlaps the 2D Young cluster at 92.5% recall (Jaccard 0.27 with the broader 2D set), confirming that 3D Young is a regulatory-rich subset of 2D Young. The differential heritability concentration across these two cluster definitions establishes that the 3D primary signal cannot be reduced to age + selection alone: the Young cluster's heritability concentration in the 3D primary requires the joint requirement of young allele age, low–moderate |iHS| and brain-restricted *cis*-regulatory specificity. Adding the ~4,200 variants without brain∩blood eQTL data (presumably tag variants without functional regulatory consequence in expression data) dilutes the per-variant heritability and shifts the heritability mass to the older, regulatorily uncharacterised credible-set sub-population. We therefore interpret the 2D sensitivity result as evidence that the brain–blood regulatory specificity dimension is functionally informative — not a redundant feature loaded into the cluster definition.

*Cluster-level partitioned heritability (stratified LD-score regression conditional on the baseline-LD v2.2 model) for PGC3 European schizophrenia under the three-dimensional primary (n = 4,918 variants) versus two-dimensional sensitivity (n = 15,739) Gaussian mixture cluster definitions. Enrichment, per-SNP heritability fold-enrichment; the s.e. column is the enrichment standard error; P, LDSC block-jackknife enrichment-test P; Coef. Z, conditional coefficient Z-score (focal-annotation significance threshold 2.81). GMM, Gaussian mixture model. Two-dimensional sensitivity check of the cluster definition in Fig. 1 and Supplementary Table 5.*

**Supplementary Table 15 | Block-jackknife n_blocks sensitivity (Tashman 2021 small-annotation robustness).**

To address the Tashman *et al.* 2021 caution that S-LDSC block-jackknife significance testing may fail to control type-1 error for annotations smaller than 0.5% of regression-weight SNPs, we re-ran partitioned LD score regression for PGC3 EUR schizophrenia under increased block count (`--n-blocks 1000`, five-fold the LDSC v1.0.1 default of 200). The C0 (Young) cluster comprises 0.029% of HapMap3 SNPs, well below Tashman's 0.5% threshold.

| **Cluster** | **n_blocks** | **Enrichment** | **s.e.** | ***P*** | **Coef. *Z*** |
| --- | --- | --- | --- | --- | --- |
| **C0 (Young)** | 200 (default) | **47.35×** | 14.86 | **2.0 × 10⁻³** | **+3.05** |
| **C0 (Young)** | 1,000 (Tashman) | **47.35×** | 14.05 | **1.0 × 10⁻³** | **+3.20** |
| **C1 (Mid)** | 200 (default) | **13.04×** | 3.89 | **2.1 × 10⁻³** | **+3.32** |
| **C1 (Mid)** | 1,000 (Tashman) | **13.04×** | 3.85 | **1.7 × 10⁻³** | **+3.34** |
| C2 (Old) | 200 (default) | 15.61× | 12.89 | 0.259 | +1.04 |
| C2 (Old) | 1,000 (Tashman) | 15.61× | 12.73 | 0.250 | +1.06 |

**Interpretation.** Point estimates are identical between n_blocks=200 and n_blocks=1000; only the standard errors differ marginally. With 1,000 blocks, both significant clusters (C0 and C1) yield slightly stronger evidence (lower *P*, higher conditional *Z*) than with the default 200 blocks. The Tashman-recommended block count therefore confirms rather than weakens the principal Young-cluster heritability concentration finding, indicating that the v10 default-block-count results are not driven by under-resolved jackknife variance estimation.

*Partitioned heritability for PGC3 European schizophrenia (n = 4,918 variants; baseline-LD v2.2) at the default 200-block versus 1,000-block jackknife resolution; point estimates are invariant to block count and only the standard errors differ. Enrichment, s.e., P and Coef. Z are defined as in Supplementary Table 14.*

**Supplementary Table 16 | GMM component-number (*k*) sensitivity for the principal Young-cluster heritability concentration.**

The primary cluster definition uses *k* = 3 by parsimony, despite BIC monotonically favouring larger *k* over the candidate range *k* = 1–6 (Supplementary Table 5). To test whether the principal Young-cluster heritability concentration is robust to this choice, we re-fit the GMM on the same 3D feature space (log_age × brain_spec × |iHS|, no MAPT) at *k* = 2 and *k* = 4, identified the Young cluster at each *k* as the component with the lowest mean unscaled log₁₀ allele age, built binary annotations from the resulting Young-cluster assignments, and ran identical partitioned LD score regression on PGC3 EUR schizophrenia.

**Section A. Young-cluster size and centroid across *k***

| **GMM *k*** | **Young cluster *n* (GMM-assigned, no MAPT)** | **Young cluster *n* (EUR LD reference)** | **Mean log₁₀ allele age (Young centroid)** | **Mean brain–blood specificity** | **Mean \** | **iHS\** |
| --- | --- | --- | --- | --- | --- | --- |
| 2 | 1,785 | 1,784 | 5.061 | 0.505 | 0.730 |  |
| **3 (primary)** | **1,745** | **1,744** | **5.054** | **0.506** | **0.701** |  |
| 4 | 1,625 | 1,624 | 5.058 | 0.504 | 0.605 |  |

The Young-cluster size (~1,700 GMM-assigned variants) and centroid (mean log₁₀ allele age ≈ 5.06 [log₁₀ years; generations × 28.1], mean brain–blood specificity ≈ 0.50, mean |iHS| ≈ 0.6–0.7) are stable across *k*, indicating that the Young sub-population is a robust feature of the joint feature space rather than an artefact of *k* = 3 selection.

**Section B. Single-annotation partitioned LD score regression of Young cluster (PGC3 EUR SCZ)**

| **GMM *k*** | **Young Prop. SNPs** | **Prop. h²** | **Prop. h² s.e.** | **Enrichment** | **s.e.** | ***P*** | **Coef. *Z*** |
| --- | --- | --- | --- | --- | --- | --- | --- |
| **2** | 0.000299 | 0.02897 | 0.00541 | **96.81×** | 18.09 | **2.4 × 10⁻⁷** | **+5.28** |
| **3 (primary)** | 0.000288 | 0.01362 | 0.00427 | **47.35×** | 14.86 | **2.0 × 10⁻³** | **+3.05** |
| **4** | 0.000272 | 0.0287 | 0.0049 | **105.34×** | 18.00 | **2.0 × 10⁻⁸** | **+5.77** |

**Interpretation.** The Young-cluster heritability concentration is significantly elevated at all three values of *k* tested. The *k* = 3 primary uses the full three-cluster (C0/C1/C2) model in which the Young-cluster enrichment is conditional on the Mid and Old clusters being separately partitioned; the *k* = 2 and *k* = 4 sensitivity analyses use single-annotation models in which the Young-cluster captures the total Young-specific-or-correlated heritability that would otherwise be split across multiple cluster annotations. The single-annotation *k* = 2 and *k* = 4 conditional *Z*-scores (+5.28 and +5.77) accordingly exceed the multi-annotation *k* = 3 *Z* (+3.05) by absorbing additional Young-correlated heritability mass. The qualitative finding — significant heritability concentration in the Young cluster — is robust to GMM *k* selection across *k* ∈ {2, 3, 4}.

*Single-annotation partitioned heritability of the Young cluster for PGC3 European schizophrenia (n = 4,918 variants; baseline-LD v2.2) at Gaussian mixture k = 2, k = 3 (primary, multi-annotation) and k = 4. Columns are defined as in Supplementary Table 14.*

**Supplementary Table 17 | Coverage selection bias quantification: GMM-included vs GMM-excluded credible-set variants.**

To address concern about discovery-substrate selection bias, we quantify the GTEx and single-cell brain-regulatory profile of the 13,977 credible-set variants excluded from the 3D GMM cluster input (those lacking at least one of: GTEx brain *cis*-eQTL nominal *P*, GTEx blood *cis*-eQTL nominal *P*, Voight |iHS|, or Atlas of Variant Age estimate). Allele ages are reported in years, obtained from the Atlas of Variant Age `age_median` (estimated in generations) converted at 28.1 yr generation⁻¹ (Wohns et al. 2022), consistent with the main text.

**Section A. GMM-included vs excluded counts (non-MAPT)**

| **Subset** | **n variants** | **% of non-MAPT credible-set** | **Median allele age (yr)** |
| --- | --- | --- | --- |
| 3D GMM input (all 4 features) | 4,918 | 26.0% | ~384,000 |
| GMM-excluded | 13,977 | 74.0% | ~316,000 |

**Section B. GTEx *cis*-eQTL coverage among GMM-excluded variants**

| **GTEx coverage profile** | **n variants** | **% of GMM-excluded** |  |  |
| --- | --- | --- | --- | --- |
| Brain eQTL only (no blood eQTL) | 3,257 | 23.3% |  |  |
| Blood eQTL only (no brain eQTL) | 1,057 | 7.6% |  |  |
| Both brain and blood eQTL (but no | iHS | or age) | 872 | 6.2% |
| Neither brain nor blood eQTL (QTL-silent) | 8,791 | 62.9% |  |  |

**Important interpretive points**:

- 23.3% of GMM-excluded variants have detectable GTEx brain *cis*-eQTL but no detectable GTEx blood *cis*-eQTL — i.e., they are *more* brain-specific than the GMM-included subset (which by definition has signal in both compartments). These variants cannot be placed on our brain–blood specificity axis simply because the denominator (blood signal) is unmeasured, not because they lack brain-regulatory function.

- The 62.9% with detectable signal in neither compartment likely represent (i) tag variants in long-LD blocks where the causal regulatory variant is elsewhere, (ii) tissue-context-specific eQTLs not assayed in adult bulk GTEx, or (iii) intergenic credible-set variants with no detected *cis*-regulatory function in current expression atlases.

- The median allele age difference (in-GMM ~384,000 yr vs out-of-GMM ~316,000 yr; medians of 13,681 vs 11,235 Atlas-of-Variant-Age generations × 28.1 yr generation⁻¹) is modest: the GMM-included subset is on average ~22% older (median ratio 13,681 / 11,235 = 1.22; this relative difference is unit-invariant) — consistent with the requirement for detectable eQTL signal favouring established, regulatory-active common variants. Both subsets are deeply ancient and predominantly pre-out-of-Africa (≫65 kyr), so this selection effect shifts allele age only marginally within an already ancestral distribution.

**Section C. Implication for the principal finding**

The 3D GMM cluster annotation is a property of expression-annotated credible-set variants (26% of the non-MAPT credible-set substrate). The principal Young-cluster heritability concentration (47.4×; *P* = 2.0 × 10⁻³) should therefore be interpreted as: "among PGC3 credible-set variants with both brain and blood GTEx *cis*-eQTL annotation, the joint-feature-defined Young sub-population concentrates SCZ heritability". The 2D feature-space sensitivity (dropping the brain-blood requirement and recovering 83% coverage) recovers a non-significant Young cluster (30.3×, *P* = 0.101; Supplementary Table 14), confirming that the partitioned-heritability signal is a property of the joint feature combination — not of any single component including allele age alone — and not a generic feature of all credible-set variants regardless of regulatory annotation.

*Discovery-substrate coverage comparison of GMM-included (n = 4,918) versus GMM-excluded (n = 13,977) non-MAPT fine-mapped credible-set variants, by GTEx and single-cell cis-eQTL coverage and allele age (Atlas of Variant Age, converted generations→years at 28.1 yr generation⁻¹); central values are medians and the ~22% median age ratio is unit-invariant. eQTL, expression quantitative trait locus.*

**Supplementary Table 18 | MAPT-included partitioned LDSC sensitivity.**

To test whether the principal Young-cluster heritability concentration is driven by exclusion of the 17q21.31 MAPT inversion, we re-fit the primary 3D GMM (k = 3, random_state = 42, n_init = 10) on the non-MAPT training set (n = 4,918; identical to the P14b primary; 100% reproducibility of cluster assignments verified at the variant level) and then predicted cluster labels for the 1,568 MAPT credible-set variants with all three features using the same trained model and StandardScaler. MAPT variants were thus held-out predictions, not training inputs — the primary cluster definition is preserved exactly. Per-chrom cluster annotation files were rebuilt; only chr17 differs from the MAPT-excluded primary annotation (other 21 chromosomes byte-identical). chr17 LD scores were recomputed with HM3 SNP restriction. Partitioned S-LDSC was run on PGC3 EUR schizophrenia.

**Section A. MAPT cluster-prediction distribution**

| **Cluster** | **Non-MAPT n** | **MAPT added n** | **Combined n** |
| --- | --- | --- | --- |
| C0 (Young) | 1,745 (4,918 GMM-input cohort) | 117 | 1,862 |
| C1 (Mid) | 317 | 103 | 420 |
| C2 (Old) | 2,856 | 1,348 | 4,204 |

The C2-dominant MAPT distribution (1,348 / 1,568 = 86%) is expected: the 17q21.31 inversion is an ancient structural variant with extended long-range LD across ~3 Mb, and the typical MAPT-locus credible-set variant has high allele age (consistent with C2 centroid ≈ 511 kyr; 18,188 generations × 28.1 yr generation⁻¹).

**Section B. S-LDSC enrichment for PGC3 EUR SCZ — MAPT-included vs MAPT-excluded primary**

| **Cluster** | **Cluster annotation** | **Enrichment** | **s.e.** | ***P*** | **Coef. *Z*** |
| --- | --- | --- | --- | --- | --- |
| C0 (Young) | MAPT-excluded (primary) | 47.4× | 14.86 | 2.0 × 10⁻³ | +3.05 |
|  | MAPT-included | 47.8× | 14.69 | 1.6 × 10⁻³ | +3.12 |
| C1 (Mid) | MAPT-excluded (primary) | 13.0× | 3.89 | 2.1 × 10⁻³ | +3.32 |
|  | MAPT-included | 17.3× | 5.17 | 1.7 × 10⁻³ | +3.34 |
| C2 (Old) | MAPT-excluded (primary) | 15.6× | 12.89 | 0.26 | +1.04 |
|  | MAPT-included | 8.5× | 10.36 | 0.47 | +0.69 |

Total observed-scale h² = 0.370 ± 0.013 (mean χ² = 2.02; intercept = 1.093 ± 0.014) — essentially identical to primary.

**Interpretation.** The C0 (Young) heritability concentration is robust to MAPT inclusion (47.4× → 47.8×, conditional Z = +3.05 → +3.12). C1 (Mid) increases (13.0 → 17.3×) because the 103 high-|iHS| MAPT variants assigned to C1 carry additional heritability mass. C2 (Old) decreases (15.6 → 8.5×) because the 1,348 MAPT variants assigned to C2 are mostly low-effect-size due to extended LD across the inversion, diluting C2's per-SNP heritability density. The Bonferroni-significant C0 and C1 enrichments survive both with and without MAPT.

*Cluster-level partitioned heritability for PGC3 European schizophrenia with the 17q21.31 MAPT inversion excluded from (primary; n = 4,918 variants) versus included in the cluster annotation (baseline-LD v2.2). Δ enrichment (included − excluded): C0 +0.4×, C1 +4.3×, C2 −7.1×. Columns are defined as in Supplementary Table 14.*

**Supplementary Table 19 | Tissue-power-matched brain_spec sensitivity.**

GTEx v10 tissue donor N differs substantially: Whole_Blood N ≈ 838, individual brain tissues N ≈ 140–280. For an identical effect size β, the P-value scales as Z = β·sqrt(N) → smaller-P in higher-N tissues. The original brain_spec = -log10(brain_minp) / [-log10(brain_minp) + -log10(blood_minp)] is therefore systematically biased downward (Whole_Blood signal artificially inflated relative to brain).

**Section A. Z-statistic reconstruction and rescaling**

For each variant with both brain and blood eQTL minimum-P-value annotations:

1. |Z|_observed = -Φ⁻¹(P/2) (two-sided inverse normal)

2. Tissue-specific donor N (GTEx v10 release notes)

3. Equal-N reference: N_ref = 220 (median brain tissue donor N)

4. |Z|_eq = |Z|_observed × sqrt(N_ref / N_tissue)

5. P_eq = 2·Φ(-|Z|_eq)

6. brain_spec_eq = -log10(P_eq_brain) / [-log10(P_eq_brain) + -log10(P_eq_blood)]

**Section B. brain_spec distribution before vs after power-matching**

| **Statistic** | **brain_spec (original)** | **brain_spec_eq (power-matched)** |
| --- | --- | --- |
| n (both eQTLs available) | 7,470 | 7,470 |
| Mean | 0.433 | 0.626 |
| Median | 0.437 | 0.586 |
| Std | 0.161 | 0.145 |
| Correlation (orig vs eq) | r = 0.696 | — |
| Median shift (orig − eq) | −0.255 | — |
| Variants flipping orig ≤0.5 → eq >0.5 (more brain-biased after correction) | 3,687 | — |
| Variants flipping orig >0.5 → eq ≤0.5 | 19 | — |

**Direction**: 3,687 variants (49.4% of overlapping set) were classified as "blood-biased" (≤ 0.5) under the original brain_spec but become "brain-biased" (> 0.5) after power-matching. The original brain_spec systematically under-estimates brain bias by ~25%.

**Section C. Re-fitted 3D GMM (log_age × brain_spec_eq × |iHS|)**

| **Cluster** | **n GMM** | **log_age centroid** | **brain_spec_eq centroid** |  | **iHS** | **centroid** | **Mean age (yr)** |
| --- | --- | --- | --- | --- | --- | --- | --- |
| C0 (Young) | 1,760 | 5.056 | 0.681 | 0.732 | 113,608 |  |  |
| C1 (Mid) | 1,052 | 5.639 | 0.474 | 1.217 | 435,297 |  |  |
| C2 (Old) | 2,106 | 5.725 | 0.727 | 0.718 | 530,050 |  |  |

Versus primary (original brain_spec):

| **Cluster** | **Primary n** | **Primary age (yr)** | **Primary brain_spec** | **Primary** | **iHS** |
| --- | --- | --- | --- | --- | --- |
| C0 (Young) | 1,745 | 113,215 | 0.506 | 0.701 |  |
| C1 (Mid) | 317 | 360,102 | 0.404 | 1.892 |  |
| C2 (Old) | 2,856 | 511,083 | 0.491 | 0.740 |  |

**Notable shifts**: Under power-matching, C0 (Young) is more strongly brain-biased (centroid 0.51 → 0.68); C1 (Mid) is more populous (n 317 → 1,052) but with lower mean |iHS| (1.89 → 1.22) — power-matching reveals that more variants exhibit moderate |iHS| in the Mid range than is apparent under the original brain_spec; C2 (Old) shifts to brain-biased (0.49 → 0.73).

**Section D. Partitioned S-LDSC for PGC3 EUR SCZ with power-matched cluster annotation**

| **Cluster** | **Cluster annotation** | **Enrichment** | **s.e.** | ***P*** | **Coef. *Z*** |
| --- | --- | --- | --- | --- | --- |
| C0 (Young) | Original brain–blood specificity (primary) | 47.4× | 14.86 | 2.0 × 10⁻³ | +3.05 |
|  | Power-matched (brain_spec_eq) | 71.9× | 19.91 | 4.1 × 10⁻⁴ | +3.53 |
| C1 (Mid) | Original brain–blood specificity (primary) | 13.0× | 3.89 | 2.1 × 10⁻³ | +3.32 |
|  | Power-matched (brain_spec_eq) | 22.7× | 10.96 | 0.049 | +1.92 |
| C2 (Old) | Original brain–blood specificity (primary) | 15.6× | 12.89 | 0.26 | +1.04 |
|  | Power-matched (brain_spec_eq) | 36.6× | 11.57 | 2.6 × 10⁻³ | +3.00 |

Total observed-scale h² = 0.368 ± 0.013 (mean χ² = 2.02; intercept = 1.094 ± 0.014) — essentially identical to primary.

**Interpretation.** Power-matching GTEx brain–blood specificity for tissue sample-size asymmetry (N_blood ≈ 838 vs N_brain ≈ 144–276) does not weaken the principal Young-cluster heritability concentration; on the contrary, the C0 enrichment strengthens from 47.4× to 71.9× and the conditional *Z*-score increases from +3.05 to +3.53. The C2 (Old) cluster transitions from non-significant (*P* = 0.26 in primary) to Bonferroni-significant (*P* = 2.6 × 10⁻³ in power-matched), suggesting that the original brain_spec formulation systematically under-estimated heritability concentration in both the Young and Old clusters by absorbing brain-vs-blood power asymmetry into the brain_spec axis. The power-matched cluster definition, despite materially changing the SNP partition (different cluster sizes; different brain_spec_eq centroids), recovers the same qualitative architecture (Young carries strongest enrichment) with stronger statistical support. Tissue sample-size asymmetry therefore does not spuriously inflate the C0 enrichment through the brain–blood specificity axis; correcting the asymmetry strengthens the primary finding.

*Cluster-level partitioned heritability for PGC3 European schizophrenia under the original versus tissue-power-matched brain–blood specificity definition (baseline-LD v2.2). Δ enrichment (power-matched − original): C0 +24.5×, C1 +9.6×, C2 +21.0×; Δ Coef. Z: +0.49, −1.40, +1.96. Columns are defined as in Supplementary Table 14.*

**Supplementary Table 19A | Brain–blood specificity-only single-annotation S-LDSC and joint conditional model.**

To test whether brain–blood specificity alone — without allele age or |iHS| — yields the same heritability concentration as the three-dimensional Young cluster, we built four S-LDSC tests on PGC3 EUR schizophrenia: (A) top-1,742 brain_spec binary, (B) continuous brain_spec, (C) C0-only single-annotation (apples-to-apples comparison with brain_spec — same baseline-only conditioning), and (D) joint conditional model with both brain_spec AND C0 in the same regression.

**Section A — Single-annotation enrichments (each annotation tested alone against baseline-LD v2.2)**

| **Annotation** | **n** | **Prop. SNPs** | **Prop. h²** | **Enrichment** | **s.e.** | ***P*** | **Coef. *Z*** |
| --- | --- | --- | --- | --- | --- | --- | --- |
| Primary 3D Young C0 (multi-annot: C0+C1+C2) | 1,744 | 0.000288 | 0.01362 | 47.4× | 14.86 | 2.0 × 10⁻³ | +3.05 |
| Top-1,742 brain_spec only (single) | 1,723 | 0.00028 | 0.01528 | 54.5× | 9.15 | 3.2 × 10⁻⁸ | +5.67 |
| Continuous brain_spec (single) | 5,790 | 0.000459 | 0.01808 | 39.4× | 4.77 | 1.1 × 10⁻¹³ | +7.82 |
| **C0-only (single annotation)** | 1,744 | 0.000293 | 0.02930 | **100.2×** | 17.63 | **4.6 × 10⁻⁸** | **+5.61** |

**Section B — Joint conditional model (brain_spec + C0 BOTH in same model)**

| **Annotation in joint model** | **Prop. SNPs** | **Prop. h²** | **Enrichment** | **s.e.** | ***P*** | **Conditional *Z** |
| --- | --- | --- | --- | --- | --- | --- |
| brain_spec_high (top-1,742) | 0.00028 | 0.01391 | 49.6× | 9.79 | 2.5 × 10⁻⁶ | +1.26 |
| **C0_only** | 0.000293 | 0.0282 | **96.4×** | 17.92 | 1.8 × 10⁻⁷ | **+4.80** |

**Section C — Overlap structure between C0 and top-1,742 brain_spec**

| **Subset** | **n SNPs** | **brain_spec median** | **brain_spec range** |
| --- | --- | --- | --- |
| Intersection (in both C0 AND Top-brain_spec) | 663 | 0.632 | 0.553–0.873 |
| Top-only (in Top, NOT in C0) | 1,060 | 0.627 | 0.553–0.880 |
| **C0-only (in C0, NOT in Top)** | 1,081 | **0.412** | 0.106–0.553 |
| Jaccard index (overlap / union) | 0.236 | — | — |

**Interpretation**

The single-annotation comparison (Section A) shows C0-only S-LDSC yields ~2× higher enrichment than top-1,742 brain_spec alone (100.2× vs 54.5×; *Z* = +5.61 vs +5.67; both highly significant). The joint cluster definition concentrates substantially more heritability per SNP than brain_spec ranking — refuting the apparent finding from the multi-annotation model (47.4× with *Z* = +3.05) where C1 and C2 partition heritability away from C0.

The joint conditional model (Section B) is the definitive test: **C0 retains *Z* = +4.80 conditional on brain_spec (Bonferroni-significant; *P* = 1.8 × 10⁻⁷), whereas brain_spec collapses to *Z* = +1.26 conditional on C0 (not significant)**. The joint 3D feature definition therefore carries primary independent signal; brain_spec ranking is largely collinear with — and absorbed by — the C0 cluster. The cluster annotation is NOT redundant with brain-regulatory specificity ranking.

The overlap structure (Section C) confirms this. C0-only variants (62% of C0, n=1,081) have brain_spec median 0.41 — below the genome-wide ascertained median (0.44) and well below the top-1,742 threshold (0.55). These variants enter C0 because of their young allele age + low–moderate |iHS| combination, not because of high brain_spec. Yet C0 in aggregate (including the moderate-brain_spec C0-only majority) yields 100.2× single-annotation heritability concentration. The joint feature combination — young age + brain-blood specificity (even moderate) + low–moderate |iHS| — captures a heritability-concentrating sub-population that brain_spec ranking alone misses.

*Single-annotation (Section A) and joint conditional (Section B) partitioned heritability for PGC3 European schizophrenia (baseline-LD v2.2); n is the number of SNPs in each annotation. Section A tests each annotation alone against the baseline-LD model; Section B includes brain–blood specificity and the C0 cluster jointly. Columns are defined as in Supplementary Table 14.*

**Supplementary Table 20 | Empirical matched-LD-MAF permutation null.**

Tashman 2021 recommends an empirical permutation null for partitioned LD score regression of small annotations (< 0.5% of HapMap3 SNPs). The C0 (Young) cluster comprises 0.029% of HapMap3 SNPs, well below this threshold. We construct a null distribution by sampling random C0-sized SNP sets matched to the primary C0 cluster in their joint distribution over baseline-L2 decile × EUR-MAF decile bins (10 × 10 = 100 bins), then running identical partitioned S-LDSC on each random set.

**Section A. Permutation methodology**

- 100 LD-bin × MAF-bin combinations (decile × decile)

- For each permutation, per bin, draw n_C0_bin SNPs without replacement from the HM3 master pool matched on bin_id; combine across bins → 1,744-SNP random C0_perm; build per-chr annotation files; compute LD scores; run S-LDSC.

**Section B. Empirical null distribution (N=15 matched-LD-MAF permutation draws)**

| **Statistic** | **Observed (primary C0 multi-annot)** | **Null distribution (N=15)** |
| --- | --- | --- |
| Enrichment | 47.35× | mean +4.76× (s.d. 17.43); range −17.56× to +31.87× |
| Conditional *Z* | **+3.05** | mean **+0.133** (s.d. 0.920); range −1.19 to **+1.61** |
| Prop. h² | 0.01362 | mean 0.00139 (s.d. 0.00509) |
| **Test** | **Value** | **Interpretation** |
| Empirical P (# null Z ≥ +3.05) / N | **0/15 = < 0.062** (1/16 with sample-size correction) | Observed Z outside the null distribution |
| Empirical Z (parametric: (obs − null mean) / null s.d.) | **+3.17** | *P* ≈ 7.6 × 10⁻⁴ (one-sided normal approximation) |
| Null max Z | **+1.61** | Below observed Z = +3.05 |
| Null Z ≥ +3.0 | 0/15 | None of 15 random draws reach observed Z magnitude |

**Section C. Comparison with the single-annotation result**

The single-annotation C0 test is even further outside the null distribution: 0/15 null draws reach Z ≥ +5.0; the empirical Z (parametric) for the single-annotation observation is (5.61 − 0.133) / 0.920 = +5.96 (empirical *P* < 0.0001 by normal approximation).

**Section D. Limitations**

We acknowledge that N = 15 permutation draws is smaller than the Tashman 2021 recommendation of ≥ 100 perms; this was a pragmatic compromise due to per-permutation S-LDSC computation time (~25 minutes per permutation on the available hardware, requiring ~6 hours total for 15 perms). The N = 15 sample is sufficient to establish that the observed Z lies clearly outside the null range (0/15 draws ≥ observed) but cannot resolve empirical P-values below approximately 1/16 ≈ 0.063 directly. The parametric empirical Z (under normal approximation of the null Z distribution) extends the resolution to P ≈ 0.026, but this normal assumption may not hold at the tail. A more conservative interpretation is therefore: the observed C0 multi-annotation Z = +3.05 is directionally and rank-order significant at the 0/15 = empirical-P < 1/16 level, with the asymptotic LDSC jackknife enrichment P-value (2.0 × 10⁻³) and the parametric empirical Z (+3.17) converging on the same qualitative conclusion that the observed enrichment is unlikely to arise under the matched-LD-MAF null. Future analyses with larger permutation sample sizes (N ≥ 100) and additional null specifications (e.g., chromosome-block-shuffle null, gene-set-matched null) would tighten the empirical resolution.

**Interpretation**

The observed primary C0 (Young) cluster heritability concentration (multi-annot Z = +3.05, single-annot Z = +5.61) lies outside the null distribution of matched-LD-MAF SNP draws. 0/15 permutations achieve Z ≥ +3.05, and 0/15 achieve Z ≥ +3.0. The N = 15 empirical-P upper bound of 1/16 (≈ 0.063) is sample-size-limited; an extended N = 1,000 matched-LD-MAF null (Methods) places the observed Z at the 100th percentile (0/1,000 draws ≥ observed; empirical P = 1.0 × 10⁻³, parametric P = 7.4 × 10⁻⁴), and the parametric empirical Z (+3.17 for multi-annot; +5.96 for single-annot) is consistent with the asymptotic LDSC jackknife enrichment P-value of 2.0 × 10⁻³ in qualitative significance. The Tashman 2021 small-annotation type-1-error concern is therefore addressed through three independent robustness anchors: (i) `--n-blocks 1000` fine-resolution jackknife (Supplementary Table 15), (ii) the matched-LD-MAF empirical null (Supplementary Table 20), and (iii) the joint conditional analysis with brain_spec annotation (Supplementary Table 19A).

*Empirical matched-LD-MAF permutation null for the C0 (Young) cluster enrichment in PGC3 European schizophrenia (baseline-LD v2.2). Random SNP sets matched to C0 on baseline-LD-score decile × EUR-MAF decile bins (N = 15, extended to N = 1,000); the null distribution is summarised as mean ± s.d. Empirical P = (number of null draws with Z ≥ observed + 1)/(N + 1). Observed values are the primary multi-annotation C0 estimates.*

**Supplementary Table 21 | Allele age distribution of PGC3 SCZ credible-set variants: a deep, pre-Out-of-Africa age structure.**

The Atlas of Variant Age² reports coalescent-based ages in generations; we convert to years at 28.1 yr generation⁻¹ (Wohns et al. 2022; main-text Methods §Allele age and Limitations). After conversion, the 20,565 PGC3 credible-set variants with a measurable estimate span deep evolutionary time: the median age is ~382 kyr and ~95% of variants predate the Out-of-Africa expansion (≥65 kyr). This table quantifies the distribution to support the main-text framing that the credible set is ancient and ancestrally shared (pre-OOA), with persistence at common frequency explained by purifying selection and mutation–selection balance rather than recent (post-OOA) origin. The k=3 cluster definition partitions the youngest, intermediate, and oldest strata within this single deeply ancestral set, not a narrow post-OOA window.

**Section A. Quantile distribution (n=20,565)**

| **Quantile** | **GEVA (generations)** | **Age (yr)** | **Age (kyr)** |
| --- | --- | --- | --- |
| Q0.1 | 711 | 19,990 | 20.0 |
| Q1.0 | 1,382 | 38,824 | 38.8 |
| Q5.0 | 2,378 | 66,824 | 66.8 |
| Q25.0 | 4,461 | 125,363 | 125.4 |
| **Q50 (median)** | **13,601** | **382,182** | **382.2** |
| Q75.0 | 18,866 | 530,143 | 530.1 |
| Q95.0 | 27,000 | 758,708 | 758.7 |
| Q99.0 | 33,937 | 953,617 | 953.6 |
| Q99.9 | 49,259 | 1,384,173 | 1,384.2 |
| **Max** | **115,545** | **3,246,815** | **3,246.8 (≈3.25 Myr)** |

**Section B. Variant counts by age threshold**

| **Threshold (Age in years)** | **n variants ≥ threshold** | **%** |
| --- | --- | --- |
| **≥ 65,000 yr (pre-Out-of-Africa)** | **19,634** | **95.47%** |
| ≥ 100,000 yr | 17,207 | 83.67% |
| **≥ 200,000 yr** | **13,014** | **63.28%** |
| ≥ 500,000 yr | 6,290 | 30.59% |
| ≥ 1,000,000 yr (1 Myr) | 152 | 0.74% |
| ≥ 2,000,000 yr (2 Myr) | 1 | 0.005% |

**Section C. Histogram bins (Age in kyr)**

| **Age bin** | **n variants** | **% of total** |
| --- | --- | --- |
| 0–65 kyr (post-OOA, youngest stratum) | 931 | 4.53% |
| 65–100 kyr | 2,427 | 11.80% |
| 100–200 kyr | 4,193 | 20.39% |
| 200–500 kyr | 6,724 | 32.70% |
| 500–1,000 kyr (0.5–1 Myr) | 6,138 | 29.85% |
| 1,000–2,000 kyr (1–2 Myr) | 151 | 0.73% |
| ≥ 2,000 kyr (>2 Myr) | 1 | 0.005% |

**Section D. Interpretation and the estimator ceiling**

1. **Biological**: these fine-mapped common variants (MAF > 0.5–1%) are predominantly ancient and pre-Out-of-Africa (median ~382 kyr; 63% ≥ 200 kyr; ≤5% younger than the ~65 kyr OOA horizon). Sustained presence at common frequency over hundreds of thousands of years is consistent with maintenance under purifying selection and mutation–selection balance acting on ancestrally shared standing variation, rather than recent positive selection or a post-OOA soft-sweep origin.

2. **Methodological (estimator ceiling, reframed)**: the Atlas-of-Variant-Age² coalescent estimator (1000G Phase 3 + Simons Genome Diversity Project) is calibrated in the European demographic context and its precision degrades for the deepest coalescent times. Because the estimates are reported in generations and converted at 28.1 yr generation⁻¹, the apparent upper tail (max ≈ 3.25 Myr) sits near the estimator's resolution ceiling and should be read as a lower-confidence bound on the deepest ages, not a precise point estimate. The honest caveat is therefore preserved: these are EUR-coalescent Atlas ages with a deep-time precision ceiling — but the central tendency and the great majority of the distribution sit unambiguously in deep, pre-OOA time.

3. **Implication for cluster framing**: the k=3 cluster definition resolves the youngest (C0, median ~113 kyr), intermediate (C1, ~358 kyr), and oldest (C2, ~508 kyr) strata of one deeply ancestral, pre-OOA set (cluster medians from the primary cluster assignment; consistent with main-text Table 1). The "Young" label denotes the youngest stratum relative to the rest of the credible set, all of which is ancient; it does not denote recent (Holocene/post-OOA) origin. The GMM-included subset is on average ~22% older than the GMM-excluded subset (Supplementary Table 17), reinforcing that ascertainment, if anything, favours the older, regulatory-active end of this ancient distribution.

**Supplementary Table 22 | Baseline age and AFR-frequency contextualisation.**

To disentangle whether the deep Atlas-of-Variant-Age² range of PGC3 credible-set variants is driven by *ascertainment* (EUR-coalescent estimator behaviour) versus *signal* (a genuine within-set age gradient relative to HapMap3-average), and to confirm the predominantly pre-Out-of-Africa (pre-OOA, ~65 kyr), ancestrally shared origin of these variants, we performed two complementary baseline analyses. Throughout, Atlas-of-Variant-Age (GEVA `AgeMedian_Mut`, TGP source) values are reported in years after converting the native generation-scaled estimates at 28.1 yr generation⁻¹ (Wohns 2022; consistent with the main-text Methods and Limitations).

**Section A. Atlas age vs. MAF + baseline-L2-matched HapMap3 control SNPs**

**Methods**: For each PGC3 credible-set variant (n = 20,637), 99.5 matched HapMap3 SNPs were drawn from a precomputed matched-control table (686,081 unique HapMap3 controls, matched on per-bin baseline-L2 score and EUR allele frequency). Atlas of Variant Age (TGP source) was looked up for both PGC3 and controls, with native generation-scaled `AgeMedian_Mut` values converted to years (×28.1); pooled and per-MAF-decile, per-cluster comparison tests were applied.

**Pooled distribution (years; ×28.1)**

| **Group** | **n** | **min (yr)** | **q01 (yr)** | **median (yr)** | **q99 (yr)** | **max (yr)** | **%<1.4 Myr** |
| --- | --- | --- | --- | --- | --- | --- | --- |
| PGC3 credible-set | 20,445 | 13,005 | 39,334 | **376,506** | 1,020,967 | 3,309,112 | 99.80% |
| MAF+L2-matched HapMap3 controls | 2,034,393 | 5,160 | 58,302 | **472,322** | 980,423 | 5,272,206 | 99.92% |

(The "%<1.4 Myr" column is the original 50,000-generation cutoff expressed in years: 50,000 gen × 28.1 = 1.405 Myr; the fractions are unchanged.)

**Tests**:

- Mann–Whitney U: *P* < 10⁻³⁰⁰ (PGC3 modestly younger within this deep window)

- Kolmogorov–Smirnov *D* = 0.137, *P* < 10⁻³⁰⁰

- Permutation test (one control per PGC3, 10,000 iterations, median diff): *P* = 0.78 (less directional under per-row resampling)

- Permutation test (pct < 1.4 Myr diff): *P* = 0.88

**MAF-decile stratified (years; ×28.1)**

| **Decile** | **MAF range** | **PGC median (yr)** | **Control median (yr)** | **Δ (yr)** | **MWU *P*** |
| --- | --- | --- | --- | --- | --- |
| 1 | 0.000–0.072 | 100,774 | 403,002 | **−302,228** | 1.0 × 10⁻²⁹⁰ |
| 2 | 0.072–0.149 | 254,092 | 414,798 | −160,706 | 2.0 × 10⁻⁵⁸ |
| 3 | 0.149–0.194 | 346,680 | 445,840 | −99,161 | 1.2 × 10⁻¹² |
| 4 | 0.194–0.206 | 481,280 | 462,683 | +18,597 | 0.75 (n.s.) |
| 5 | 0.206–0.264 | 378,610 | 464,791 | −86,181 | 3.8 × 10⁻¹⁴ |
| 6 | 0.264–0.311 | 430,058 | 480,867 | −50,809 | 7.2 × 10⁻²⁷ |
| 7 | 0.311–0.356 | 263,896 | 492,604 | **−228,708** | 5.3 × 10⁻¹⁰⁷ |
| 8 | 0.356–0.404 | 405,053 | 498,632 | −93,579 | 1.6 × 10⁻³³ |
| 9 | 0.404–0.463 | 422,566 | 504,833 | −82,267 | 6.4 × 10⁻⁴⁶ |
| 10 | 0.463–1.000 | 399,406 | 510,066 | −110,659 | 1.1 × 10⁻⁶⁶ |

Within this deep, predominantly pre-OOA window, PGC3 credible-set variants carry a modestly *younger* coalescent estimate than MAF-matched controls in 9 of 10 MAF deciles — a relative within-window gradient, not a difference between recent and ancient variation (all medians are hundreds of kyr).

**Per-cluster (vs MAF-matched controls; years, ×28.1)**

| **Cluster** | **n PGC** | **PGC median (yr)** | **PGC %<1.4 Myr** | **n CTL** | **CTL median (yr)** | **CTL %<1.4 Myr** | **Δ median (yr)** | **MWU *P*** |
| --- | --- | --- | --- | --- | --- | --- | --- | --- |
| **C0 (Young)** | 1,744 | **112,602** | 100.00% | 148,180 | 480,635 | 99.90% | **−368,033** | < 10⁻³⁰⁰ |
| C1 (Mid) | 316 | 357,995 | 97.15% | 26,628 | 487,403 | 99.92% | −129,408 | 1.7 × 10⁻⁹ |
| C2 (Old) | 2,854 | 517,003 | 99.68% | 211,879 | 483,890 | 99.88% | +33,113 | 2.8 × 10⁻⁶⁹ |

(Cluster medians correspond to main-text Table 1: C0 ≈ 113 kyr, C1 ≈ 358 kyr, C2 ≈ 508 kyr on the full credible-set distribution; the 517 kyr value above is the median of the C2 subset with available matched controls.)

**Interpretation**: All three clusters have median allele ages of ~113 kyr (C0) to ~517 kyr (C2) — i.e. predominantly pre-OOA (~65 kyr), deeply ancestral, with even the youngest cluster (C0) older than the Out-of-Africa bottleneck. The cluster ordering C0 < C1 < C2 is a real *within-window* age gradient (C0 ≈ 368 kyr younger by Atlas estimate than its MAF + L2-matched controls), not an ascertainment artefact, but "Young/Mid/Old" denote the youngest, intermediate and oldest strata of one deeply ancient set rather than recent versus ancient origins. The C2 cluster median is essentially indistinguishable from its matched controls (Δ = +33 kyr against a ~500 kyr baseline; statistically significant but absolutely small), consistent with C2 representing typically deep, ancestral variation.

**Section B. AFR allele-frequency baseline**

**Methods**: For each PGC3 credible-set variant (n = 20,637), we looked up African-American (PGC3 AFRAM, *n* = 5,998 cases + 3,826 controls = 9,824 total) allele frequency from the PGC3 wave-3 AFRAM sumstats. Variants matched in AFRAM: 19,733/20,637 (95.6%). The complement (904 variants, 4.4%) was absent from AFRAM (likely low imputation quality, reference mismatch, or genuine AFR absence) and was not classifiable.

**AFR partition classification**

| **Partition** | **Threshold** | **Overall** | **C0** | **C1** | **C2** |
| --- | --- | --- | --- | --- | --- |
| AFR_polymorphic | f_AFR ≥ 0.01 | 19,733 (95.6%) | 1,744 (99.9%) | 317 (100.0%) | 2,856 (100.0%) |
| AFR_rare | 0 < f_AFR < 0.01 | 0 | 0 | 0 | 0 |
| AFR_monomorphic | f_AFR = 0 | 0 | 0 | 0 | 0 |
| Absent from AFRAM | n/a | 904 (4.4%) | 1 | 0 | 0 |

**100% of cluster-assigned PGC3 credible-set variants are polymorphic in AFR ancestry**, with zero AFR-monomorphic (lineage-private post-OOA-derived) variants — consistent with an ancient, ancestrally shared, predominantly pre-OOA origin.

**AFR allele frequency distribution per cluster**

| **Cluster** | **n** | **median f_AFR** | **mean f_AFR** | **%≥1%** | **%≥5%** |
| --- | --- | --- | --- | --- | --- |
| **C0 (Young)** | 1,744 | **0.860** | 0.808 | 100% | 100% |
| C1 (Mid) | 317 | 0.370 | 0.528 | 100% | 100% |
| C2 (Old) | 2,856 | 0.490 | 0.496 | 100% | 100% |

**Section C. Interpretation**

The matched-control-age and AFR-allele-frequency baselines together confirm the deep, predominantly pre-OOA, ancestrally shared origin of the credible-set variants:

1. **The cluster-level age stratification (C0 < C1 < C2 by Atlas coalescent age) is a real within-window gradient, not an ascertainment artefact** — C0 variants are ~368 kyr younger by Atlas estimate than MAF + L2-matched HapMap3 controls, yet still ~113 kyr old (pre-OOA) in absolute terms.

2. **The variants are ancestrally shared, not lineage-private** — every cluster-assigned PGC3 credible-set variant is polymorphic in AFR ancestry, with the Young cluster (C0) most so (median f_AFR = 0.86, common in AFR), as expected for ancient pre-OOA variation.

3. **The within-set age ordering is an EUR-coalescent-axis property, consistent with deep shared TMRCA** — even C0, the youngest stratum (Atlas EUR-coalescent median ≈ 113 kyr), is AFR-common (f_AFR = 0.86). The EUR-coalescent (GEVA Atlas) estimator is expected to differ from an African-lineage genealogy at these AFR-polymorphic loci: the Wohns 2022 unified tree-sequence African-lineage median for C0 is ≈ 311 kyr (Fig 5), older still than the Atlas estimate. Both estimators place the variants deep in the pre-OOA past; the Atlas EUR-coalescent axis indexes the youngest-to-oldest *strata* of this ancestral set, not recent post-OOA origins.

4. **Implication for paper framing**: The three cluster identities (Young/Mid/Old) are the youngest, intermediate and oldest strata of a single, deeply ancient, predominantly pre-OOA, ancestrally shared set of SCZ credible-set variants. Their persistence at appreciable frequency is consistent with purifying selection and mutation–selection balance acting on long-standing ancestral variation, not with recent or post-OOA soft sweeps. This interpretation is used in the main-text Methods (allele-age coverage and generation-to-year conversion), Results (¶29/¶47) and Discussion (non-antagonistic pleiotropy framing), and is consistent with main-text Table 1 and Fig 5.
