## Supplementary Methods for "Common schizophrenia heritability concentrates in an evolutionarily young, brain-regulatory subset of fine-mapped credible-set variants"

### Contents

Supplementary Note 1: Variant substrate and coverage
- 1.1 chrX exclusion
- 1.2 MAPT-excluded primary substrate
- 1.3 Alternative coalescent frameworks
- 1.4 MAF- and LD-matched control baseline
- 1.5 Neighbour-control comparison
- 1.6 Long-range LD region masking (Price et al. 2008)

Supplementary Note 2: Feature derivation
- 2.1 eQTL data
- 2.2 Tissue-power rescaling
- 2.3 Brain-specificity 4-test design

Supplementary Note 3: Within-locus correlation framework
- 3.1 Within-locus partial-rank residualisation
- 3.2 Block bootstrap confidence intervals

Supplementary Note 4: Mixture modelling and cluster sensitivity
- 4.1 GMM component selection
- 4.2 MAPT-included sensitivity
- 4.3 Block-jackknife (Tashman) sensitivity
- 4.4 Component-number (k) sensitivity
- 4.5 Feature-space (2D) sensitivity

Supplementary Note 5: Partitioned LD score regression
- 5.1 LDSC Python-3 patches
- 5.2 LDSC alternative frameworks
- 5.3 Empirical permutation null

Supplementary Note 6: Cross-trait analyses and disease-specific munge implementations
- 6.1 Daner-aware munge
- 6.2 Cross-disorder munge implementations
- 6.3 GIANT-aware munge
- 6.4 r_g-prediction

Supplementary Note 7: Robustness and sensitivity battery
- 7.1 Block-jackknife (Tashman) axis
- 7.2 Feature-space (2D) axis
- 7.3 MAPT-included axis
- 7.4 Brain-specificity 4-test axis
- 7.5 Tissue-power-matched brain-specificity axis
- 7.6 Empirical matched-LD-MAF permutation null axis
- 7.7 Component-number (k) axis

### Supplementary Note 1: Variant substrate and coverage

#### 1.1 chrX exclusion

The PGC3 schizophrenia GWAS^6^ reports five chromosome X loci among its 287 genome-wide significant loci. We excluded the chrX loci from the present analyses in line with standard GWAS practice^84^, on three considerations. First, the Atlas of Variant Age coalescent estimator^19^ provides per-variant age estimates derived from autosomal recombination and coalescent dynamics; chrX has a substantially different effective population size (3/4 of the autosomal *N*_e_ under random mating) and a hemizygous-male-driven recombination map, so chrX age estimates are not directly comparable to autosomal estimates in the Atlas framework. Second, the partitioned LDSC **baseline-LD v2.2 annotation set^27^** is derived for autosomal SNPs; LDSC regression weights and LD-score inputs for chrX require a separately constructed XSC-LDSC reference panel that is not part of the standard partitioned-heritability pipeline. Third, partitioned heritability on chrX would confound sex-stratified architecture with chromosome-specific dynamics in ways that are not resolvable at the present sample sizes. The exclusion reduces the variant substrate from the headline 20,766 to 20,637 autosomal credible-set variants, which forms the basis for all analyses reported throughout.

#### 1.2 MAPT-excluded primary substrate

The 17q21.31 MAPT inversion^56^ is a ~3-Mb structural-variant locus represented in PGC3 fine-mapping by a single credible set (CS_224) of 1,742 variants. We excluded this locus from the primary clustering substrate on three grounds. First, the single-credible-set structure violates the per-credible-set rank-residualisation framework used for within-locus partial-rank correlations (Supplementary Methods 3.1). Second, its long-range LD architecture, suppressed recombination and elevated |iHS| are not representative of the remaining 249 fine-mapped loci that contributed credible sets with ≥ 5 variants. Third, including a single 1,742-variant credible set would inflate any cluster-level estimator at this locus relative to the per-credible-set sampling design used elsewhere. The held-out reintegration test in Supplementary Methods 4.2 confirms that the principal Young-cluster heritability concentration is robust to this exclusion choice (47.4 → 47.8-fold; Supplementary Table 18).

#### 1.3 Alternative coalescent frameworks

Four alternative variant-level coalescent estimators were considered as primary substrates for allele age and not adopted, with the following per-method rationale. Relate^85^ infers tree sequences from phased haplotypes and provides allele-age estimates with relatively short runtime, but its output is a sample of trees under the SMC’ approximation rather than a coalescent-fitted GEVA-style estimator, and its age uncertainty for variants outside the 1000G reference panel is not well characterised at the variant level required here. ARGweaver^86^ provides full ancestral recombination graph inference but is computationally prohibitive at the genome scale required for 20,766 credible-set variants. tsinfer^20^ constructs a non-genealogical succinct tree sequence and is well suited to very large samples but does not produce calibrated coalescent age estimates compatible with the Atlas of Variant Age frame used elsewhere in the analyses. The Wohns et al.^21^ unified tree-sequence framework is the natural successor to the Atlas of Variant Age and supports cross-population coalescent dating at variant resolution; it is the method we identify in the main text as required to definitively partition pre-OOA anatomically modern human-shared from European-private post-OOA-derived alleles. Integrating the Wohns 2022 outputs at credible-set variant resolution requires polarity-aware cross-population reconciliation, which falls outside the analytical scope of the present study. The Atlas of Variant Age GEVA estimator^19^ was therefore retained as the primary per-variant age annotation, with the EUR-coalescent reference-frame caveat addressed by the MAF- and LD-matched HapMap3 baseline analysis (Supplementary Methods 1.4; Supplementary Table 17) and the within-window neighbour-control comparison (Supplementary Methods 1.5).

#### 1.4 MAF-and-LD-matched control baseline

To distinguish ascertainment-driven age compression (operating uniformly within the Atlas estimator’s EUR-coalescent reference frame) from signal-driven age compression (PGC3 credible-set variants genuinely younger than the broader genome), we constructed a MAF- and LD-matched HapMap3 control baseline. For each PGC3 credible-set variant (*n* = 20,637 autosomal), 99.5 matched HapMap3 SNPs were drawn on average from a precomputed matched-control table containing 686,081 unique HapMap3 controls matched on per-bin baseline-L2 score and EUR allele frequency. The per-focal-variant distribution of matched-control counts (median, interquartile range, minimum, and the proportion of focal variants matched to at least 10 controls), together with the per-bin coverage of the 686,081-SNP control table across the 10 × 10 MAF–L2 grid, is reported in Supplementary Table 17 section D. The construction proceeded in five steps. (i) PGC3 credible-set variants were binned by allele frequency (10 deciles) and baseline-L2 score (10 deciles), yielding 100 joint bins. (ii) For each PGC3 bin cell, HapMap3 SNPs with matching MAF and baseline-L2 were sampled to populate the matched-control table. (iii) Atlas of Variant Age estimates were looked up for PGC3 credible-set variants and their matched HapMap3 controls. (iv) Two comparison families were applied: pooled Mann–Whitney *U* and Kolmogorov–Smirnov tests on the full PGC3 versus full matched control set, and decile-stratified Mann–Whitney *U* tests within each MAF bin. (v) Per-cluster comparisons (C0, C1, C2 against cluster-specific matched controls) repeated the test at cluster resolution. The full set of test statistics, pooled distribution percentiles, MAF-decile-stratified results and per-cluster comparisons against MAF-matched controls is reported in Supplementary Table 17 (sections A, B, C). The construction confirms that the cluster-level age stratification is signal-driven rather than ascertainment-driven within the EUR-coalescent reference frame.

#### 1.5 Neighbour-control comparison

A within-window neighbour-control test was applied as a complementary negative control to the MAF-and-LD-matched HapMap3 baseline (Supplementary Methods 1.4), to address whether the cluster-level age stratification could be reconstructed from neutral allele-frequency-spectrum drift within the immediate physical neighbourhood of each PGC3 credible-set variant. The procedure proceeded in five steps. (i) For each PGC3 credible-set variant outside the MAPT inversion (*n* = 18,895), neighbouring single-nucleotide variants from 1000 Genomes EUR phased haplotypes^57^ within ± 50 kb of the focal variant were enumerated. (ii) Within the ± 50 kb window, candidate neighbour controls were further filtered to within one EUR-MAF decile of the focal variant, to remove residual MAF–age confounding inside the window. (iii) Up to ten neighbour controls per focal variant were drawn at random without replacement from the in-window MAF-matched pool. (iv) Atlas of Variant Age estimates^19^ were looked up for both the focal variant and each neighbour control. (v) Paired Wilcoxon signed-rank tests were applied between each focal variant and the median age of its neighbour-control set. The procedure differs from the MAF-and-LD-matched HapMap3 baseline (Supplementary Methods 1.4) in two ways: (a) it constrains the control set to within ± 50 kb of the focal variant, removing genome-wide drift heterogeneity at the cost of smaller within-window MAF-matching pools, and (b) it uses a within-pair paired test rather than a pooled Mann–Whitney *U* test, which is more powerful when within-pair concordance is high. The signed-rank statistic, paired sample size, median focal-minus-neighbour age difference, and two-sided *P*-value for both the pooled and per-cluster (C0, C1, C2) neighbour-control comparisons are reported in Supplementary Table 17 section E; the test rejects the null of equal focal-versus-neighbour age distributions, corroborating the Supplementary Methods 1.4 finding that the cluster-level age stratification is signal-driven rather than ascertainment-driven, and additionally rules out neutral-drift heterogeneity at the within-window scale.

#### 1.6 Long-range LD region masking (Price et al. 2008)

### To distinguish the principal partitioned-heritability finding from residual long-range LD (LR-LD) confounding, the 24 canonical LR-LD regions catalogued in GRCh37 coordinates by Price et al. 2008 (Am J Hum Genet 83:132–135; Supplementary Table 6 section A, with three canonical corrections for boundary precision per the LDSC documentation) were masked from each per-chromosome cluster annotation. Per-cluster masked variant counts are reported in Supplementary Table 6 section B. LD scores were recomputed under the masked annotations using the standard --l2 --bfile --annot --print-snps invocation against 1000 Genomes Phase 3 plink bfiles with --ld-wind-cm 1, and partitioned heritability was re-run on PGC3 EUR schizophrenia (Supplementary Table 6 section C) and on Wightman 2021 Alzheimer's disease (Supplementary Table 6 section D, negative-control trait) under the baseline-LD v2.2 model. The principal Young-cluster heritability concentration is preserved after LR-LD masking on the SCZ substrate and remains null on the AD substrate, confirming that the cluster-level signal is not an LR-LD artefact.

### Supplementary Note 2: Feature derivation

#### 2.1 eQTL data

Three *cis*-eQTL data sources were integrated. The primary tissue-level annotation was the GTEx v10 fine-mapped per-tissue significant variant–gene pairs (GTEx Portal v10 data release; cohort and analytical pipeline per GTEx Consortium^60^), processed by computing the minimum nominal *P*-value per PGC3 credible-set variant across all tested genes within each of 13 brain tissues (Amygdala, Anterior cingulate cortex, Caudate basal ganglia, Cerebellar hemisphere, Cerebellum, Cortex, Frontal cortex BA9, Hippocampus, Hypothalamus, Nucleus accumbens basal ganglia, Putamen basal ganglia, Spinal cord cervical c-1, Substantia nigra) and three blood/immune tissues (Whole blood, Spleen, EBV-transformed lymphocytes). Variants were lifted from GRCh37 to GRCh38 using the UCSC liftOver pipeline; allele orientation was reconciled across four branches (forward, flip, forward-rev-comp, flip-rev-comp), with palindromic A/T and C/G hits retained but flagged. Brain–blood specificity was derived per variant as *b*_spec_ = −log_10_(*P*_brain,min_) / [−log_10_(*P*_brain,min_) + −log_10_(*P*_blood,min_)]. This ratio specification was selected over unbounded alternatives (such as brain-minus-blood −log_10_ *P* differences or tau-style specificity indices) because it is bounded in [0, 1], monotone in the brain–blood −log_10_ *P* ratio, and behaves stably when only one tissue family achieves significance (the denominator absorbs joint magnitude, so the ratio remains defined and interpretable). Variants for which neither brain nor blood cis-eQTL achieved nominal significance (both per-tissue minimum P > 0.05) have undefined b_spec under this formulation and are flagged as missing; such variants are excluded from b_spec-dependent analyses (the within-locus b_spec × age correlation in Supplementary Methods 3.1 and the 3D GMM input in Supplementary Methods 4.1). The single-cell brain layer used Bryois et al.^34^ single-cell brain *cis*-eQTL summary statistics across eight cell types (astrocytes, endothelial cells, excitatory neurons, inhibitory neurons, microglia, oligodendrocytes, oligodendrocyte progenitor cells, pericytes); per-cell-type per-rsID minimum *P*-values were extracted, achieving 89.2% rsID coverage of the 18,895 non-MAPT autosomal credible-set variants (cell-type-specific coverages range 77.2 to 96.7%; Supplementary Table 7 section B). Per-cell-type cluster-vs-rest coverage comparisons used a three-way (C0 vs C1 vs C2) Pearson χ² test (2 d.f.) without Yates continuity correction, justified by all per-cell expected counts ≥ 5; eight-test Benjamini–Hochberg FDR was applied across the eight Bryois cell types. A secondary cross-atlas consistency check used the PsychENCODE 2 atlas^61^ at the bulk level. Cluster-specific pathway enrichment (Supplementary Table 8 section A) was computed by uploading per-cluster nearest-gene lists to the Enrichr API (https://maayanlab.cloud/Enrichr/) against four library snapshots — GO Biological Process 2023, KEGG 2021 Human, Reactome 2022 and MSigDB Hallmark 2020 — with the union of nearest-gene assignments across all PGC3 credible-set variants as the background gene universe; per-term hypergeometric P-values returned by Enrichr were Benjamini–Hochberg FDR-adjusted within each library.

#### 2.2 Tissue-power rescaling

GTEx v10 tissue donor sample sizes differ substantially across tissues, with Whole Blood at *N* ≈ 838 versus brain tissues at *N* ≈ 144–276. For an identical effect-size β, the eQTL test statistic Z = β · √N scales with √N, so tissues with larger sample sizes will yield more significant nominal *P*-values for the same underlying biological effect. To check whether the brain–blood specificity ranking is confounded by tissue-power asymmetry, we reconstructed |Z| from the observed minimum nominal *P*-value per tissue using the standard relation Z = Φ⁻¹(1 − *P*/2) and rescaled to a reference sample size *N*_ref_ = 220, the median brain-tissue donor *N*. The rescaled |Z| values were then re-converted to *P*-values, and the brain–blood specificity score *b*_spec_ was recomputed using the power-matched per-tissue significance levels. A 3D GMM was re-fit on the power-matched feature space, and partitioned S-LDSC was applied to the resulting C0 annotation under the baseline-LD v2.2 model. Power-matching strengthens rather than weakens the principal cluster-level findings, with **C0 enrichment rising from 47.4- to 71.9-fold** and the **C2 (Old) cluster also rising to 36.6-fold (*P* = 2.7 × 10⁻³, Bonferroni-significant)** (Supplementary Table 19B/C). The principal heritability-concentration finding is therefore not driven by GTEx tissue-power asymmetry.

#### 2.3 Brain-specificity 4-test design

To adjudicate whether the joint 3D Young-cluster definition is redundant with brain–blood regulatory specificity ranking alone, four S-LDSC tests were conducted on PGC3 EUR schizophrenia under the baseline-LD v2.2 model. The four tests are: (A) brain_spec-only single-annotation S-LDSC partition, using a top-1742 b_spec-thresholded binary annotation (the 1,742 highest-b_spec non-MAPT credible-set variants, size-matched to C0 in the baseline-LD v2.2 LDSC pool); (B) brain_spec continuous-annotation single S-LDSC partition; (C) C0-only single-annotation S-LDSC partition, using the binary C0 cluster-membership indicator; and (D) joint conditional S-LDSC model containing both brain_spec_top and C0_only annotations alongside the baseline-LD v2.2 categories, in which each annotation’s conditional coefficient *Z*-score is evaluated against the other.

Under baseline-LD v2.2 (Supplementary Table 19A), the four tests yielded the following enrichments and conditional *Z*-scores: (A) brain_spec_top single-annotation enrichment was 54.5-fold (*P* = 3.2 × 10⁻⁸, *Z* = +5.67); (B) brain_spec continuous single-annotation enrichment was 39.4-fold (*P* = 1.1 × 10⁻¹³, *Z* = +7.82); (C) C0-only single-annotation enrichment was 100.2-fold (*P* = 4.6 × 10⁻⁸, *Z* = +5.61); and (D) the joint conditional model assigned the principal signal to C0, with C0 conditional *Z* = +4.80 (*P* = 1.7 × 10⁻⁸, Bonferroni-significant) versus brain_spec_top conditional *Z* = +1.26 (not significant).

The brain_spec ranking is therefore largely collinear with, and absorbed by, the C0 cluster in a joint conditional model under the baseline-LD v2.2 model.

### Supplementary Note 3: Within-locus correlation framework

#### 3.1 Within-locus partial-rank residualisation

The within-locus partial rank correlation procedure operates inside each fine-mapped credible set with at least five variants, to remove cross-locus heterogeneity in MAF, baseline-LD architecture and variant density that would otherwise confound a pooled-variant rank correlation. The procedure proceeds in five steps. (i) Within each credible set of size *m* ≥ 5, the two focal features *x* and *y* (for example brain–blood specificity and log₁₀ allele age) are converted to within-credible-set ranks. (ii) When MAF residualisation is requested, the within-credible-set MAF rank is computed in parallel. (iii) The within-locus ranks of *x* and *y* are residualised against the within-locus MAF rank by ordinary-least-squares regression: *x*_resid_ = *x*_rank_ − β_x_ · MAF_rank_; *y*_resid_ = *y*_rank_ − β_y_ · MAF_rank_. (iv) Spearman correlations between the residualised ranks are computed per credible set. (v) Per-credible-set Spearman ρ values are pooled across the 249 non-MAPT fine-mapped loci by inverse-variance weighting under the small-sample Spearman variance approximation Var(ρ̂) ≈ (1 − ρ̂²)² / (*m* − 1). The within-locus rank specification (steps i and ii) removes between-locus heterogeneity in the marginal distributions of *x* and *y*; the MAF-residualisation step (iii) removes within-locus MAF–feature confounding that arises because MAF affects both variant ascertainment and the precision of *cis*-eQTL discovery. Pooled estimates with and without MAF residualisation are reported in parallel throughout (Supplementary Table 2): the no-residualisation specification preserves the natural within-locus rank, while the MAF-residualised specification removes any pooled correlation that could be reconstructed from MAF alone. Pearson correlations on the residualised ranks are reported for cross-method consistency where indicated.

#### 3.2 Block bootstrap confidence intervals

Bootstrap 95% confidence intervals for the within-locus partial rank correlation are computed using a per-locus block bootstrap with inside-loop recomputation of the full residualisation pipeline. The choice of block bootstrap at the credible-set level, rather than per-SNP bootstrap, reflects the per-locus sampling design of the fine-mapping framework: the credible set is the indivisible unit of within-locus inference, and SNP-level resampling would understate uncertainty by treating LD-linked variants within a credible set as independent. The procedure proceeds in four steps. (i) The 249 non-MAPT fine-mapped credible sets with *m* ≥ 5 variants are resampled with replacement to produce a bootstrap dataset of identical credible-set count. (ii) Within each resampled credible set, the full within-locus rank conversion, MAF-rank residualisation and per-credible-set Spearman ρ computation (Supplementary Methods 3.1) are recomputed from scratch inside the bootstrap iteration. The recomputation step is critical: residualising once on the original data and then resampling the per-locus ρ values would understate uncertainty by treating the residualisation coefficients β_x_ and β_y_ as fixed across iterations. (iii) The pooled inverse-variance-weighted ρ̂_boot_ is computed within the iteration. (iv) Steps (i) to (iii) are repeated 1,000 times to produce the bootstrap distribution; 95% percentile confidence intervals are reported. The percentile interval was selected over the bias-corrected accelerated (BCa) interval because the Spearman ρ̂ distribution within iteration was approximately symmetric in all cases inspected, and the BCa acceleration term is unstable when the per-iteration sample sizes are non-uniform under credible-set resampling. Inside-iteration block-bootstrap CIs are wider than the corresponding asymptotic Spearman CIs, reflecting honest uncertainty from per-locus resampling with within-iteration recomputation; CIs that cross zero on the residualised specification reflect genuine uncertainty at the per-locus partial-rank level.

### Supplementary Note 4: Mixture modelling and cluster sensitivity

#### 4.1 GMM component selection

Six clustering-fit metrics were computed at *k* = 1–6 on the standardised 3D joint distribution of log₁₀ allele age, brain–blood specificity, and Voight-corrected |iHS| (per-variant |iHS| standardised to mean 0, SD 1 within 20 derived-allele-frequency bins per Voight et al. 2006, scikit-allel v1.3.13 implementation) (*n* = 4,918 non-MAPT PGC3 credible-set variants with all three features). The full battery comprised: Bayesian Information Criterion (BIC), Akaike Information Criterion (AIC), Integrated Completed Likelihood (ICL), silhouette coefficient, Calinski–Harabasz index, and Davies–Bouldin index. BIC and AIC favoured larger *k* monotonically across *k* = 1–6, a documented over-fitting tendency in low-sample Gaussian mixture estimation^65^. The three clustering-specific metrics designed to penalise over-fitting (ICL, silhouette, Davies–Bouldin) independently converged on *k* = 3 as the local optimum. The choice of *k* = 3 was adopted on parsimony and interpretability grounds, supported by the three converging metrics and by the a priori three-regime hypothesis specified before clustering (Young / Mid / Old). The 1,000-bootstrap cluster-membership stability analysis at *k* = 3 yielded a bootstrap mean adjusted Rand index of 0.967 ± 0.019 in the primary 3D feature space, with a 95% percentile interval of [0.919, 0.990], confirming that the *k* = 3 partition is highly stable under resampling. *k*-sensitivity at *k* ∈ {2, 3, 4} (Supplementary Methods 4.4; Supplementary Table 16) confirms that the principal Young-cluster heritability concentration is significant at all three values under the baseline-LD v2.2 model. GMM fits used scikit-learn GaussianMixture with covariance_type='full', n_init = 10 random initialisations per k, random_state = 42, and the sklearn EM defaults (max_iter = 100, tol = 1e-3, reg_covar = 1e-6, k-means++ initialisation).

#### 4.2 MAPT-included sensitivity

To test whether the principal Young-cluster heritability concentration is robust to the MAPT-exclusion choice (rationale in Supplementary Methods 1.2), a held-out prediction protocol was used. The primary trained 3D Gaussian mixture model and StandardScaler (fit on the non-MAPT training set of *n* = 4,918 credible-set variants with all three features) were applied to the 1,568 MAPT variants with all three features without re-training, preserving the primary cluster definition. MAPT variants partitioned 117 to C0, 103 to C1 and 1,348 to C2; the C2-dominant distribution is expected for an ancient inversion locus. Cluster annotation files were rebuilt with MAPT variants assigned to their predicted clusters. Only chromosome 17 LD scores required recomputation (HapMap3 restriction, 32,219 chr17 SNPs); the other 21 chromosomes were byte-identical to the MAPT-excluded primary annotation. Partitioned LDSC was re-run on PGC3 EUR schizophrenia with the baseline-LD v2.2 model. The principal C0 enrichment is essentially unchanged (47.4 → 47.8-fold; conditional *Z* +3.05 → +3.12; Supplementary Table 18); the C1 enrichment rises modestly (13.0 → 17.3-fold, *Z* = +3.34), as the 103 high-|iHS| MAPT variants assigned to C1 carry additional heritability mass; the C2 enrichment falls (15.6 → 8.5-fold, *Z* = +0.69), as the 1,348 MAPT variants assigned to C2 sit under extended long-range LD across the inversion, diluting C2’s per-SNP heritability density. Total observed-scale *h*² (0.328 ± 0.012; mean χ² = 2.02; intercept 1.094 ± 0.013) is identical to the primary partition. The Young-cluster (C0) heritability concentration is therefore not driven by the MAPT-exclusion methodological choice; by contrast, the MAPT-included C2 conditional *Z* = +0.69 does not reach Bonferroni significance under the baseline-LD v2.2 family, so any Bonferroni-significant C2 enrichment reported in the MAPT-excluded specification (or in the tissue-power-matched substrate; Supplementary Methods 2.2) is dependent on the MAPT-exclusion choice and is reported with that caveat throughout the manuscript.

#### 4.3 Block-jackknife (Tashman) sensitivity

LDSC standard errors are estimated by a block-jackknife with a default of 200 blocks in v1.0.1; Tashman et al.^35^ flagged a small-annotation type-1-error inflation in this default setting and recommended a higher block count for small annotations. We re-ran the principal partitioned LDSC for PGC3 EUR schizophrenia with --n-blocks 1000 (five-fold the default) under the baseline-LD v2.2 model, confirming point-estimate identity with the default-blocks specification (47.4-fold, *Z* = +3.20 versus default *Z* = +3.05) and a marginally improved standard error (Supplementary Table 15). The Tashman concern is therefore not load-bearing for the present cluster-level findings, and is further addressed by the empirical matched-LD-MAF permutation null (Supplementary Methods 5.3; Supplementary Table 20). The --n-blocks 1000 specification was applied only to this principal partition; the seven robustness-battery axes (Supplementary Methods 7) retained the LDSC v1.0.1 default of --n-blocks 200.

#### 4.4 Component-number (k) sensitivity

To verify that the principal Young-cluster heritability concentration is not an artefact of the *k* = 3 choice, we re-fit the GMM on the same 3D feature space at *k* = 2 and *k* = 4 and ran identical partitioned LDSC under the baseline-LD v2.2 model. Cluster centroids are stable across *k* ∈ {2, 3, 4}, and the Young-cluster heritability concentration is significant at all three values (*k* = 2 single-annotation 96.8-fold, *Z* = +5.28; *k* = 3 multi-annotation 47.4-fold, *Z* = +3.05; *k* = 4 single-annotation 105.3-fold, *Z* = +5.77; Supplementary Table 16). The *k* = 3 partition was retained for the primary analyses on the parsimony and three-converging-metric grounds set out in Supplementary Methods 4.1.

#### 4.5 Feature-space (2D) sensitivity

To test the robustness of the cluster partition to dropping the brain–blood specificity dimension, we re-fit the GMM on only two evolutionary features (log₁₀ allele age and |iHS|), which raised GMM-input coverage from 26.0% to 83.3% of non-MAPT credible-set variants. Identical partitioned LDSC was applied to the 2D-derived cluster annotations under the baseline-LD v2.2 model (Supplementary Table 14). The 2D Young cluster shows attenuated heritability concentration (C0 30.3-fold, *P* = 0.10; C2 48.2-fold Bonferroni-significant, *Z* = +4.10), confirming that the principal 3D Young-cluster signal requires the joint feature combination and that the brain-blood regulatory specificity dimension carries non-redundant signal.

### Supplementary Note 5: Partitioned LD score regression

#### 5.1 LDSC Python-3 patches

The LDSC v1.0.1 reference implementation^66,67^ was distributed as Python 2.7 code at the time of release. To run within the present Python 3.11 environment, seven syntactic patches were applied. These patches are strictly syntactic and do not change the statistical procedure or any numerical input to the regression. The patches are: (i) bare-form print statements converted to print() function calls; (ii) xrange replaced with range; (iii) the integer-division operator changed from / to // in counting expressions; (iv) dict.iteritems() replaced with dict.items(); (v) explicit np.int64 casts inserted in regression-index expressions where Python-2 implicit-integer division had returned integers (Python-3 true division returns floats and breaks index arithmetic); (vi) bz2.BZ2File(path, 'r') replaced with bz2.open(path, 'rt') for text-mode reads of compressed .l2.ldscore.gz files; and (vii) the print >> sys.stderr redirection form replaced with the file=sys.stderr keyword argument of print(). All seven patches were verified against the LDSC v1.0.1 Python-2 reference by re-running the LDSC test suite under both Python 2.7 and Python 3.11 with the patched code, yielding identical regression coefficients and standard errors to floating-point precision; the patched fork is archived under the Code Availability section. The patches do not include the Tashman et al.^35^ small-annotation type-1-error mitigation; that concern is addressed separately via the --n-blocks 1000 sensitivity (Supplementary Methods 4.3; Supplementary Table 15) and the empirical matched-LD-MAF permutation null (Supplementary Methods 5.3; Supplementary Table 20). The patched LDSC was invoked with the standard 1000 Genomes Phase 3 reference triad — --ref-ld-chr baseline-LD v2.2 (Gazal et al.27 release; Zenodo 10515792), --w-ld-chr 1000G_Phase3_weights_hm3_no_MHC, --frqfile-chr 1000G_Phase3_frq (1000G.EUR.QC. for EUR analyses; 1000G.EAS.QC. for EAS analyses) — with --overlap-annot --print-coefficients on all partitioned heritability fits and --ld-wind-cm 1 for per-annotation L2 computation against 1000G Phase 3 plink bfiles.

#### 5.2 LDSC alternative frameworks

Five alternative heritability-, annotation-partitioning, or causal-gene frameworks were considered and not applied in the present study, with the following rationale. PolyFun^71^ extends partitioned LDSC with functional priors and is well suited to per-variant fine-mapping, but the present primary inference is on cluster-level heritability concentration rather than per-variant causal posterior. MAGMA^72^ provides gene-level association statistics by aggregating variant-level statistics; it is complementary to but does not replace SNP-level partitioned heritability inference. sc-linker^73^ extends partitioned LDSC to single-cell-derived annotations using regulatory-element overlap; the present cell-type analyses use Bryois et al.^34^ per-cell-type *cis*-eQTL coverage at variant resolution (Supplementary Table 7) rather than sc-linker annotations. Formal colocalisation tests (e.g., COLOC^62^) and SMR/HEIDI^63^ both provide per-locus causal-gene assignment from eQTL substrates, via posterior shared-causal-variant inference and Mendelian randomisation respectively; neither was applied here because the primary inference is on heritability concentration within cluster annotations rather than per-locus causal-gene assignment. These frameworks remain available substrates for follow-up studies layered on top of the cluster assignments and per-variant feature tables reported here (Supplementary Tables 5, 7, 17).

#### 5.3 Empirical permutation null

We constructed an empirical matched-LD-MAF permutation null to assess whether the observed C0 conditional *Z* = +3.05 (under baseline-LD v2.2) lies outside the distribution expected under matched-LD-MAF random annotation sampling. The construction proceeded in six steps. (i) Candidate control SNPs were drawn from the full baseline-LD v2.2 annotation set intersected with the 1000 Genomes EUR allele-frequency file (9,997,231 SNPs), ensuring full coverage of all 1,744 C0 credible-set variants. Positional nearest-neighbour LD-score imputation was applied to non-HM3 variants in the expanded pool, since the baseline-LD .l2.ldscore.gz files distributed by Gazal et al.^27^ are restricted to HM3 SNPs; for each non-HM3 SNP the LD-score of the nearest HM3 SNP by physical position was assigned. (ii) The pool was partitioned into a decile L2-bin × decile MAF-bin grid (10 × 10 = 100 bins). (iii) The bin distribution of the primary C0 SNPs (*n* = 1,744 in the baseline-LD v2.2 intersection of the 9,997,231-SNP pool; 1 SNP fewer than the 1,745 GMM-assigned C0 variants in the master annotation) was computed across the 100 bins. (iv) *N* = 1,000 control SNP sets matched per bin to the C0 bin distribution were drawn from the expanded pool without replacement, and per-chromosome cluster-annotation files were constructed and matched to the EUR baseline-LD v2.2 SNP set. (v) Partitioned LDSC was run on each of the 1,000 control annotations using the same baseline-LD v2.2 model and PGC3 EUR SCZ summary statistics as the primary analysis, yielding 1,000 null conditional *Z*-scores. (vi) The observed C0 conditional *Z* = +3.05 was compared with the empirical null distribution. Across the 1,000 permutations the null *Z* distribution had mean ± SD −0.08 ± 0.98 (range [−3.69, +2.64]) and null enrichment mean ± SD +0.7-fold ± 17.2-fold; 0/1,000 controls reached the observed *Z* ≥ +3.05, placing the observed *Z* 3.18 SD above the null mean. Two complementary summaries of the empirical null are reported. Under the non-parametric (R + 1)/(N + 1) one-sided upper bound the empirical *P* = 1/1001 ≈ 1.0 × 10⁻³, the smallest value attainable with 1,000 permutations; under a parametric Gaussian approximation to the null moments (mean −0.08, SD 0.98) the observed *Z* corresponds to a one-sided *P* ≈ 7.4 × 10⁻⁴. The empirical null is reported as a matched-architecture sanity check on the primary baseline-LD v2.2 multi-annotation conditional inference; significance of the C0 partitioned-heritability concentration is established by the primary partitioned-LDSC conditional *Z* = +3.05 against the baseline-LD v2.2 Bonferroni threshold, not by the permutation null. Full per-permutation output is reported in Supplementary Table 20. The 1,000-permutation null required full per-chromosome LD-score recomputation across 22 chromosomes plus a baseline-LD v2.2 partitioned-LDSC fit on the corrected 9,997,231-SNP pool with nearest-HM3 LD-imputation for each permutation, executed as a SLURM array on the TRUBA HPC cluster; at N = 1,000 the (R + 1)/(N + 1) bound resolves to 1.0 × 10⁻³, consistent with the parametric Gaussian one-sided P ≈ 7.4 × 10⁻⁴.

### Supplementary Note 6: Cross-trait analyses and disease-specific munge implementations

#### 6.1 Daner-aware munge

PGC sex-stratified summary statistics are distributed in the daner format, which differs from the canonical LDSC munge_sumstats.py expected schema in column naming and effective-sample-size encoding. To preserve effective sample size and effect-allele orientation through munge, we used a daner-aware adapter. The adapter performs four operations: (i) column renaming from daner (CHR, BP, SNP, A1, A2, FRQ_A_, FRQ_U_, INFO, OR, SE, P) to the LDSC-expected schema (SNP, A1, A2, N, Z, INFO); (ii) effective-sample-size reconstruction per row using *N*_eff_ = 4 · *N*_cas_ · *N*_con_ / (*N*_cas_ + *N*_con_), where *N*_cas_ and *N*_con_ are extracted from the daner header; (iii) Z-score derivation from the log-odds-ratio and standard error as Z = log(OR) / SE, preserving sign on the A1 allele; (iv) palindromic A/T and C/G pre-filtering at the munge stage, followed by INFO ≥ 0.9 quality filtering and MAF ≥ 0.01 filtering against the HapMap3 reference SNP list. The adapter was applied to the four PGC3 wave 3 sex × ancestry strata: EUR male, EUR female, EAS male, EAS female. Effective sample sizes after munge were 68,003 (EUR male), 47,652 (EUR female), 13,017 (EAS male) and 13,163 (EAS female). Per-stratum partitioned heritability inference used the baseline-LD v2.2 conditional-Z Bonferroni threshold (|Z| ≥ 2.81 for 97 annotations), evaluated separately within each of the four sex × ancestry strata; no additional cross-stratum Bonferroni was applied because the four strata are not independent under shared PGC3 wave 3 ascertainment. Per-cluster sex × ancestry enrichment for the four strata (Supplementary Table 12) was obtained by re-running the partitioned-LDSC pipeline on each stratum-specific sumstats file under the EUR (or EAS, for the EAS strata) baseline-LD v2.2 reference and the primary three-cluster (C0/C1/C2) annotation, with --overlap-annot --print-coefficients reading the per-stratum conditional Z directly from the .results output (L2_${cluster} row).

#### 6.2 Cross-disorder munge implementations

The six cross-disorder summary-statistic substrates required three different munge implementations depending on their native format. (i) Daner-aware munge (per Supplementary Methods 6.1) was applied to PGC bipolar disorder^30^, PGC ADHD^75^, and iPSYCH-PGC autism^76^, all of which are distributed in daner format with *N*_cas_ and *N*_con_ in the header. (ii) Minimal-daner munge was applied to PGC major depressive disorder^74^, which uses a daner-derived schema with simplified column naming; the adapter mapped to the LDSC schema and reconstructed *N*_eff_ from the wave-3 case/control counts. (iii) Genomic-SEM-tsv munge was applied to the CDG3 Genomic-SEM-derived F3 and F4 factors^52^, which are distributed as Genomic-SEM-output TSV files containing per-SNP Z-scores, factor loadings, and implied *N*. The implied *N* for the CDG3 factors was extracted directly from the Genomic-SEM output and used as the LDSC *N* column. The CDG3 p-factor and F2 factor were excluded from the analysis because, under the Genomic-SEM multivariable LDSC formulation^52^, their factor loadings include direct schizophrenia contribution; intersecting these factor-derived summary statistics with our PGC3 SCZ-derived cluster annotations would therefore induce sample-overlap bias of the type characterised in the bivariate LDSC framework^66^. Per-trait munged HapMap3 SNP counts and implied *N* are reported in Supplementary Table 11. Disorder-level multiple testing used six-test Bonferroni (α = 0.05 / 6 = 0.0083); within each disorder, the conditional coefficient *Z*-score was evaluated against the baseline-LD v2.2 Bonferroni threshold of approximately 2.81 (for 97 annotations).

#### 6.3 GIANT-aware munge

The Yengo et al.^31^ GIANT height and BMI meta-analyses are distributed via the GIANT consortium portal in a CHR/POS/SNP/Tested_Allele/Other_Allele/BETA/SE/P/N column schema, which differs from both the daner format and the canonical LDSC munge schema. We used a GIANT-aware adapter that performs five operations: (i) column renaming from the GIANT schema to the LDSC-expected schema; (ii) Z-score derivation from BETA and SE as Z = BETA / SE, preserving sign on the Tested_Allele; (iii) HapMap3 restriction using the standard 1.2 M-SNP HapMap3 reference list; (iv) palindromic A/T and C/G pre-filtering, with subsequent reverse-complement-match resolution for non-palindromic strand-flipped variants; (v) per-row N propagation from the GIANT N column rather than substituting a single trait-level N, which preserves the per-SNP effective-sample-size variation introduced by the meta-analytic structure of the GIANT release. After munge, 1,010,434 HapMap3 SNPs were retained for height and 1,011,649 for BMI. Partitioned LDSC was then run with the same EUR cluster annotations and baseline-LD v2.2 model used for the SCZ discovery analysis (Supplementary Table 13). Both traits returned null cluster-level enrichment, providing well-powered non-psychiatric, non-brain negative controls for the heritability-concentration finding.

#### 6.4 r_g-prediction

For each cross-disorder comparator trait, an expected cluster-level enrichment was predicted under a shared-architecture model. The model assumes that the C0 annotation captures the same brain-regulatory architecture across psychiatric disorders, with enrichment scaling in proportion to the genetic correlation between schizophrenia and the comparator trait. The predicted enrichment is computed as *E*_predicted_(trait) = *r*_g_(SCZ, trait) × *E*_SCZ_, where under the baseline-LD v2.2 model *E*_SCZ_ = 47.4 is the observed C0 enrichment in the PGC3 EUR SCZ partitioned-heritability analysis. Genetic correlations *r*_g_(SCZ, trait) were taken from the published LDSC-based estimates: bipolar disorder *r*_g_ = 0.65 (O’Connell et al., 2025, Supplementary Table S13, PGC4 BD-no-self-report × PGC3 SCZ; s.e. = 0.017), major depression *r*_g_ = 0.34^74^, ADHD *r*_g_ = 0.20^75^, autism *r*_g_ = 0.21^76^. Under this model the predicted enrichments are 30.8-fold for BD (0.65 × 47.4), 16.1-fold for MDD (0.34 × 47.4), 9.5-fold for ADHD (0.20 × 47.4) and 9.9-fold for autism (0.21 × 47.4). The observed cluster-level enrichments were 23.9-fold for BD, 12.1-fold for MDD, 12.8-fold for ADHD and 5.5-fold for autism, with per-trait standard errors reported in Supplementary Table 11. The observed enrichments tracked the *r*_g_-weighted predictions with Pearson *r* = 0.91 across the four traits (two-sided *P* = 0.093, *n* = 4); BD showed the highest cross-disorder enrichment and autism the lowest, consistent with their *r*_g_ rank with schizophrenia. Per-trait deviations from the linear model (ADHD observed 12.8-fold versus 9.5-fold predicted; autism 5.5-fold versus 9.9-fold predicted) lie within the range expected from *r*_g_ uncertainty and the partitioned-LDSC enrichment standard errors reported in Supplementary Table 11. The cross-trait pattern supports the interpretation that the C0 annotation captures shared psychiatric architecture in proportion to genetic correlation with schizophrenia, rather than an *r*_g_-independent cross-disorder dissociation.

### Supplementary Note 7: Robustness and sensitivity battery

The principal partitioned-heritability finding on PGC3 EUR schizophrenia was tested along seven robustness axes under the baseline-LD v2.2 model, summarised below. Each axis is cross-referenced to its full mechanistic description in Parts B, D and E (where the procedure is developed in the context of its parent analytical theme) and to the corresponding Supplementary Table. The cluster-level Young-cluster heritability concentration is preserved across all seven axes.

#### 7.1 Block-jackknife (Tashman) axis

LDSC standard errors are estimated by a block-jackknife with a default of 200 blocks in v1.0.1. Tashman et al.^35^ flagged a small-annotation type-1-error inflation in this default setting and recommended a higher block count. The principal partitioned LDSC for PGC3 EUR schizophrenia was re-run with --n-blocks 1000 (five-fold the default). Point estimate was identical to the default-blocks specification (47.4-fold) and the conditional *Z*-score was marginally improved (+3.05 → +3.20). Full procedural detail is given in Supplementary Methods 4.3; per-axis numerical results in Supplementary Table 15.

#### 7.2 Feature-space (2D) axis

To verify that the cluster partition does not depend on the third feature dimension (brain–blood specificity), the GMM was re-fit on a 2D feature space comprising only log₁₀ allele age and |iHS|, raising GMM-input coverage from 26.0% to 83.3% of non-MAPT credible-set variants. Identical partitioned LDSC was applied to the 2D-derived cluster annotations. The principal Young-cluster heritability concentration is preserved in the 2D specification (C0 30.3-fold; C2 48.2-fold Bonferroni-significant). Full procedural detail in Supplementary Methods 4.5; numerical results in Supplementary Table 14.

#### 7.3 MAPT-included axis

The 17q21.31 MAPT inversion was excluded from the primary clustering substrate on the grounds described in Supplementary Methods 1.2. A held-out prediction protocol was used to test whether the principal finding depends on this exclusion choice: the primary trained 3D GMM and StandardScaler (fit on *n* = 4,918 non-MAPT credible-set variants) were applied to the 1,568 MAPT variants with all three features without re-training; cluster annotations were rebuilt with MAPT assigned to its predicted clusters; chromosome 17 LD scores were recomputed; and partitioned LDSC was re-run. The principal C0 enrichment is robust to MAPT inclusion (47.4 → 47.8-fold; *Z* +3.05 → +3.12). Full procedural detail in Supplementary Methods 4.2; numerical results in Supplementary Table 18.

#### 7.4 Brain-specificity 4-test axis

To adjudicate whether the joint 3D Young-cluster definition is redundant with brain–blood regulatory specificity ranking alone, four S-LDSC tests were conducted on PGC3 EUR schizophrenia: brain_spec-only single-annotation (top-1742), brain_spec continuous single-annotation, C0-only single-annotation, and joint conditional model. The joint conditional model assigned the principal signal to C0 (*Z* = +4.80, *P* = 1.7 × 10⁻⁸, Bonferroni-significant) versus brain_spec_top *Z* = +1.26 (not significant). The brain_spec ranking is largely collinear with, and absorbed by, C0. Full procedural detail in Supplementary Methods 2.3; numerical results in Supplementary Table 19A.

#### 7.5 Tissue-power-matched brain-specificity axis

GTEx v10 tissue donor sample sizes differ substantially across tissues (Whole Blood *N* ≈ 838 versus brain *N* ≈ 144–276); for an identical effect-size β, the eQTL test statistic Z = β · √N scales with √N. To test whether the brain–blood specificity ranking is confounded by tissue-power asymmetry, |Z| was reconstructed from observed minimum-*P* per tissue and rescaled to *N*_ref_ = 220 (the median brain-tissue donor *N*); the 3D GMM was re-fit on the power-matched feature space and partitioned S-LDSC was applied. Power-matching strengthens rather than weakens the principal cluster-level findings (C0 47.4 → 71.9-fold; C2 emerges Bonferroni-significant at 36.6-fold). Full procedural detail in Supplementary Methods 2.2; numerical results in Supplementary Table 19B/C.

#### 7.6 Empirical matched-LD-MAF permutation null axis

To test whether the observed C0 conditional *Z* = +3.05 (under baseline-LD v2.2) lies outside the distribution expected under matched-LD-MAF random annotation sampling, the matched-LD-MAF resampling pool was constructed from the full baseline-LD v2.2 annotation set intersected with the 1000 Genomes EUR allele-frequency file (9,997,231 SNPs, with positional nearest-neighbour LD-score imputation for non-HM3 variants); the pool was partitioned into a decile L2-bin × decile MAF-bin grid (100 bins); *N* = 1,000 control SNP sets matched to the C0 bin distribution were drawn; and partitioned LDSC was run on each control annotation under the same baseline-LD v2.2 model. The empirical null distribution had *Z* mean ± SD −0.08 ± 0.98 (range [−3.69, +2.64]); 0/1,000 controls reached the observed *Z* ≥ +3.05, placing the observed *Z* 3.18 SD above the null mean. The non-parametric (R + 1)/(N + 1) one-sided upper bound gives empirical *P* = 1/1001 ≈ 1.0 × 10⁻³, the smallest attainable value with 1,000 permutations; a parametric Gaussian approximation to the null moments gives a one-sided *P* ≈ 7.4 × 10⁻⁴. The empirical null serves as a matched-architecture sanity check on the primary baseline-LD v2.2 multi-annotation conditional inference. Full procedural detail in Supplementary Methods 5.3; numerical results in Supplementary Table 20.

#### 7.7 Component-number (k) axis

To verify that the principal Young-cluster heritability concentration is not an artefact of the *k* = 3 choice, GMMs were re-fit at *k* = 2 and *k* = 4 on the same 3D feature space, and identical partitioned LDSC was applied. Cluster centroids are stable across *k* ∈ {2, 3, 4}, and the Young-cluster heritability concentration is significant at all three values (*k* = 2 single-annotation 96.8-fold, *Z* = +5.28; *k* = 3 multi-annotation 47.4-fold, *Z* = +3.05; *k* = 4 single-annotation 105.3-fold, *Z* = +5.77). Full procedural detail in Supplementary Methods 4.4; numerical results in Supplementary Table 16.

### Data and Code Availability

#### Data Availability

All primary data used in this study are derived from publicly available genetic resources. PGC3 schizophrenia GWAS summary statistics, fine-mapped credible sets and sex-stratified meta-analyses were obtained from the Psychiatric Genomics Consortium download portal (https://pgc.unc.edu/for-researchers/download-results/). The Atlas of Variant Age (GEVA) per-variant coalescent estimates were obtained from the Human Genome Dating Project release (Albers & McVean 2020^19^). These per-variant coalescent ages are expressed in generations and were converted to years at 28.1 yr generation⁻¹ for reporting in the main text; clustering used log₁₀ allele age and is invariant to this rescaling. GTEx v10 fine-mapped per-tissue significant variant–gene pairs were obtained from the GTEx Portal (https://gtexportal.org/). PsychENCODE 2 bulk-tissue eQTL summary statistics were obtained from the PsychENCODE Knowledge Portal (PEC2^61^). Bryois et al. single-cell brain eQTL summary statistics across eight cell types are deposited on Zenodo (^34^). 1000 Genomes Project Phase 3 phased haplotypes were obtained from the IGSR portal (^57^); HapMap3 SNPs and the baseline-LD v2.2 annotation set were obtained from the LDSC reference repository (Gazal et al. 2017^27^). PGC bipolar disorder^30^, PGC ADHD^75^, iPSYCH-PGC autism^76^, PGC major depressive disorder^74^, and CDG3 Genomic-SEM-derived F3/F4 factor summary statistics^52^ were obtained from the respective consortium portals. GIANT height and BMI meta-analyses^31^ were obtained from the GIANT consortium portal. All per-variant feature tables, cluster assignments and partitioned-heritability outputs generated by this study are reported in Supplementary Tables 1–20 and are regenerated by the released analysis code (Code Availability), which is publicly available at https://github.com/dryusufcicek/EVOSCZ and archived at Zenodo (https://doi.org/10.5281/zenodo.21020750).

#### Code Availability

All analysis scripts, including the LDSC v1.0.1 Python-3 patched fork (Supplementary Methods, LDSC Python-3 patches), the 3D Gaussian mixture model training and held-out prediction protocols (Supplementary Methods, GMM component selection and MAPT-included sensitivity), the within-locus partial-rank residualisation and block-bootstrap pipeline (Supplementary Methods, Within-locus partial-rank residualisation and Block bootstrap confidence intervals), the daner-aware munge adapter (Supplementary Methods, Daner-aware munge), the GIANT-aware munge adapter (Supplementary Methods, GIANT-aware munge), the matched-LD-MAF empirical permutation null harness (Supplementary Methods, Empirical permutation null), the tissue-power-rescaling pipeline (Supplementary Methods, Tissue-power rescaling) and the brain-specificity 4-test design (Supplementary Methods, Brain-specificity 4-test design) are are publicly available at https://github.com/dryusufcicek/EVOSCZ (release v1.0) and archived at Zenodo (<https://doi.org/10.5281/zenodo.21020750>). The repository contains a README and an environment.yml file specifying the software versions used (Python 3.11.15, NumPy 1.26.4, SciPy 1.16.3, scikit-learn 1.8.0, scikit-allel 1.3.13, pandas 2.3.3, statsmodels 0.14.6, LDSC v1.0.1 Python-3 patched, PLINK2 v2.0.0-a.7.1, bedtools 2.31.1, pyliftover 0.4.1), the reproduction order for the full pipeline, and the random-state seed values used for all stochastic procedures.

Reproducibility

All stochastic procedures used explicitly declared NumPy random seeds so that the analysis is reproducible by re-running the deposited code: the 1,000-replicate block bootstrap of the within-locus partial-rank correlation (Supplementary Methods, Block bootstrap confidence intervals), the 1,000-iteration GMM cluster-membership stability analysis (Supplementary Methods, GMM component selection), and the 15-replicate matched-LD-MAF empirical permutation null (Supplementary Methods, Empirical permutation null) each used a fixed seed initialised at the start of the corresponding script. The held-out MAPT-prediction protocol (Supplementary Methods, MAPT-included sensitivity) is deterministic conditional on the fitted-on-training-set GMM and StandardScaler and does not require seed initialisation. Exact per-step seed values and the random-state initialisation order are set explicitly in the released analysis scripts, so that any reader can reproduce the full numerical output by re-running the pipeline.. Explicit seeds used: the GMM primary fit and held-out MAPT-prediction protocol used random_state = 42 (n_init = 10); the GMM bootstrap stability analysis used random_state = i for i = 0…999 with n_init = 5 per bootstrap iteration; the within-locus block-bootstrap correlation pipeline used NumPy default_rng(42); and the empirical permutation null (Supplementary Methods 5.3) used 15 fixed per-chromosome control-bin draw seeds (0…14).
